# The Identification of Key Genes and Biological Pathways in Cardiac Arrest by Integrated Bioinformatics and Next Generation Sequencing Data Analysis

**DOI:** 10.1101/2024.08.15.608116

**Authors:** Basavaraj Vastrad, Chanabasayya Vastrad

## Abstract

Cardiac arrest (CA) is a common cause of death world wide. The disease has lacks effective treatment. Efforts have been made to elucidate the molecular pathogenesis of CA, but the molecular mechanisms remain elusive. To identify key genes and pathways in CA, the next generation sequencing (NGS) GSE200117 dataset was downloaded from the Gene Expression Omnibus (GEO) database. DESeq2 tool was used to recognize differentially expressed genes (DEGs). Gene ontology (GO) and REACTOME pathway enrichment analyses were performed to analyze the DEGs and associated signal pathways in the g:Profiler database. The IID database was used to construct the protein-protein interaction (PPI) network, and modules analysis was performed using Cytoscape. A miRNA-hub gene regulatory network and TF-hub gene regulatory network were then constructed to screen miRNAs, TFs and hub genes by miRNet and NetworkAnalyst database and Cityscape software. Receiver operating characteristic curve (ROC) analysis used to verified the hub genes. In total, 844 DEGs were identified, comprising 414 up regulated genes and 430 down regulated genes. GO and REACTOME pathway enrichment analyses indicated that the DEGs for CA were mainly enriched in organonitrogen compound metabolic process, response to stimulus, translation and immune system. Ten hub genes (up-regulated: HSPA8, HOXA1, INCA1 and TP53; down-regulated: HSPB1, LMNA, SNCA, ADAMTSL4 and PDLIM7) were screened. We also predicted miRNAs (hsa-mir-1914-5p and hsa-mir-598-3p) and TFs (JUN and PRRX2) targeting hub genes. This study uses a series of bioinformatics technologies to obtain hug genes, miRNAs and TFs, and key pathways related to CA. These analysis results provide us with new ideas for finding biomarkers and treatment methods for CA.

## Introduction

Cardiac arrest (CA) is the sudden loss of all heart activity characterized by low cardiac output or ventricular systolic or diastolic dysfunction and ultimately leading to death if untreated [1]. CA contributing significantly to mortality rates in both developed and developing countries [2]. CA often causes sudden collapse, no pulse, no breathing and loss of consciousness [3]. The pathogenesis of CA remains poorly understood [4]. Thus, various other factors, including genetic [5], inflammatory [6], neurologic complications [7], hypertension [8], obesity [9], diabetes mellitus [10] and high cholesterol [11], should be considered to better understand the complex pathophysiology of CA. Implantable cardioverter-defibrillator [12], coronary angioplasty [13], coronary artery bypass surgery [14], radiofrequency catheter ablation [15] and corrective heart surgery [16] are the major therapeutic option for CA. Beta blockers [17], Angiotensin-converting enzyme (ACE) inhibitors [18] and calcium channel blockers [19] are alternative treatment options. However, the efficacy of these treatment strategies, whether surgical or drug treatments, is still limited due to the high recurrence rate of the CA. Therefore, the discovery of novel molecular markers of CA is essential for early diagnosis and development of novel drug targets.

The molecular markers and signaling pathways are being used for the etiological diagnosis of CA. Recently, a system-level analysis of whole genome wide expression data identified several genes include GPC5 [20], TGFBR2 [21], KCNA5 [22], CASQ2 [23] and NOS3 [24], and signaling pathways include ATF6 and Nrf2 signaling pathways [25], TNFSF8/AMPK/JNK signaling pathway [26], AdipoR1□AMPK signaling pathway [27], GSK-3β/Nrf2/HO-1 signaling pathway [28] and PI3K/Akt signaling pathway [29] were significantly associated with CA. Next generation sequencing (NGS) is a high-throughput technology used for collecting global gene expression data from recruited samples of various diseases [30–31]. These NGS data are usually deposited and available in free public websites, such as the NCBI-Gene Expression Omnibus database (NCBI-GEO) (https://www.ncbi.nlm.nih.gov/geo) [32]. Integrated bioinformatics analyses of NGS data derived from different investigation of CA could help identify the hub genes and further demonstrate their related functions and potential therapeutic targets in inflammatory CA.

In the current investigation, public NGS data of GSE200117 [33] from NCBI-GEO were downloaded. 106 CA and 46 normal control samples in GSE200117 was available. DEGs between CA and normal control samples were filtered and obtained using the DESeq2 [34]. Gene ontology (GO) and REACTOME pathway enrichment analyses of the DEGs were performed using the g:Profiler. The functions of the DEGs were further assessed by PPI network and modular analyses to identify the hub genes in CA. Using bioinformatics analysis of the miRNA-hub gene regulatory network and TF-hub gene regulatory network, we predicted microRNAs and TFs that might be involved in the CA. The findings were further validated by receiver operating characteristic curve (ROC) analysis. The molecular pathogenesis of CA is still unclear due to its heterogeneity, therefore, our findings would be beneficial for the diagnosis and prognosis of CA.

## Materials and Methods

### Next generation sequencing (NGS) data source

GSE200117 [33] was obtained from NCBI-GEO, a public database of NGS, to filter the DEGs between CA and normal control samples. The microarray profile GSE200117 was based on GPL18573 Illumina NextSeq 500 (Homo sapiens) platform and consisted of 106 CA and 46 normal control samples.

### Identification of DEGs

The DESeq2 in R bioconductor tool was used to analyze the DEGs between CA and normal control samples in the NGS data of GSE200117. The adjusted P-value and [log□FC] were calculated. The Benjamini & Hochberg false discovery rate method was used as a correction factor for the adjusted P-value in DESeq2 [35]. The statistically significant DEGs were identified according to P<0.05, [log□FC] > 0 for up regulated genes and log□FC] < −0.71 for down regulated genes. Volcano plot was constructed by ggplot2 package based on R language. The “gplot” package of R language was used to construct a heatmap of the DEGs.

### GO and pathway enrichment analyses of DEGs

g:Profiler (http://biit.cs.ut.ee/gprofiler/) [36] is an open database that integrates biological data and analytical tools for functional annotation of genes and pathways. Gene Ontology (GO) annotation (http://www.geneontology.org) [37] is a bioinformatics tool for annotating genes and analyzing the biological processes (BP), cell components (CC) and molecular function (MF) they are involved in CA. REACTOME (https://reactome.org/) [38] is a database for analyzing relevant signaling pathways in large scale molecular datasets generated by high-throughput experimental techniques. P value□<□0.05 was defined statistically significant.

### Construction of the PPI network and module analysis

The common DEGs of GSE200117 was analyzed using the online website “Integrated Interactions Database (IID)” (http://iid.ophid.utoronto.ca/search_by_proteins/) [39]. Then, the software Cytoscape (v 3.10.2) (http://www.cytoscape.org/) [40] was used to establish a PPI network. The Network Analyzer in Cytoscape was utilized to calculate node degree [41], betweenness [42], stress [43] and closeness [44] for used to identify the hub genes in the PPI network. Plug-in PEWCC Cytoscape software is an APP was used to perform modular analysis [45]. Finally, GO and pathway enrichment analyses of candidate genes in each module of PPI network were performed using g:Profiler.

### Construction of the miRNA-hub gene regulatory

The miRNAs associated with CA were identified in miRNet (https://www.mirnet.ca/) [46], which is an experimentally validated miRNA-hub gene interaction database. The intersection of miRNAs and hub genes in CA was used to construct the miRNA-hub gene regulated network. Cytoscape software [40] was used to visualize the network.

### Construction of the TF-hub gene regulatory network

The TFs associated with CA were identified in NetworkAnalyst (https://www.networkanalyst.ca/) [47], which is an experimentally validated TF-hub gene interaction database. The intersection of TFs and hub genes in CA was used to construct the TF- hub gene regulated network. Cytoscape software [40] was used to visualize the network.

### Receiver operating characteristic curve (ROC) analysis

The diagnostic accuracy of the hub genes was analyzed by ROC curves. To determine the true positive rate (Sensitivity) and false-positive rate (Specificity) of the identified hub genes using the “pROC” package in R [48]. Area under curve (AUC) was calculated and used to screen the ROC values.

## Results

### Identification of DEGs

There are 844 DEGs screened from GSE200117 between CA group with normal control group which consists of 414 up-regulated and 430 down-regulated genes (Table 1). Significant up-regulated and down-regulated genes were shown in Volcano plot (Fig.1). The DEGs with significant difference between the CA groups with normal control groups were shown in the heatmap (Fig.2).

**Table 1.**
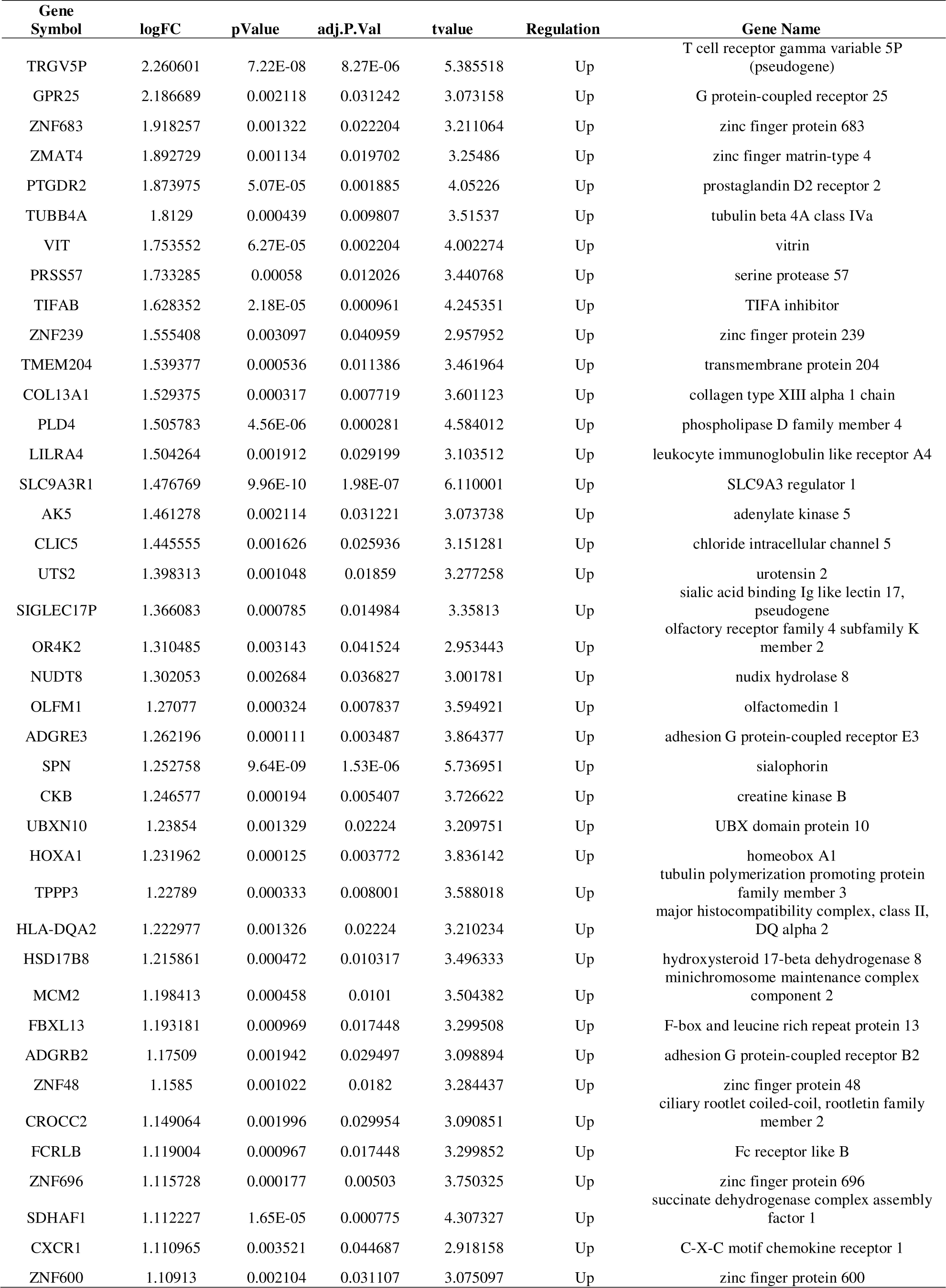
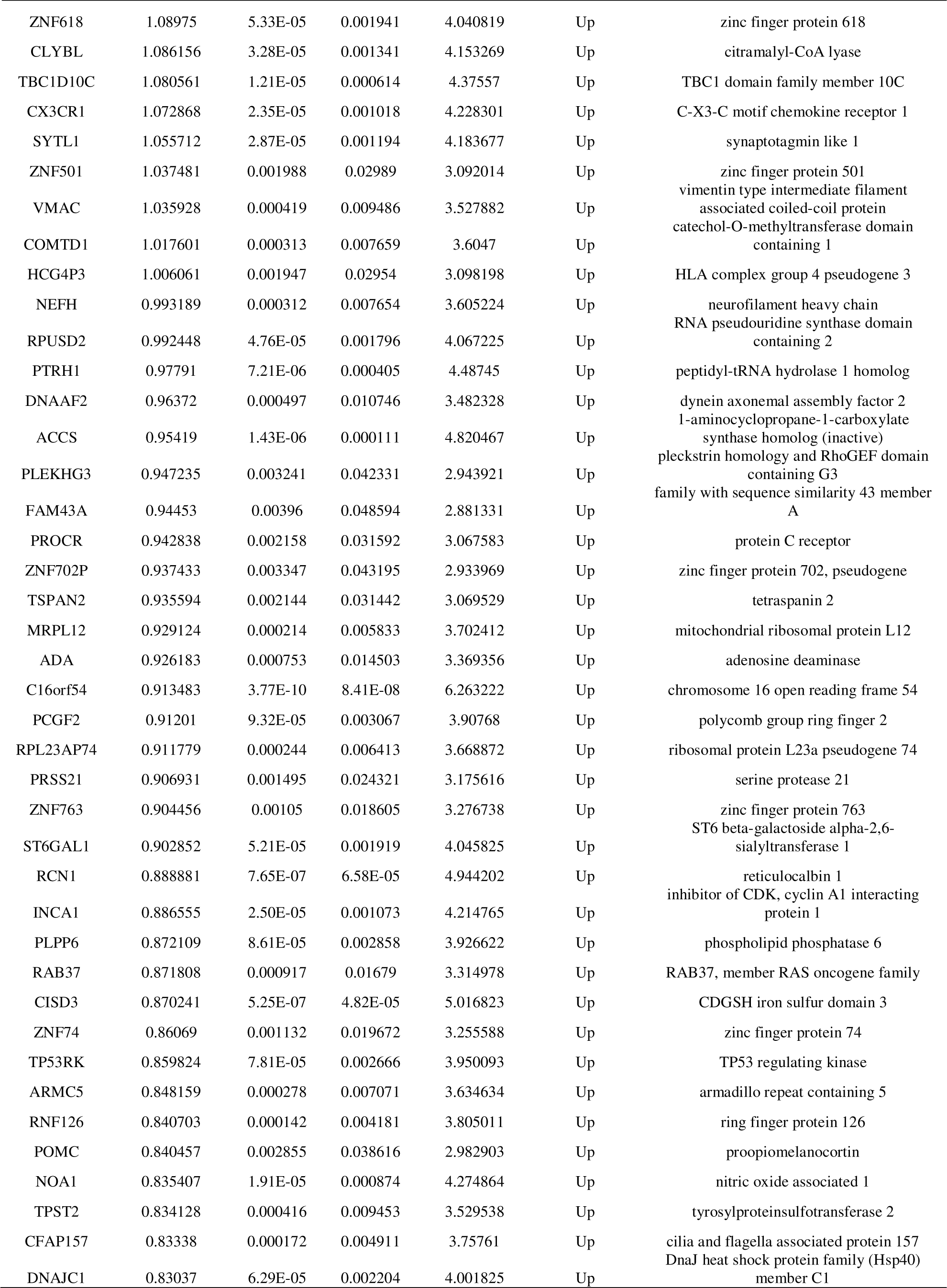
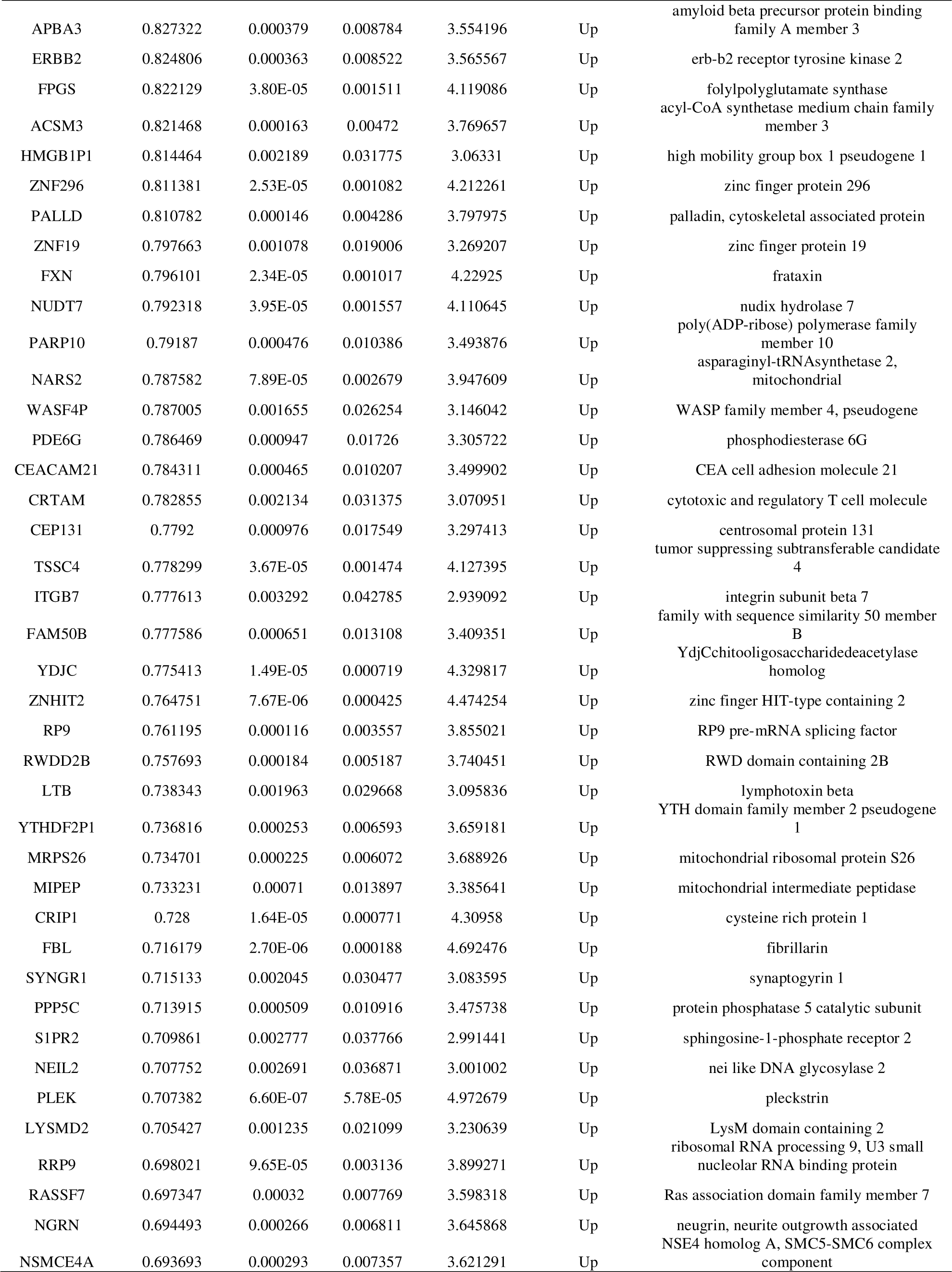
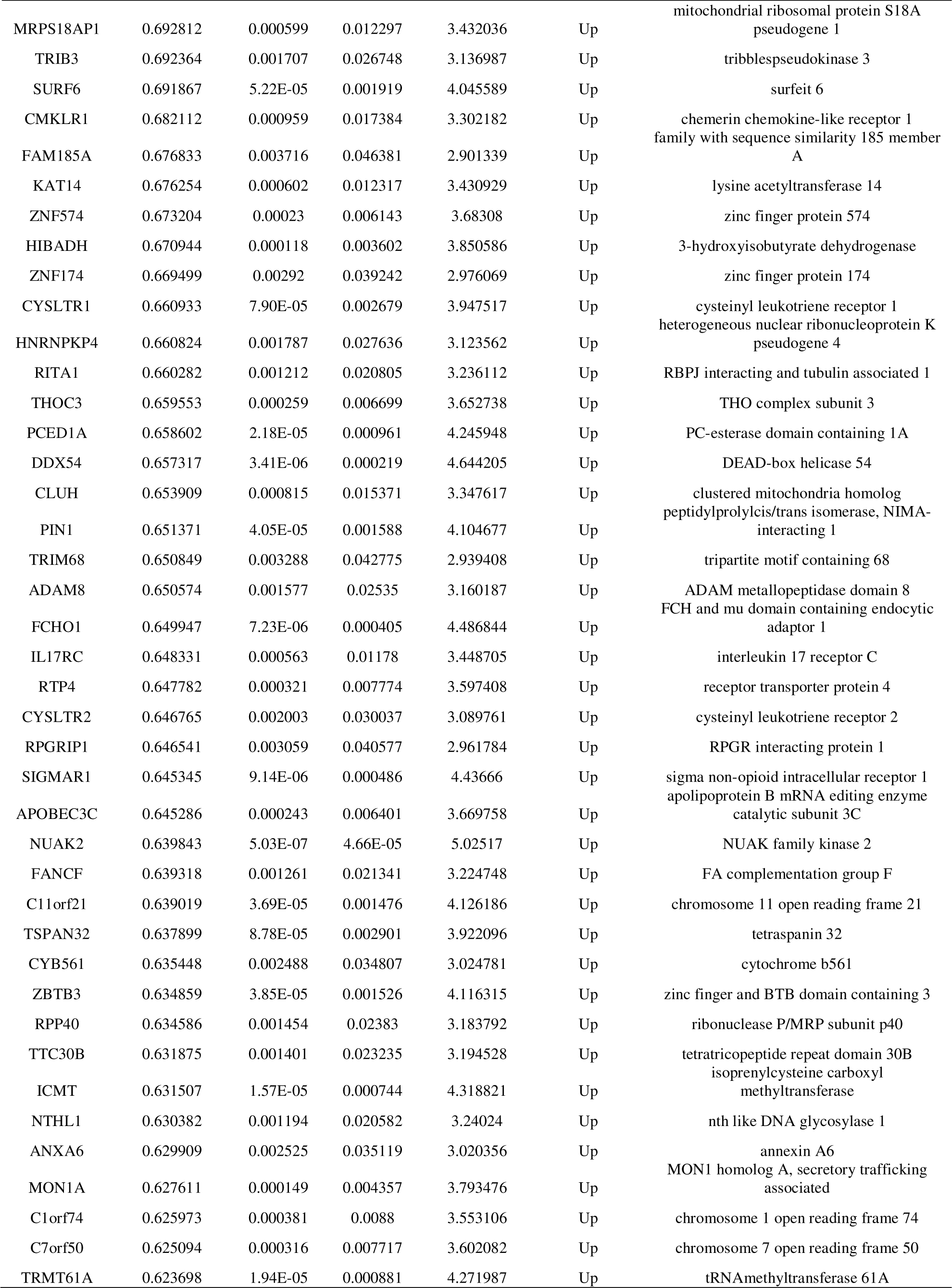
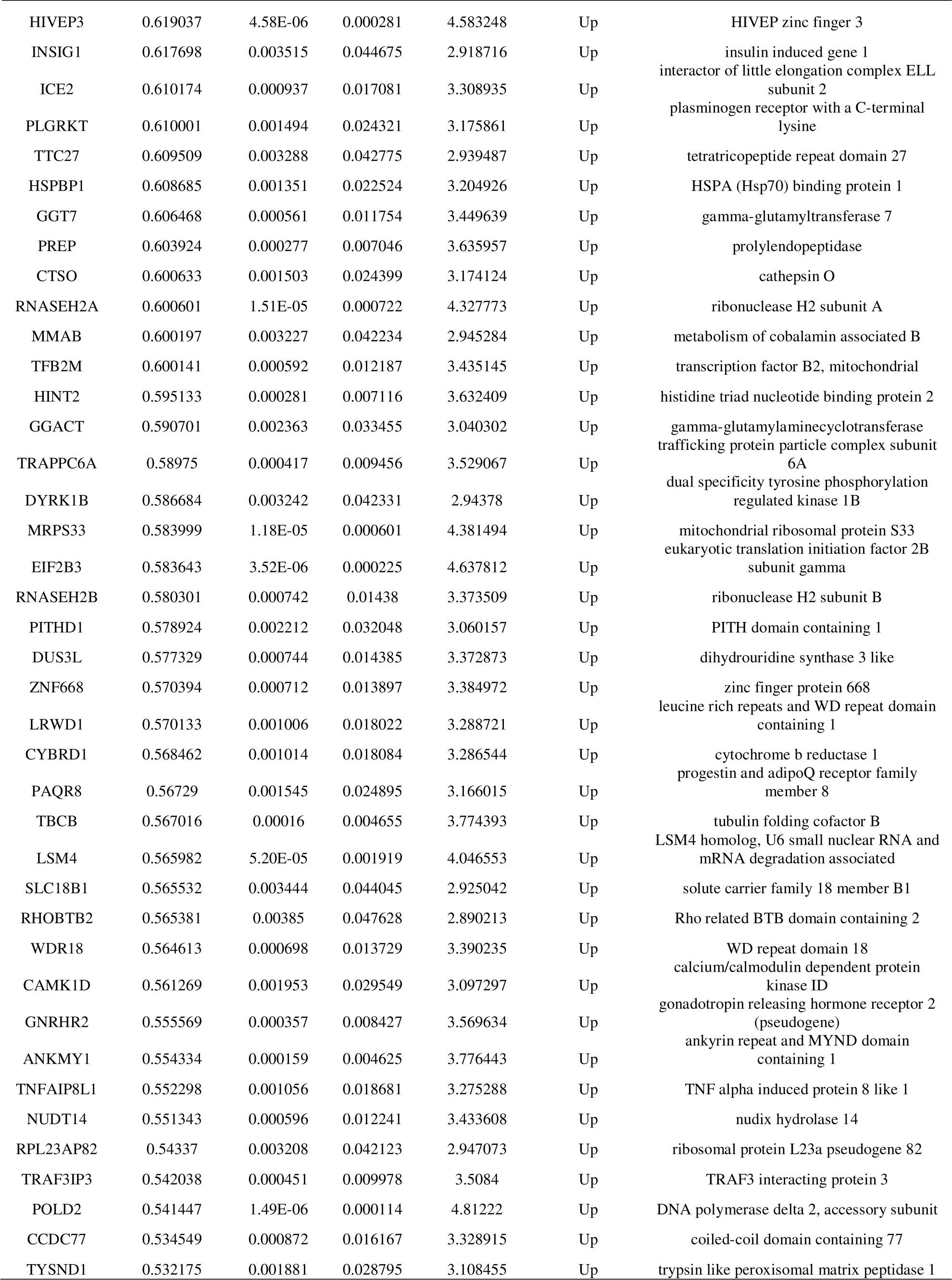
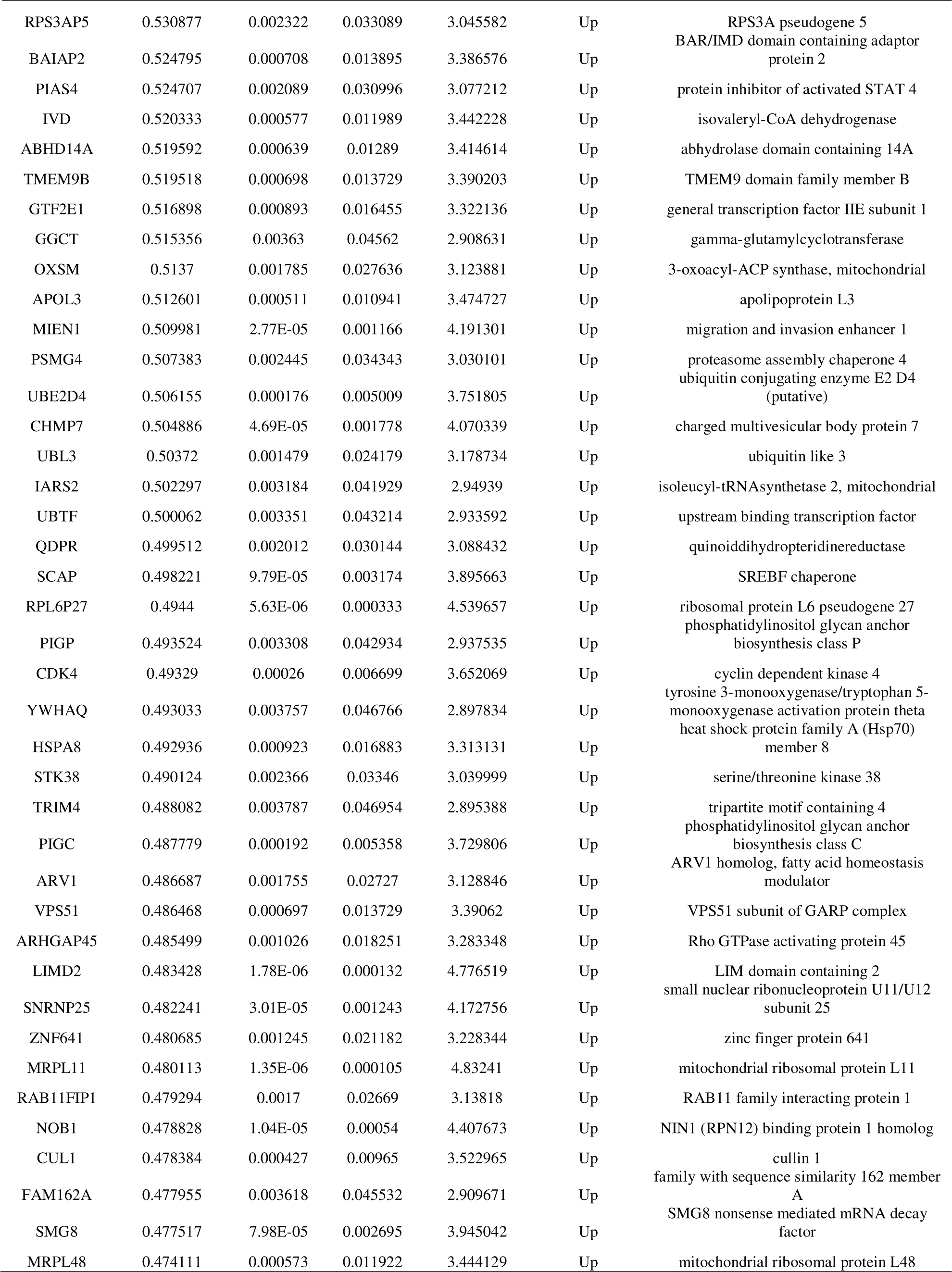
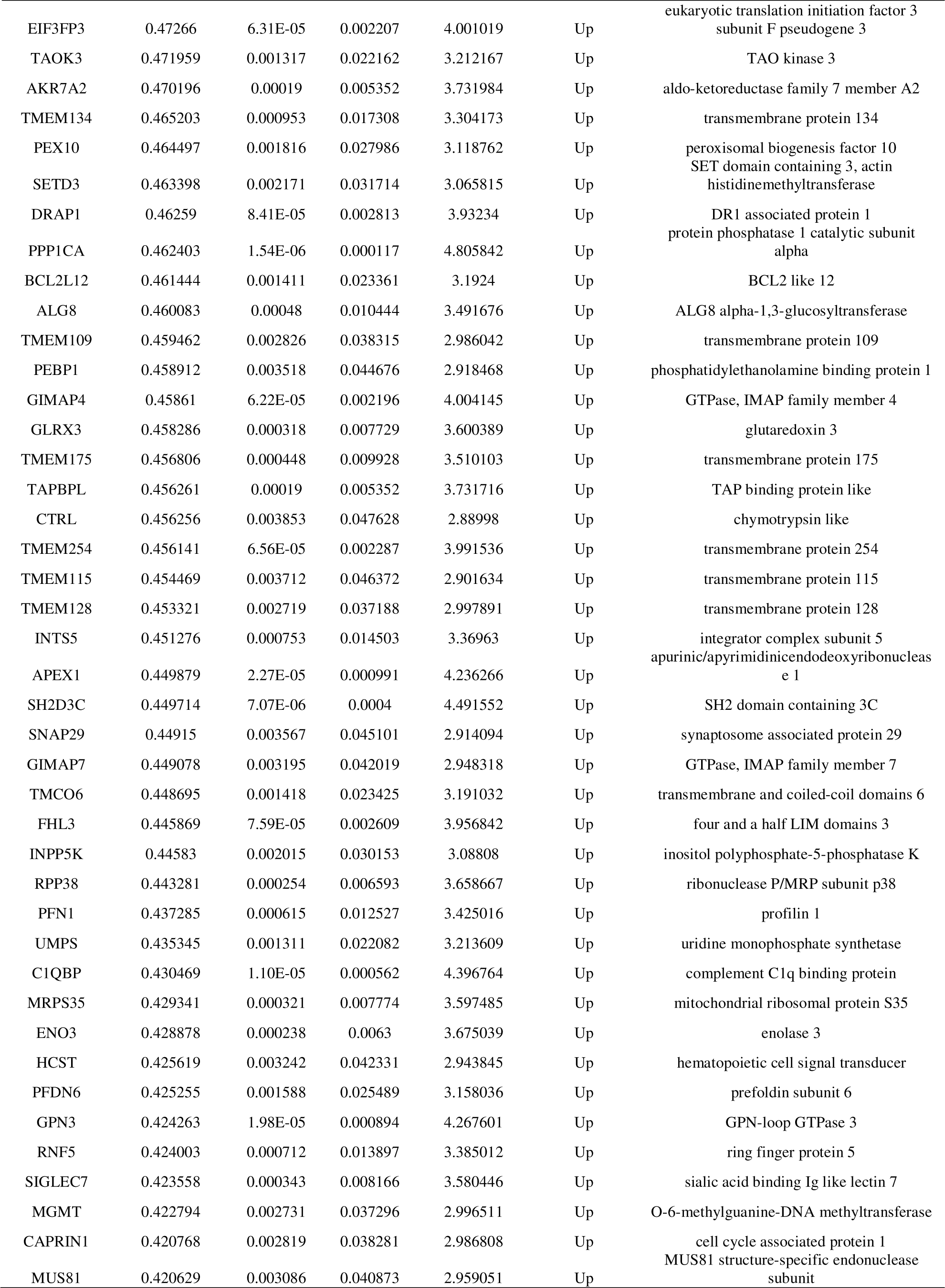
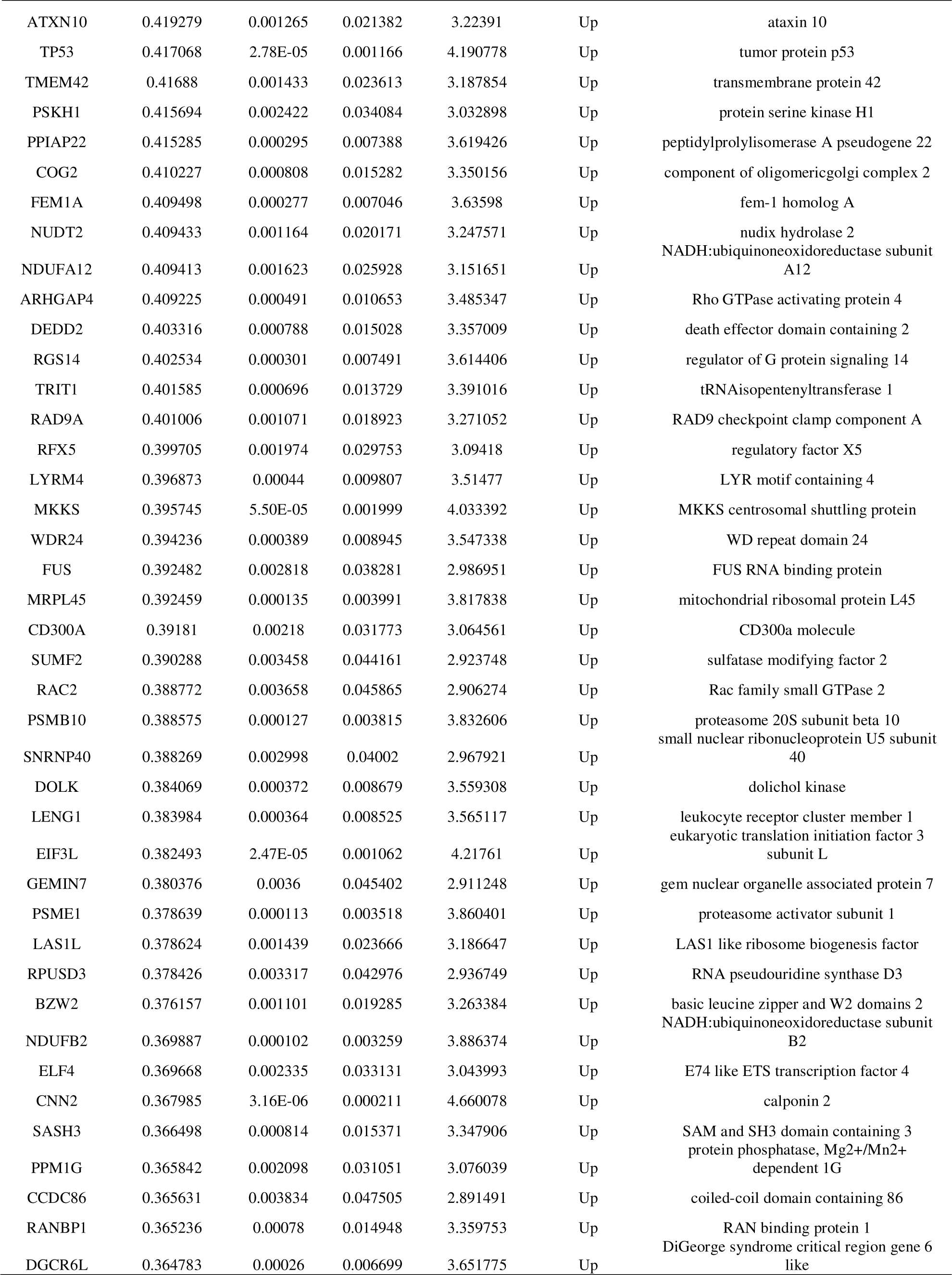
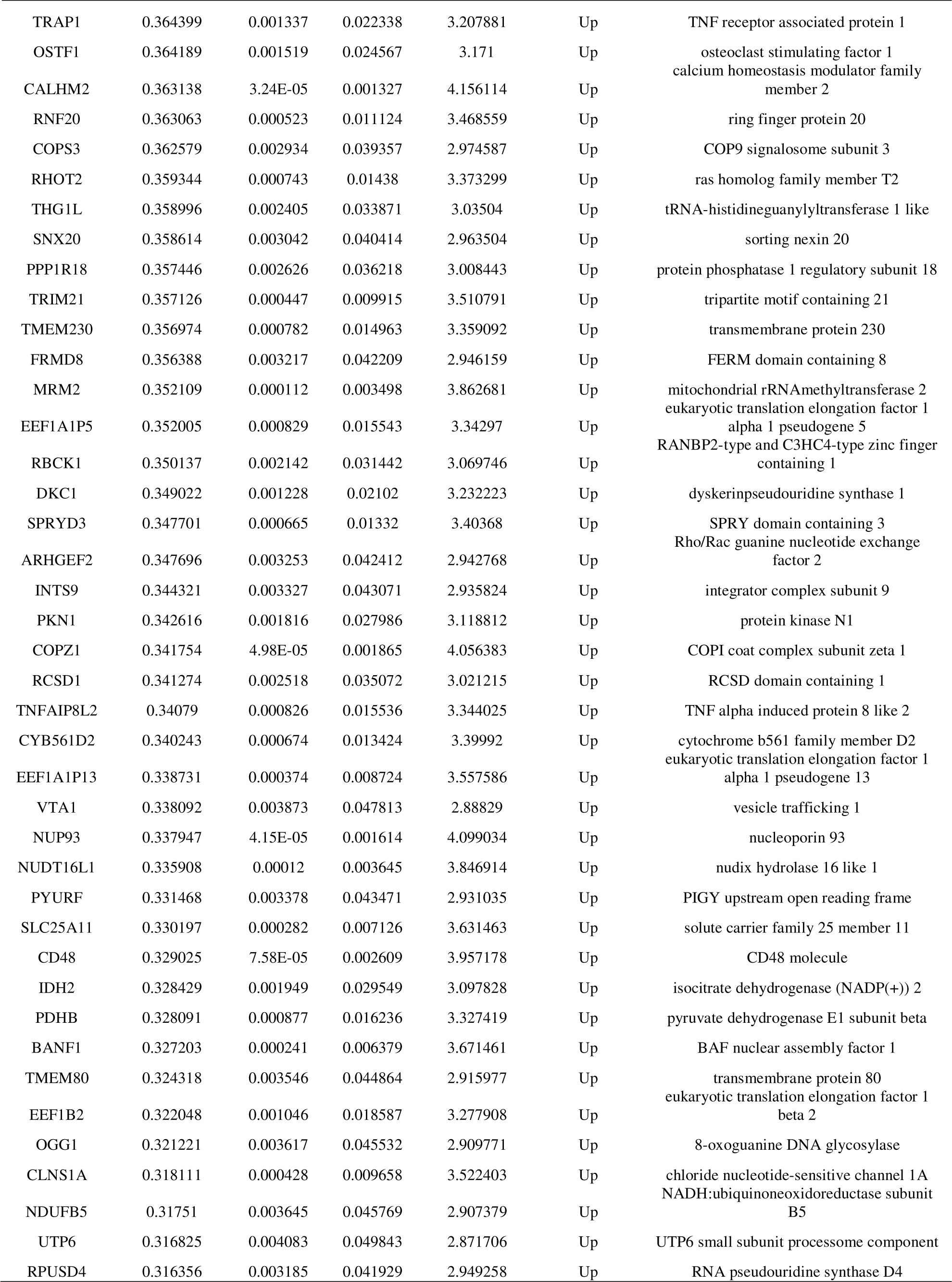
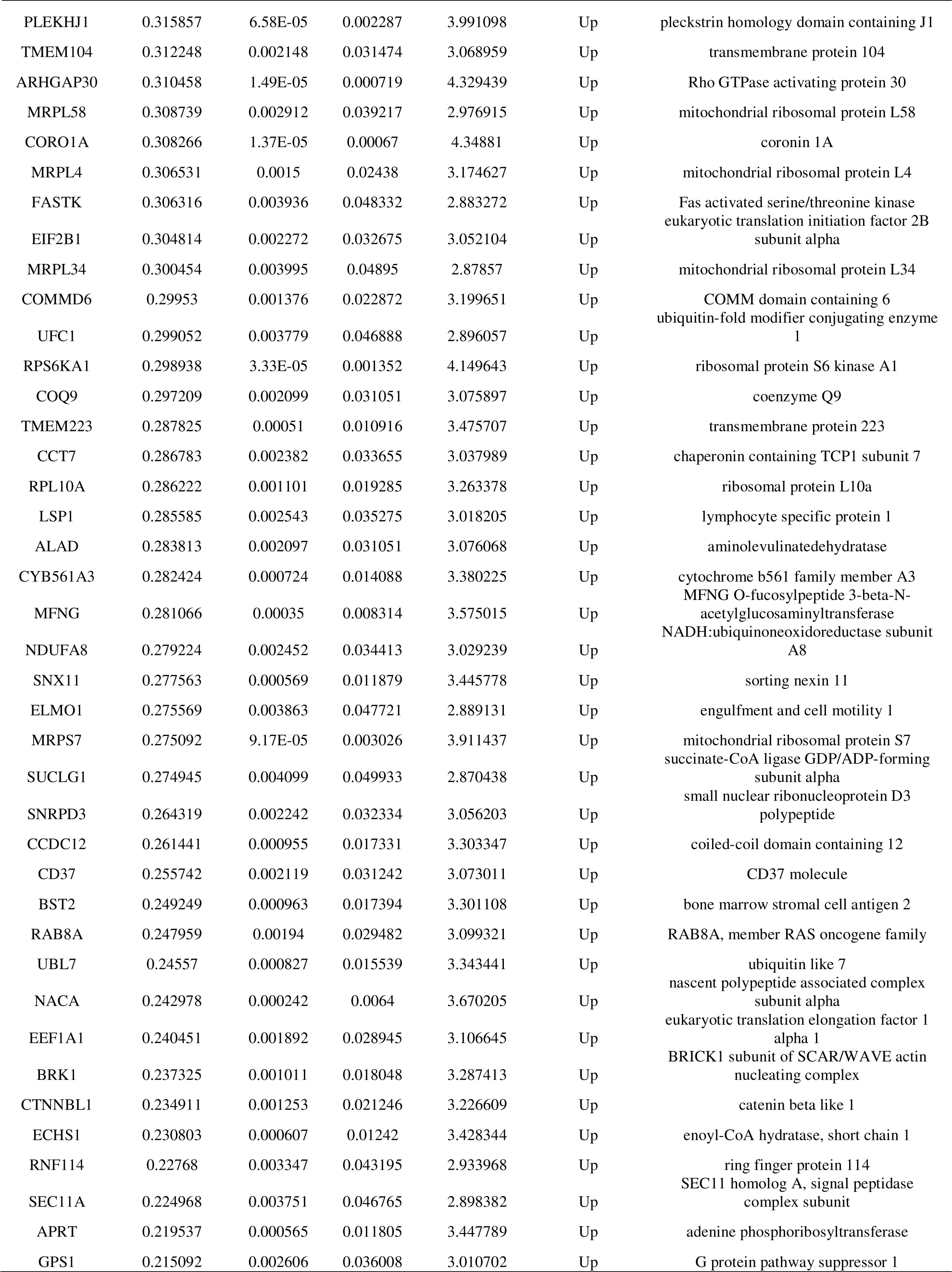
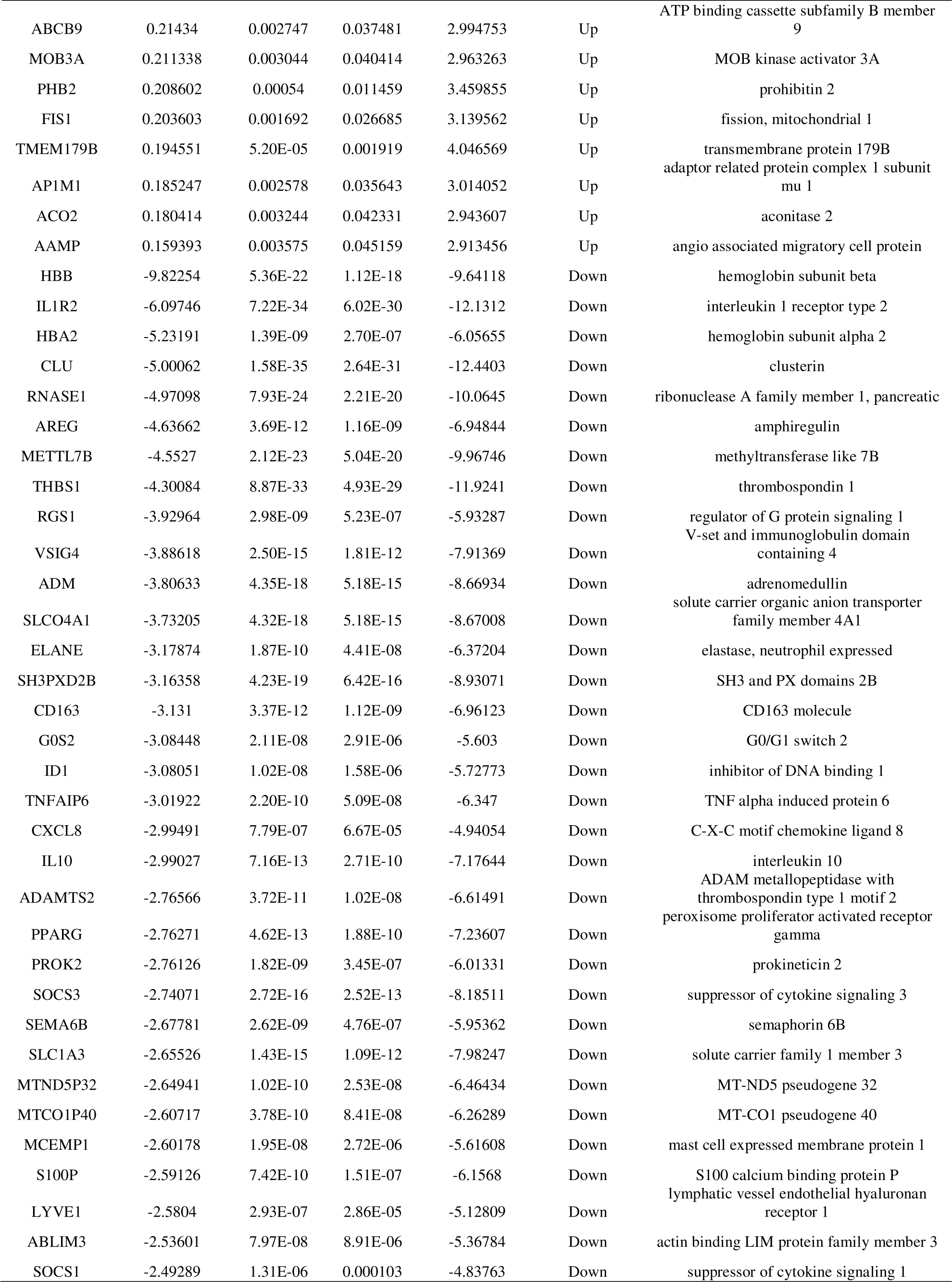
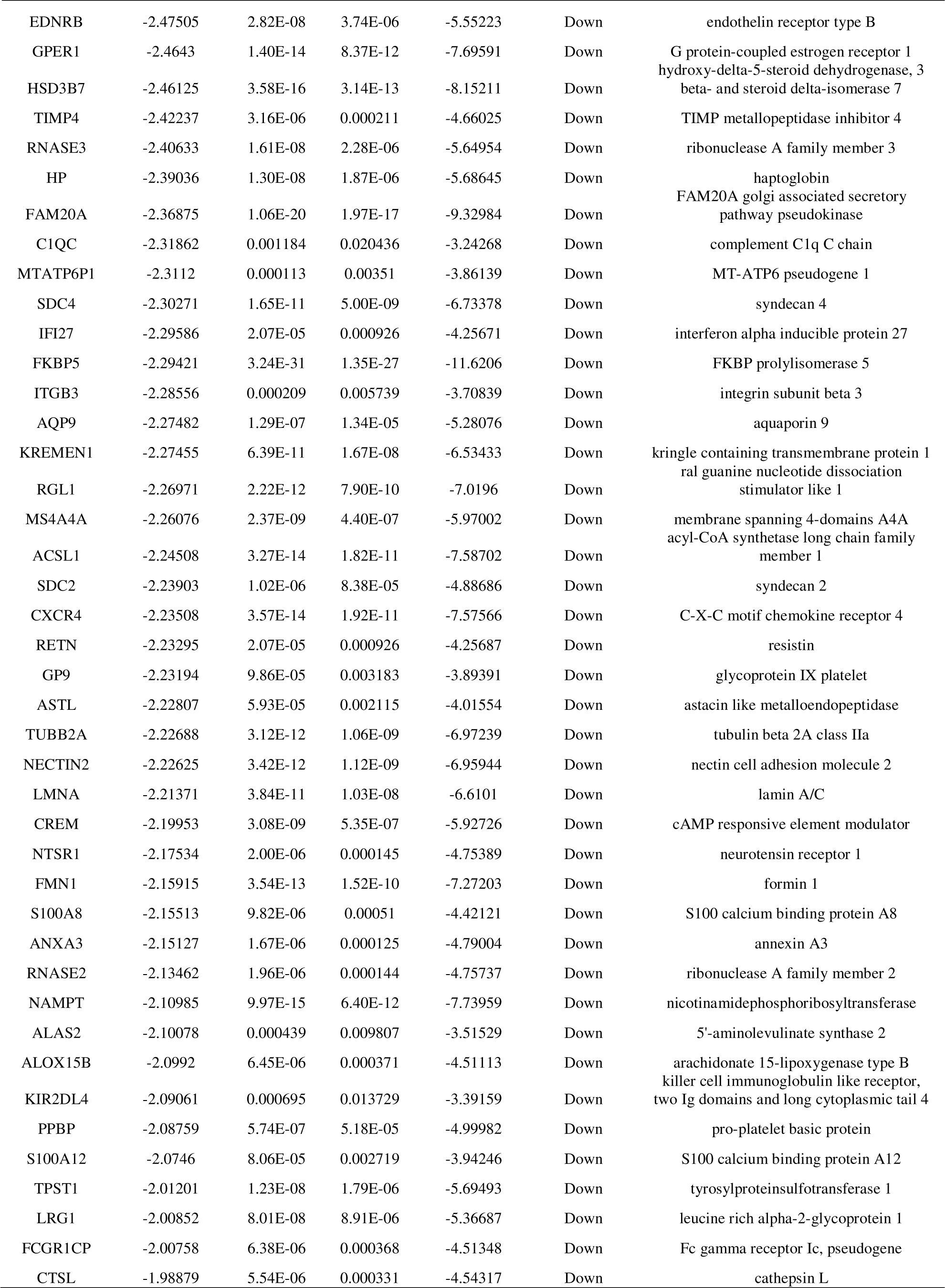
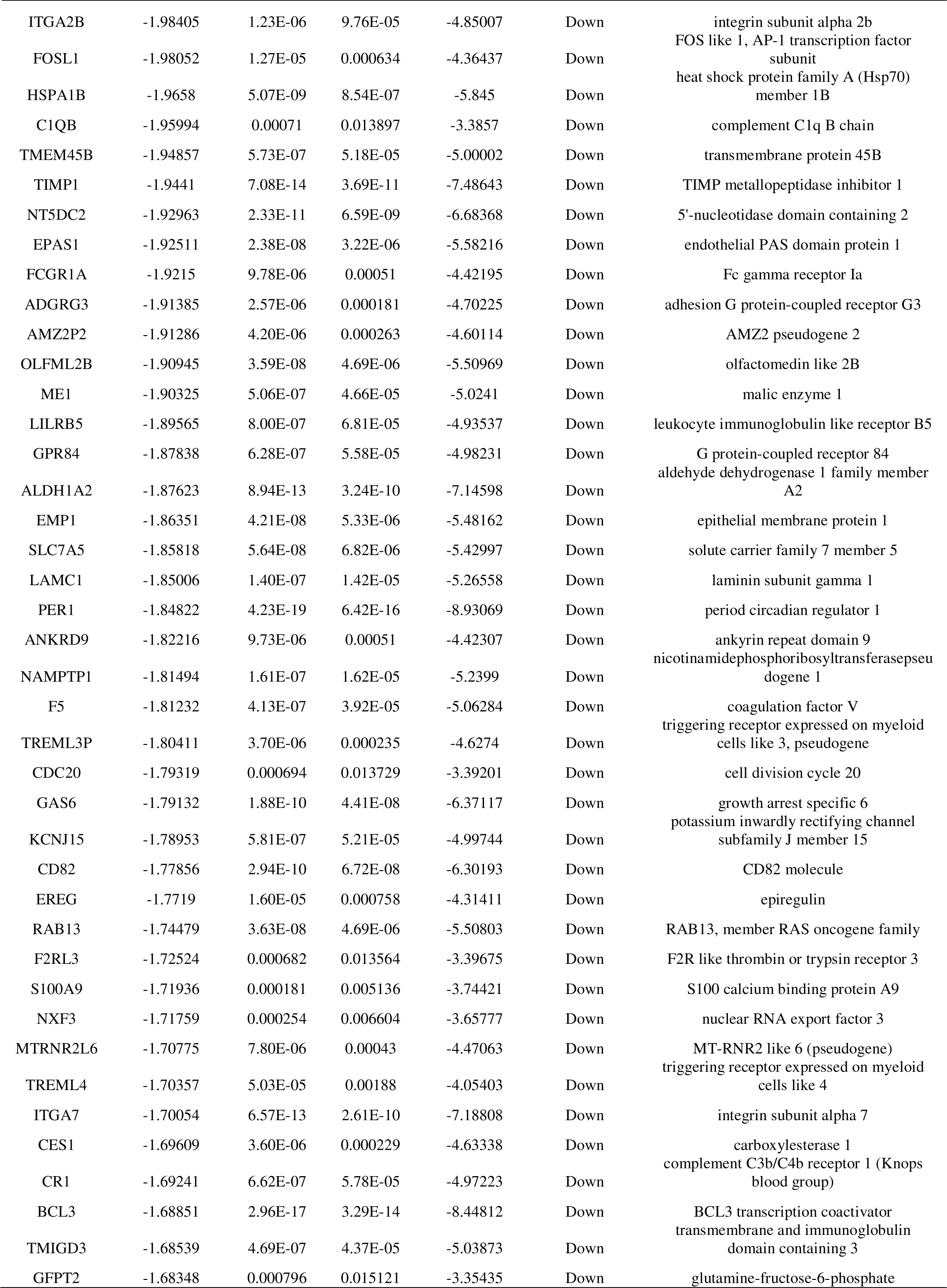
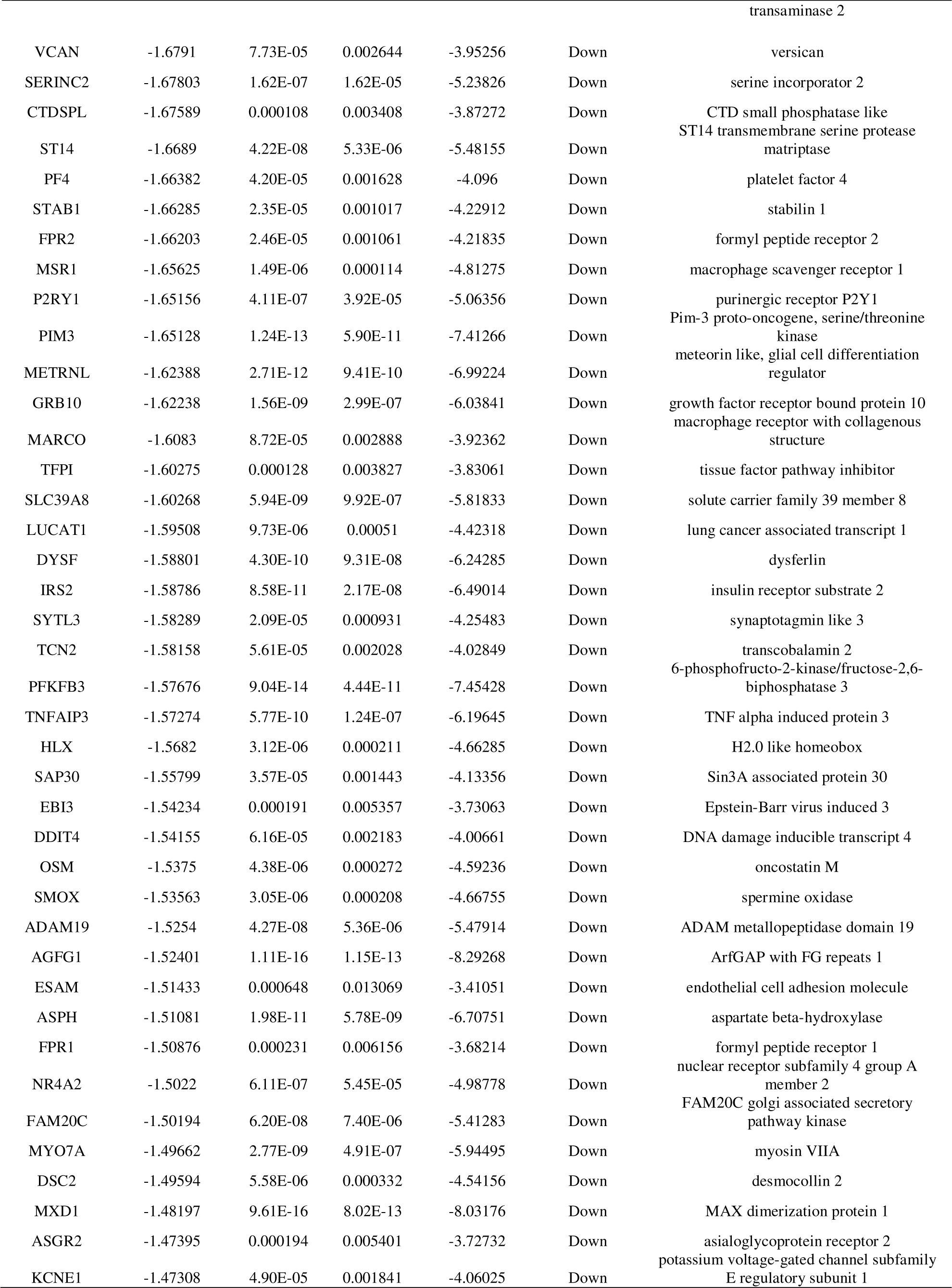
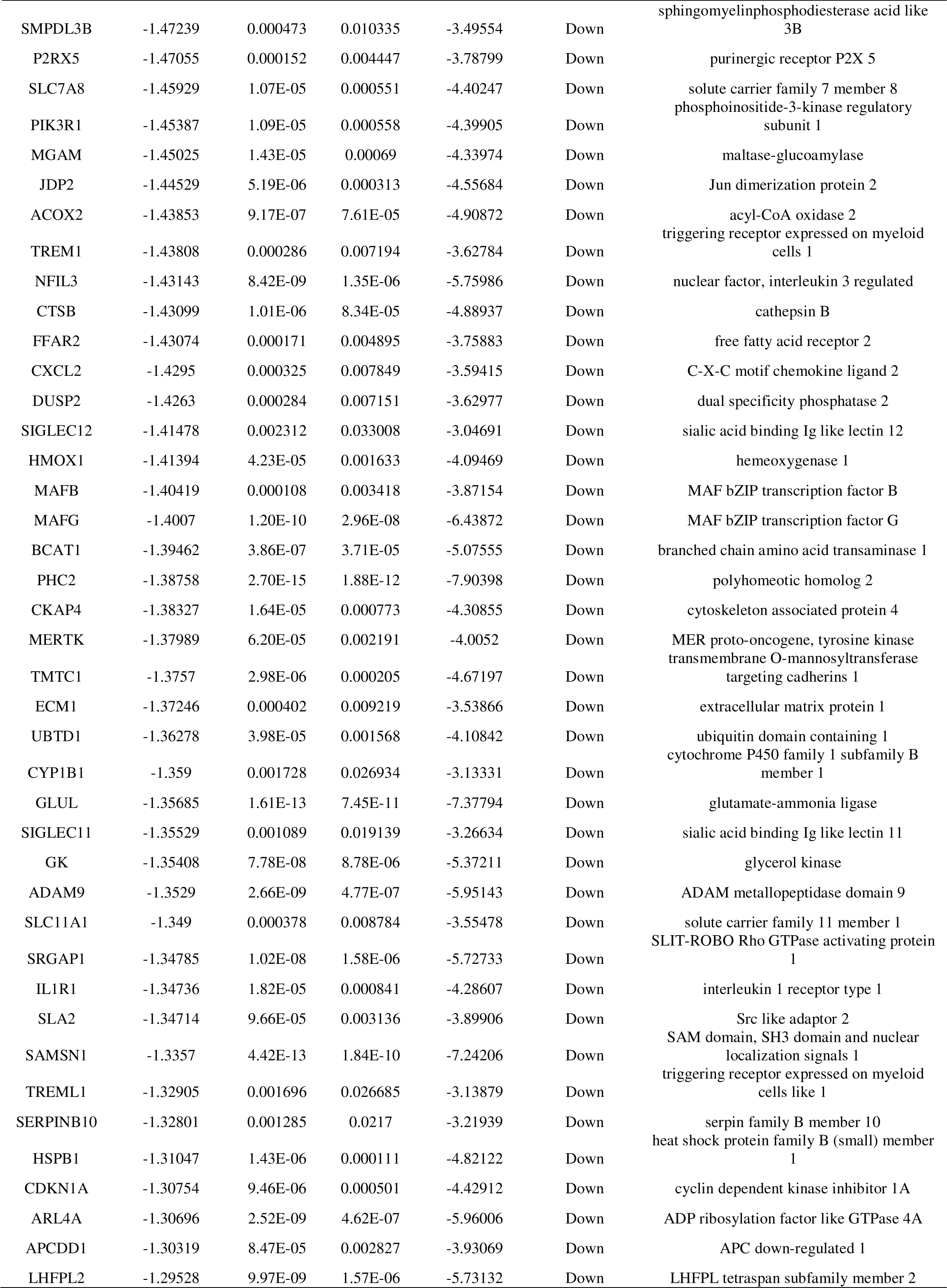
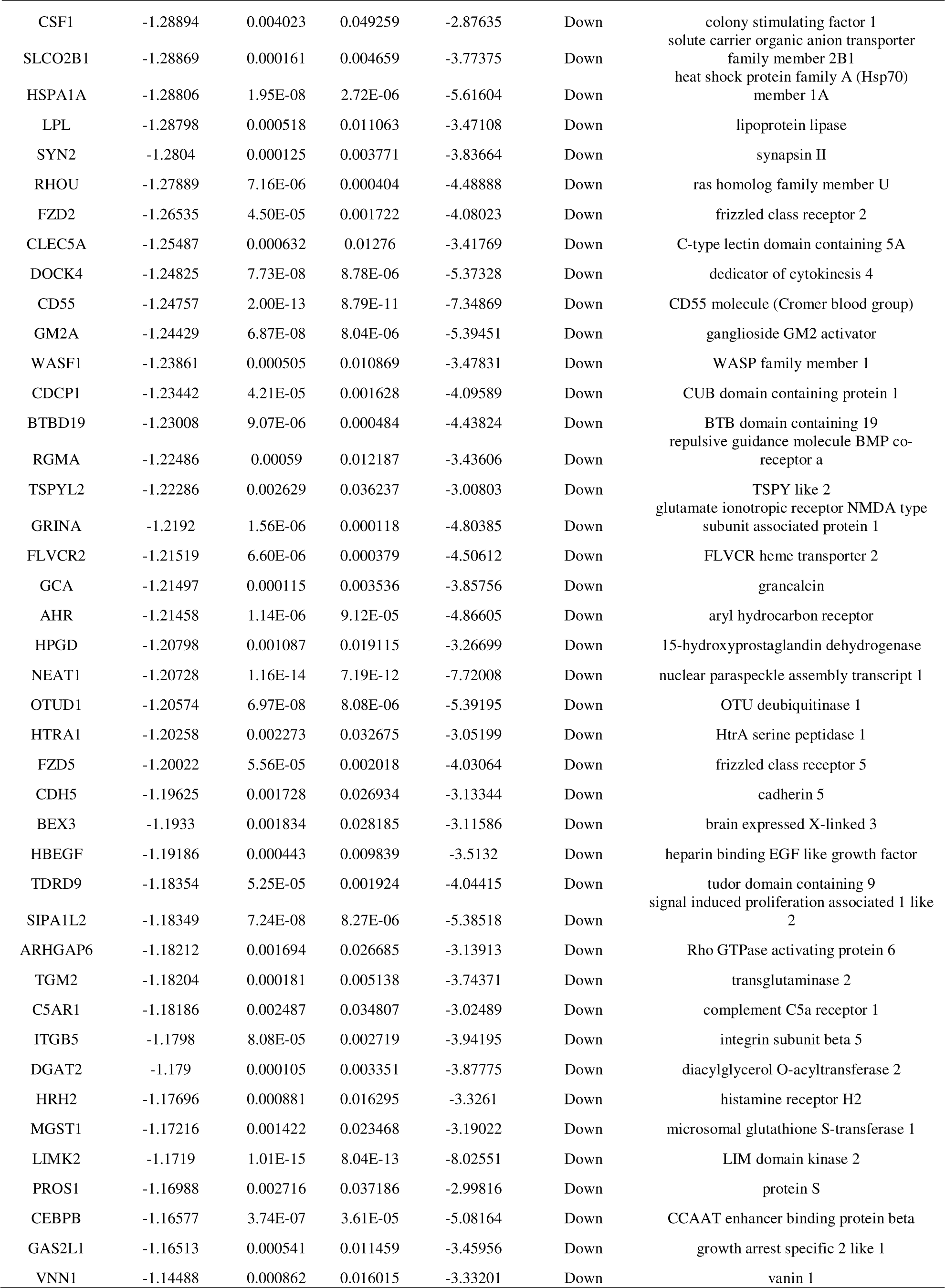
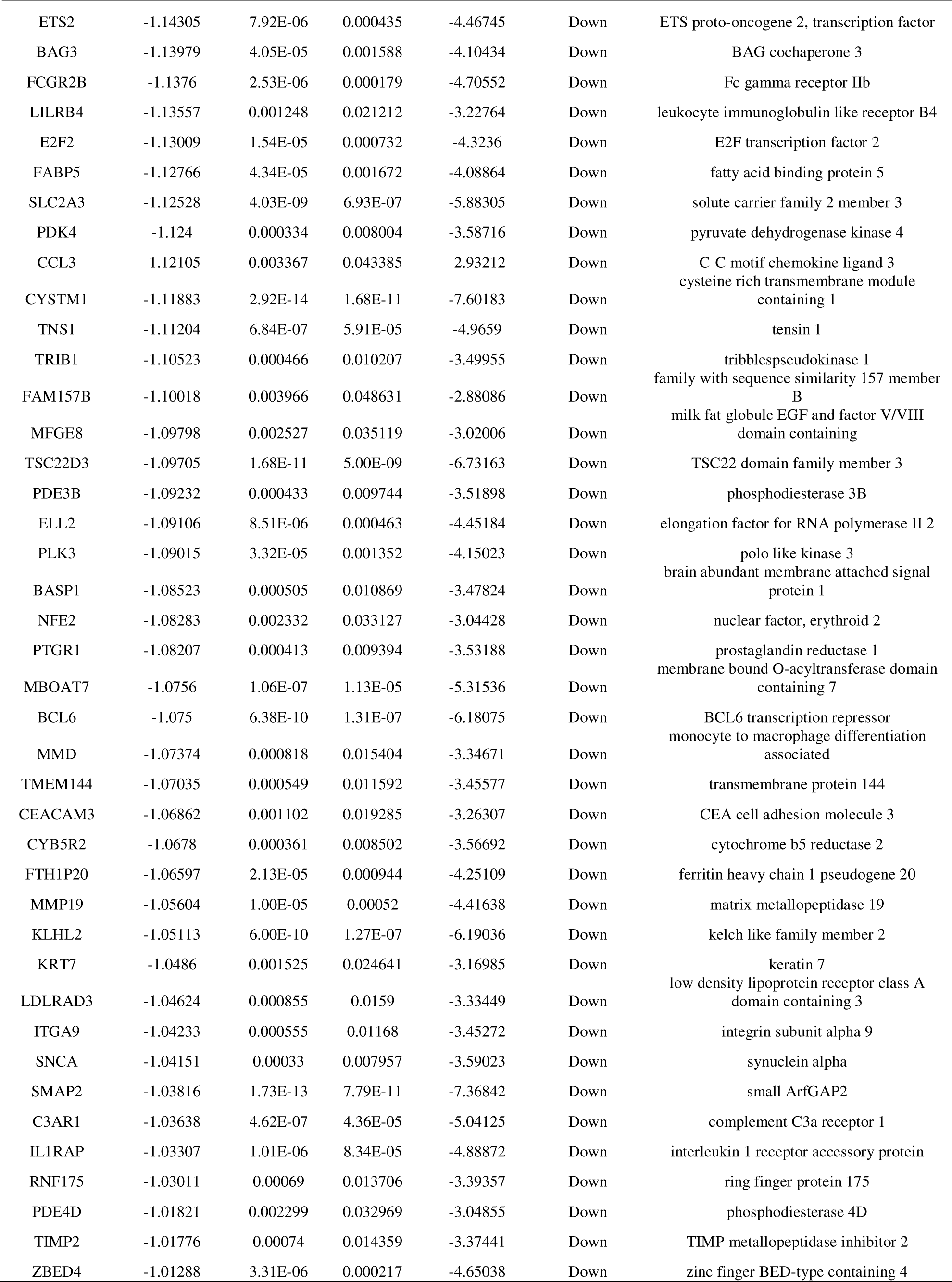
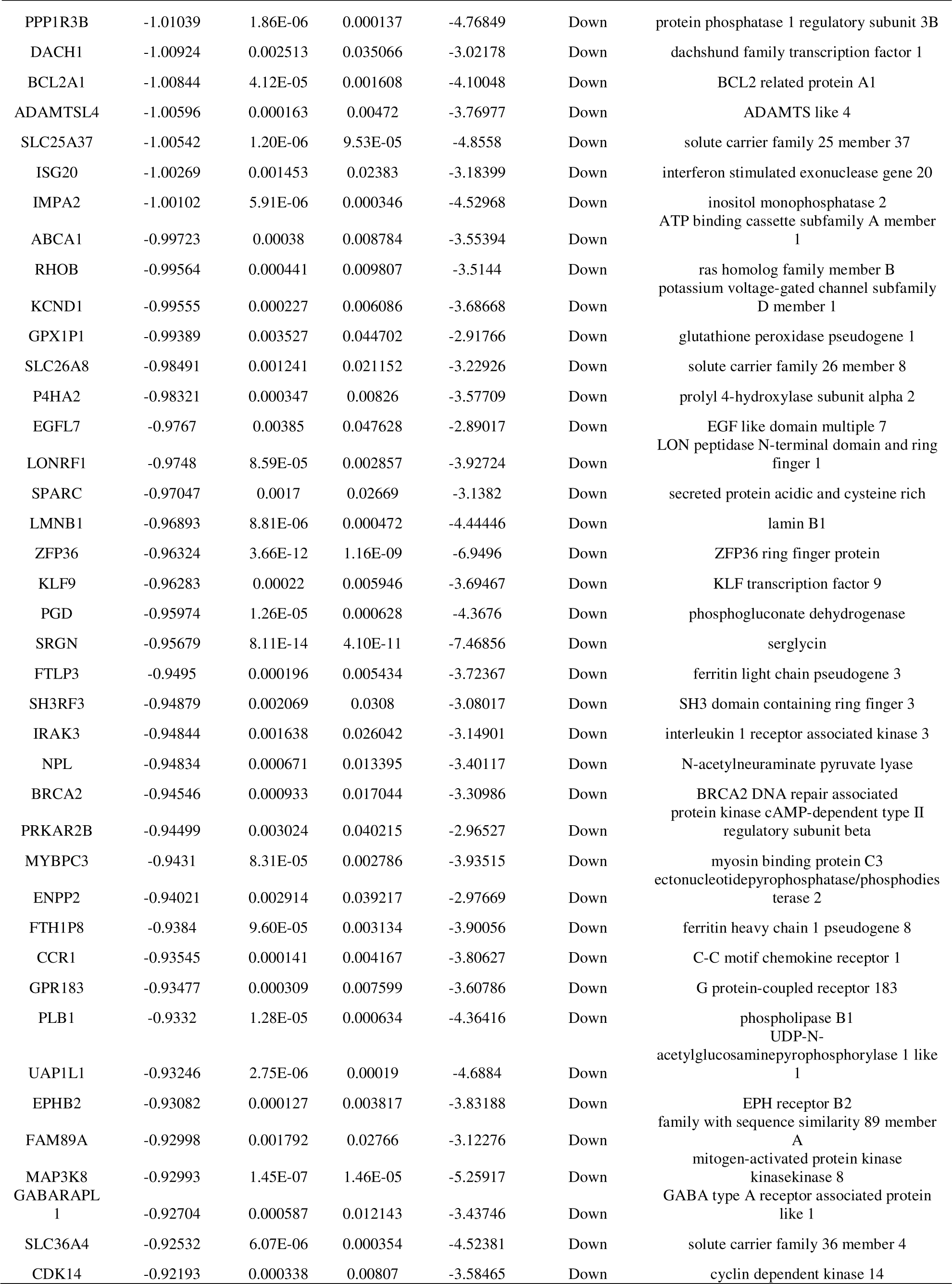
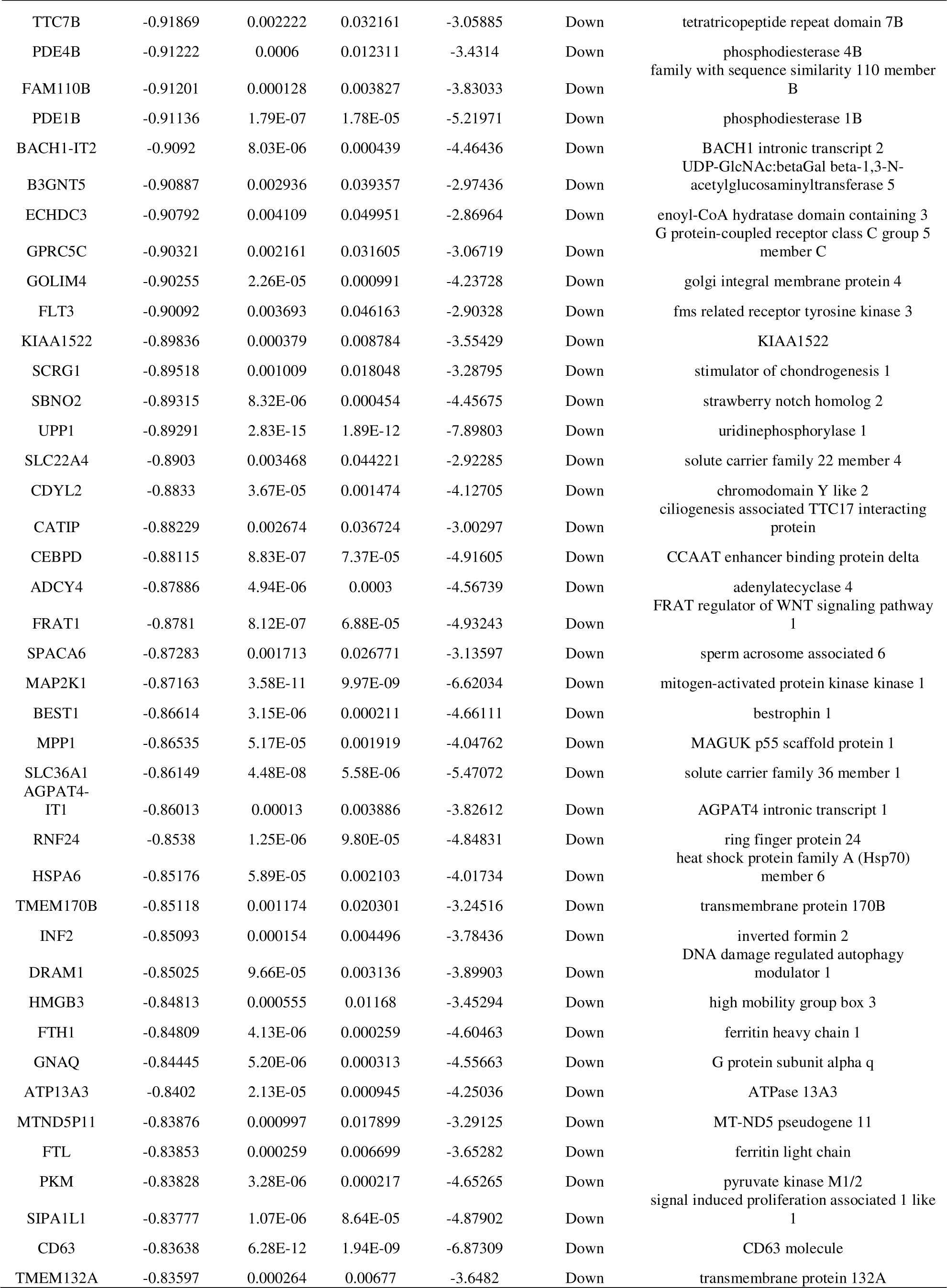
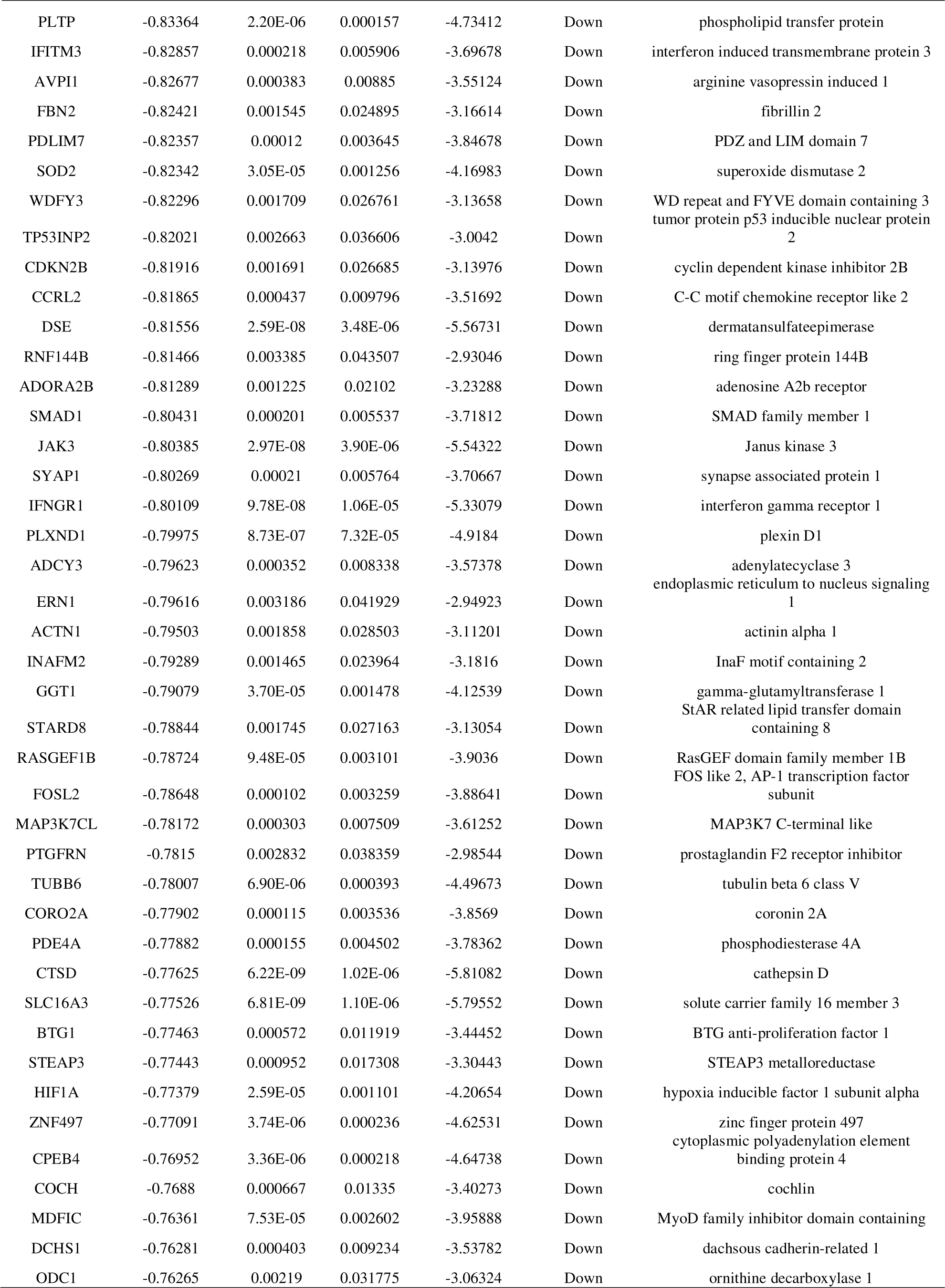
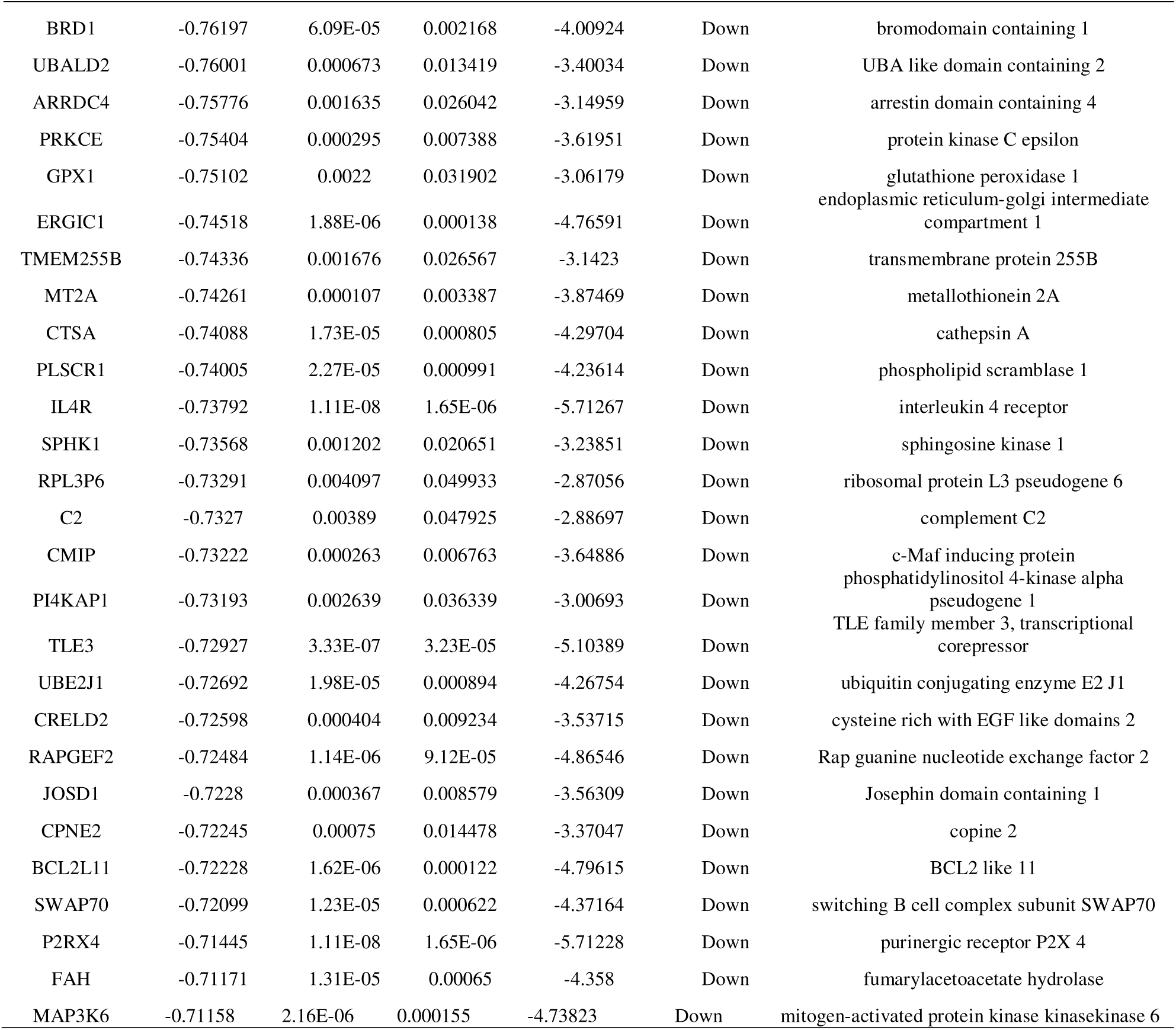
The statistical metrics for key differentially expressed genes (DEGs)

**Fig. 1.**
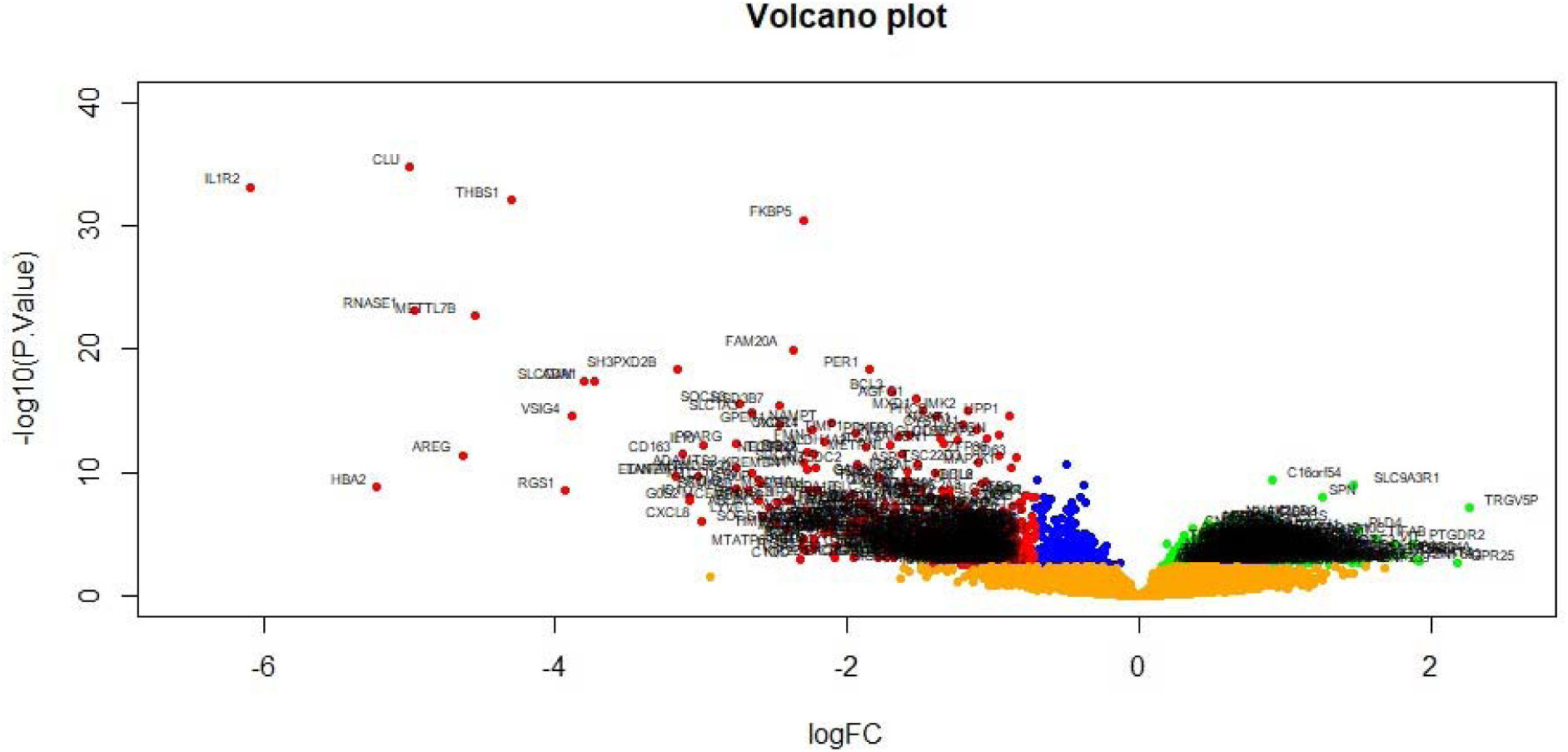
Volcano plot of differentially expressed genes. Genes with a significant change of more than two-fold were selected. Green dot represented up regulated significant genes and red dot represented down regulated significant genes.

**Fig. 2.**
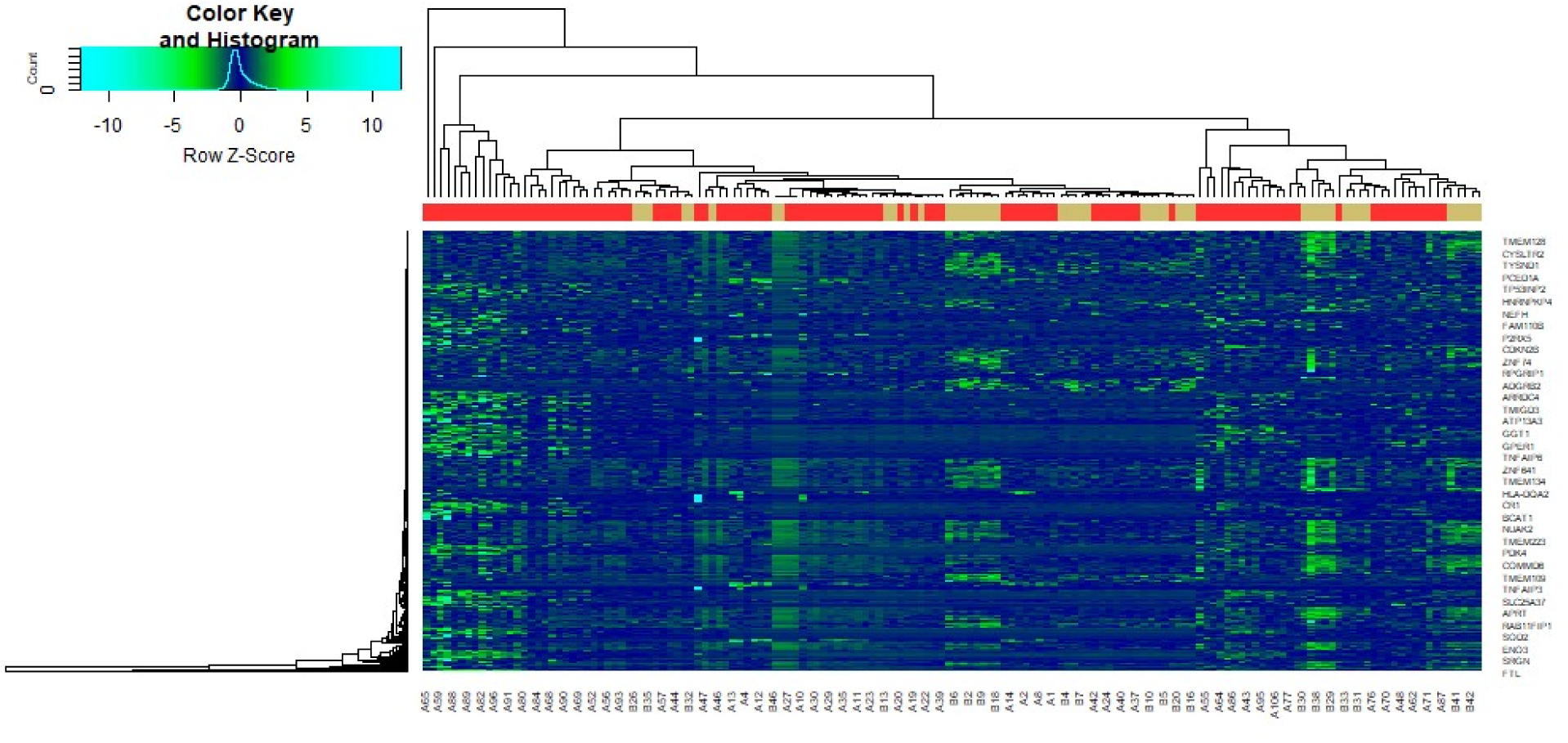
Heat map of differentially expressed genes. Legend on the top left indicate log fold change of genes. (A1 – A106 = HD samples; B1 – B46 = Normal control samples)

### GO and pathway enrichment analyses of DEGs

A total of 844 DEGs were uploaded to g:Profiler for GO term and REACTOME pathway enrichment analyses. The terms of each GO terms are provided in Table 2. Most DEGs were enriched in the BP include organonitrogen compound metabolic process, biosynthetic process, response to stimulus and cell communication; the CC include intracellular anatomical structure, organelle, cytoplasm and extracellular region; and the MF include catalytic activity, purine nucleotide binding, signaling receptor binding and molecular function regulator activity. The results of REACTOME pathway enrichment are shown in Table 3. Most DEGs were enriched in the pathway include translation, metabolism of RNA, immune system and platelet activation, signaling and aggregation.

**Table 2.**
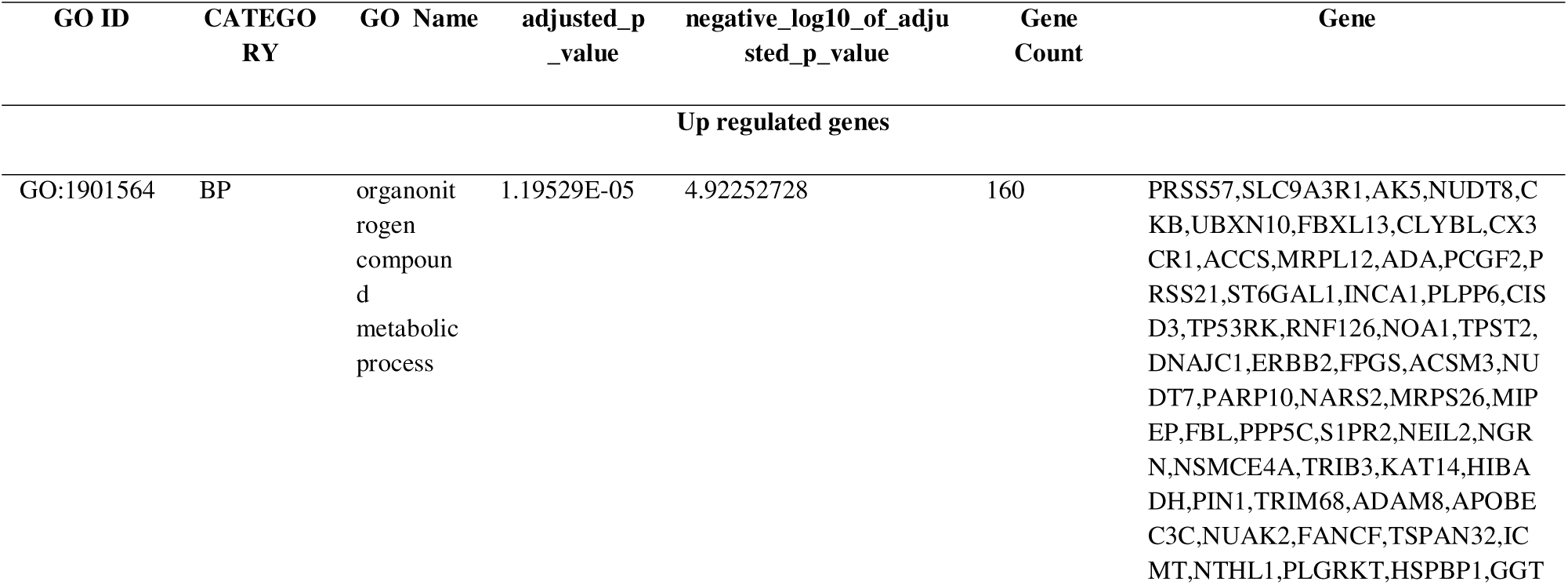
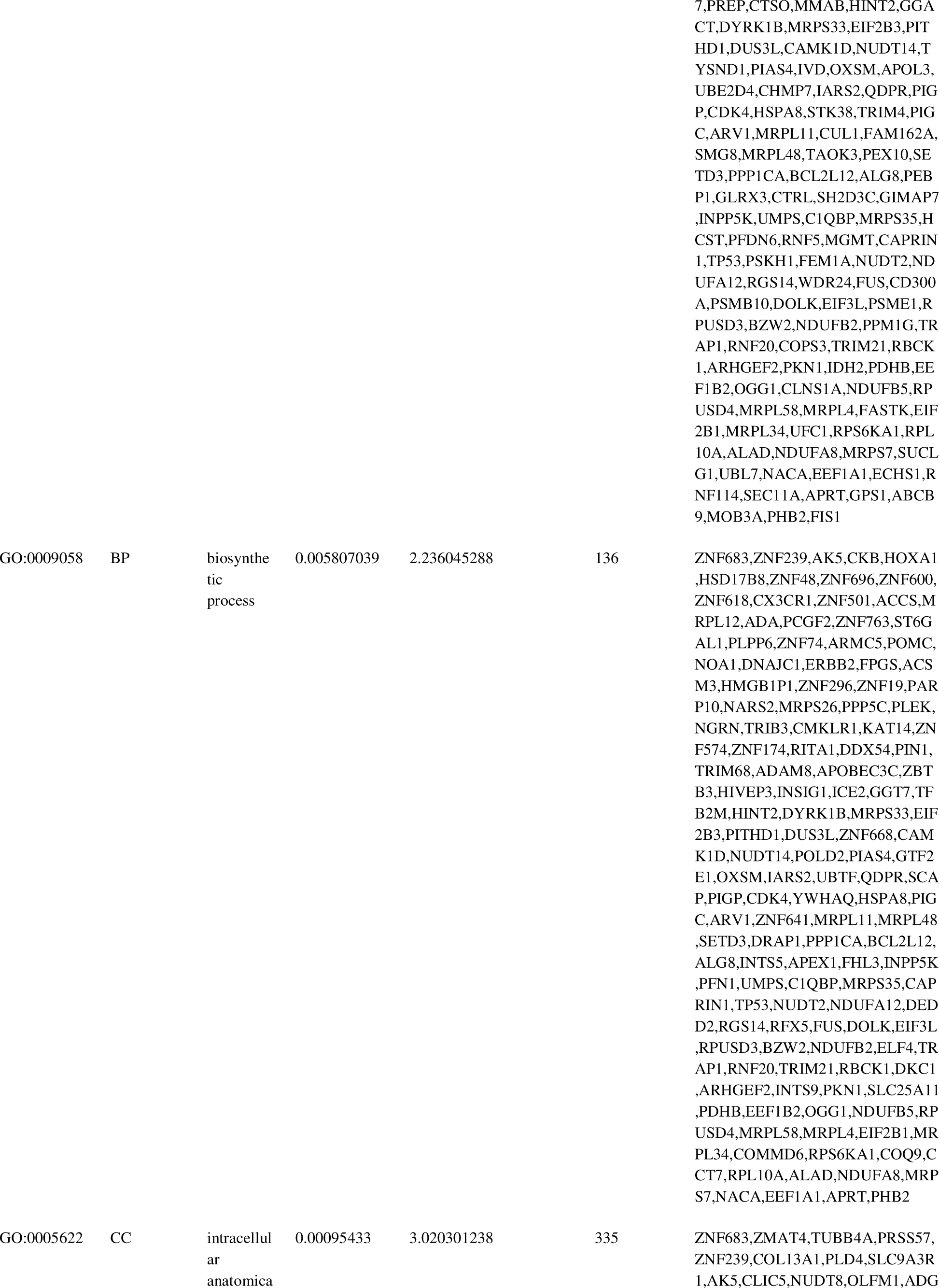
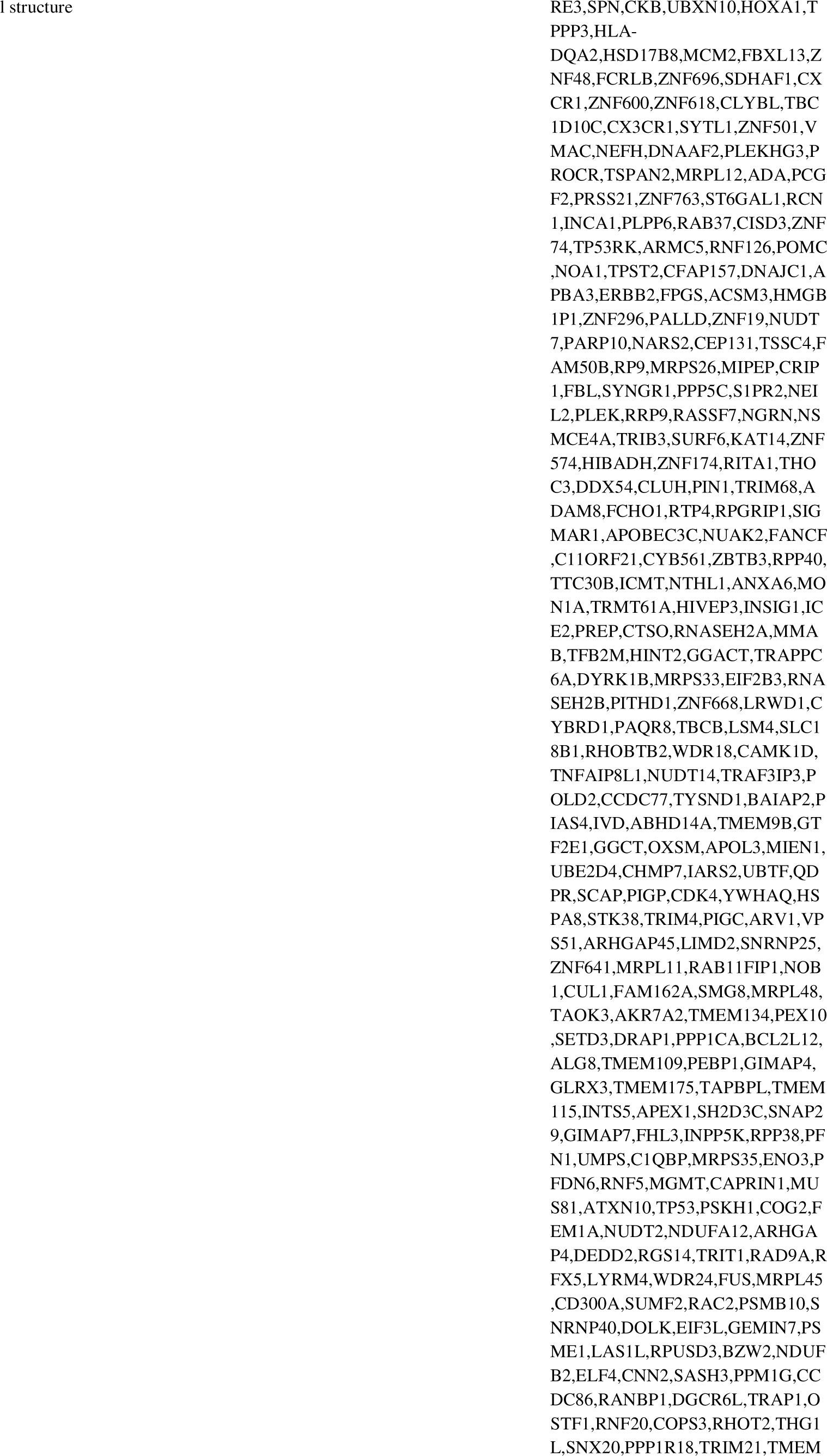
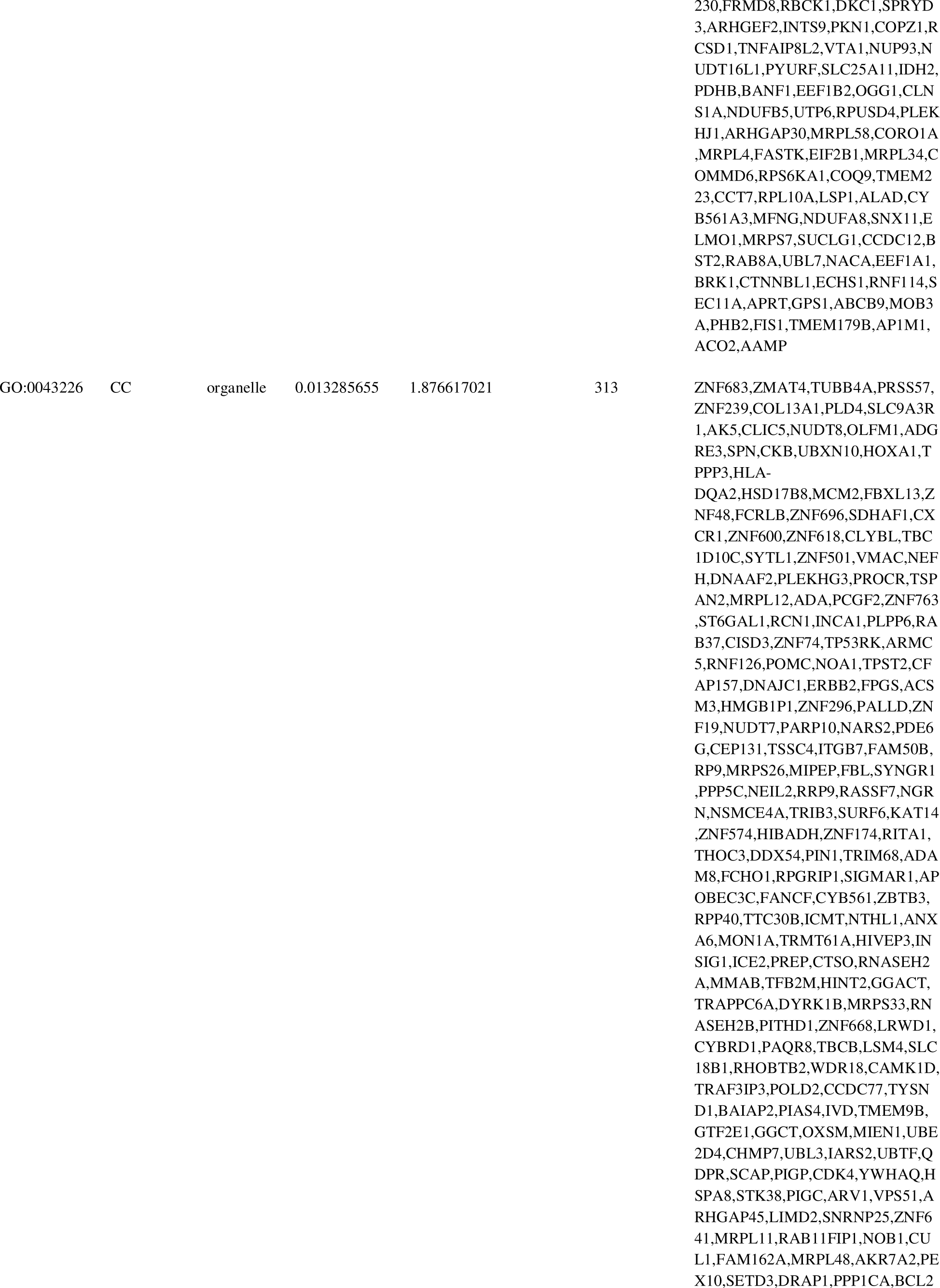
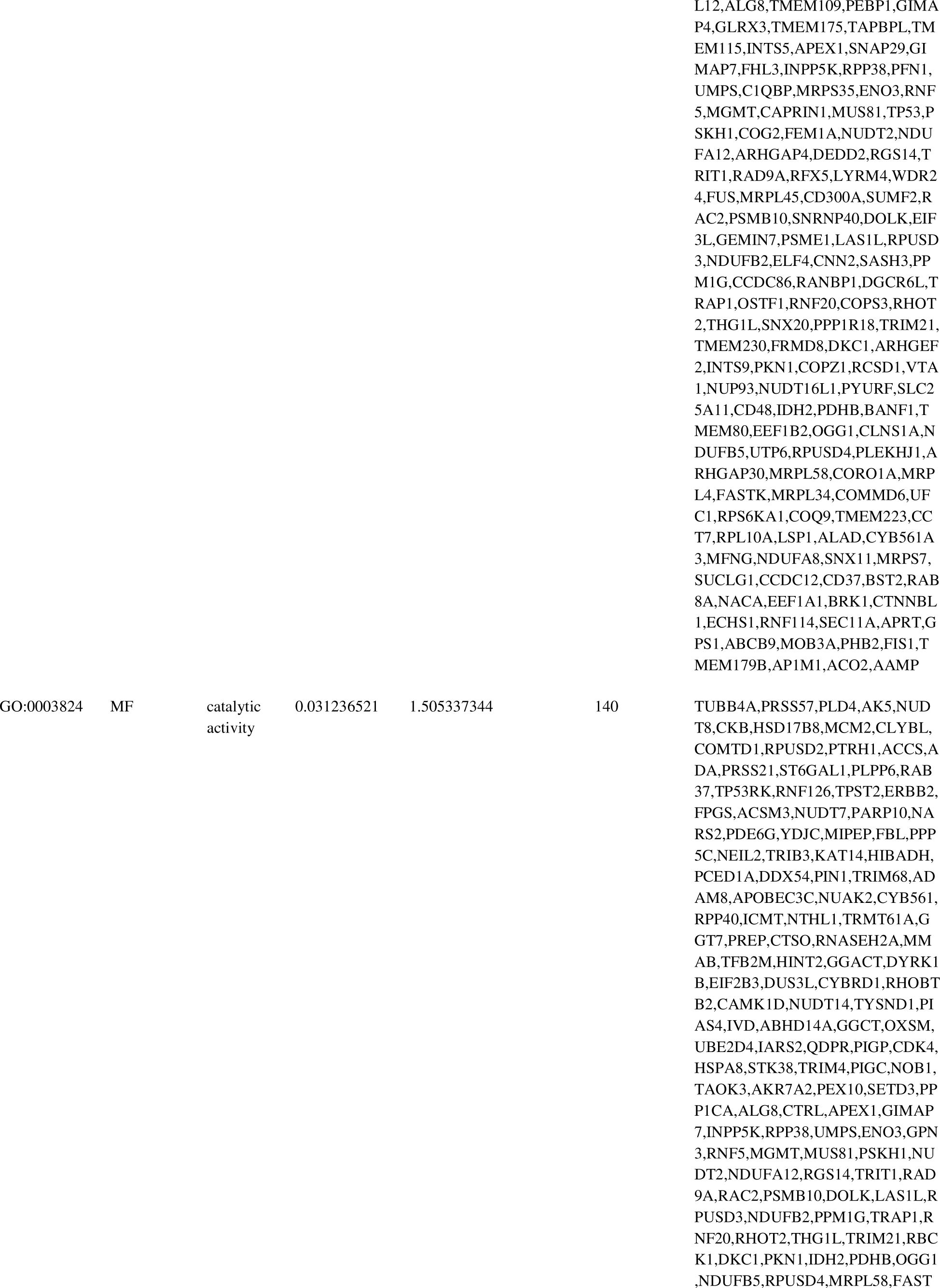
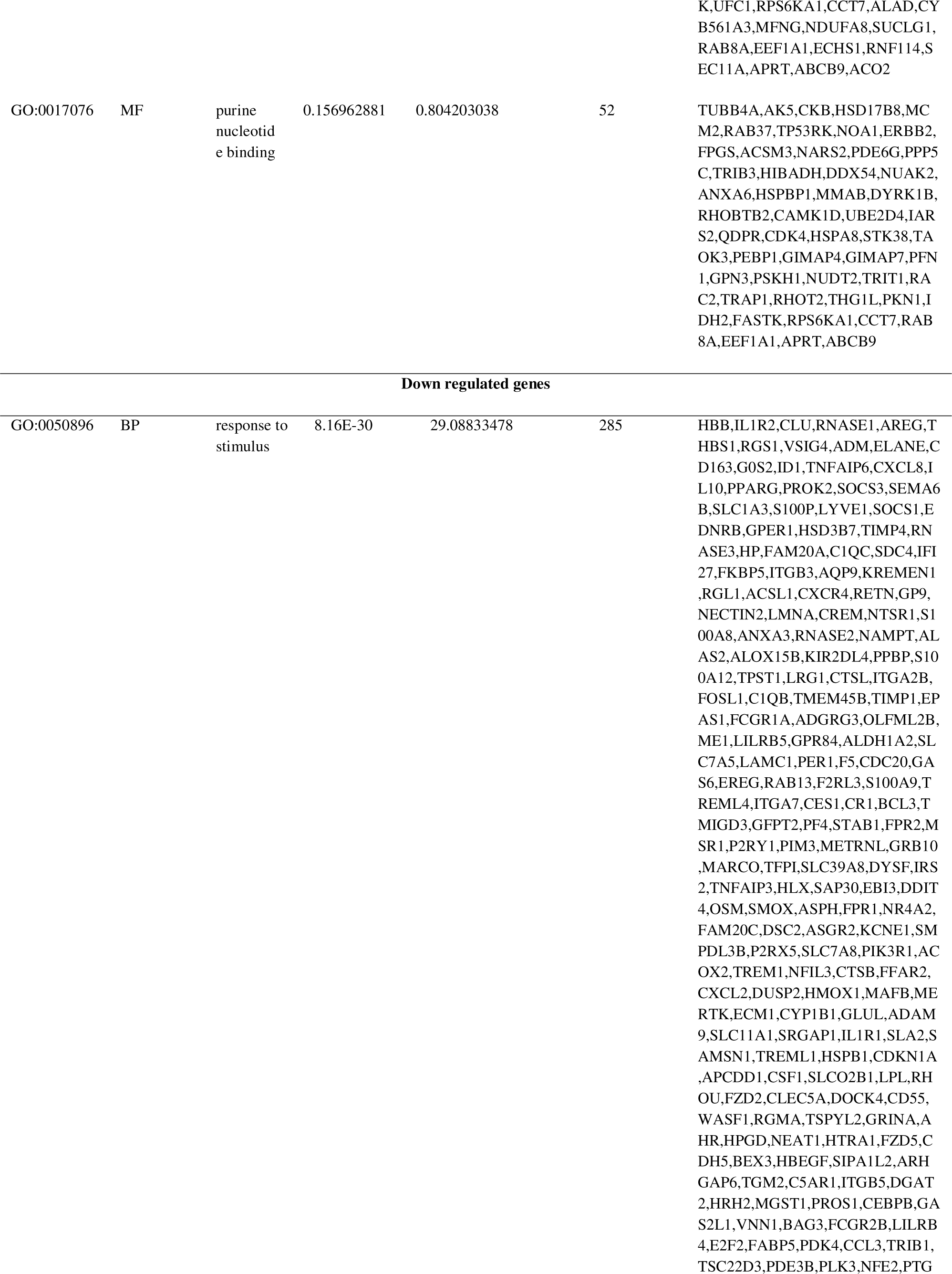
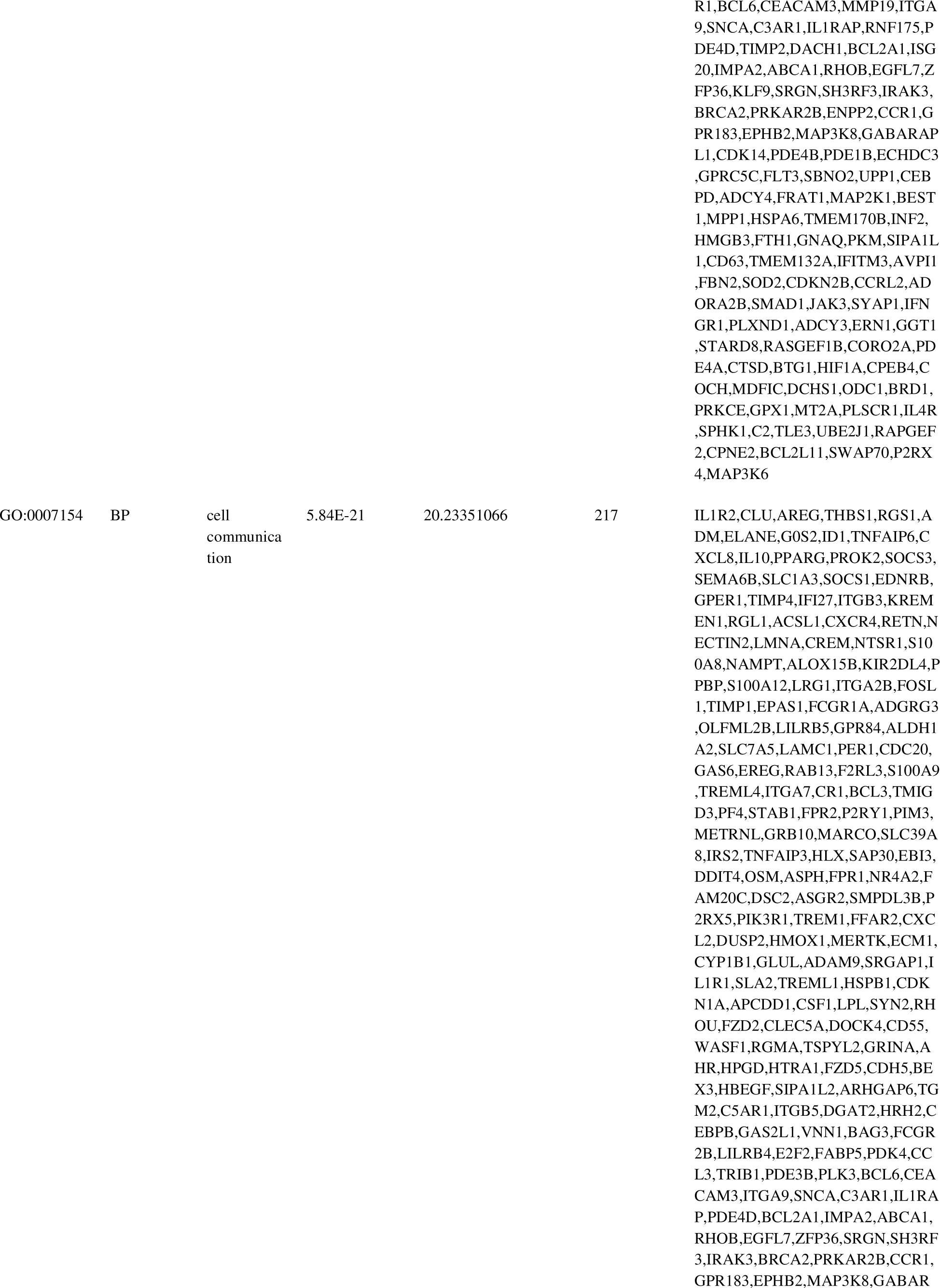
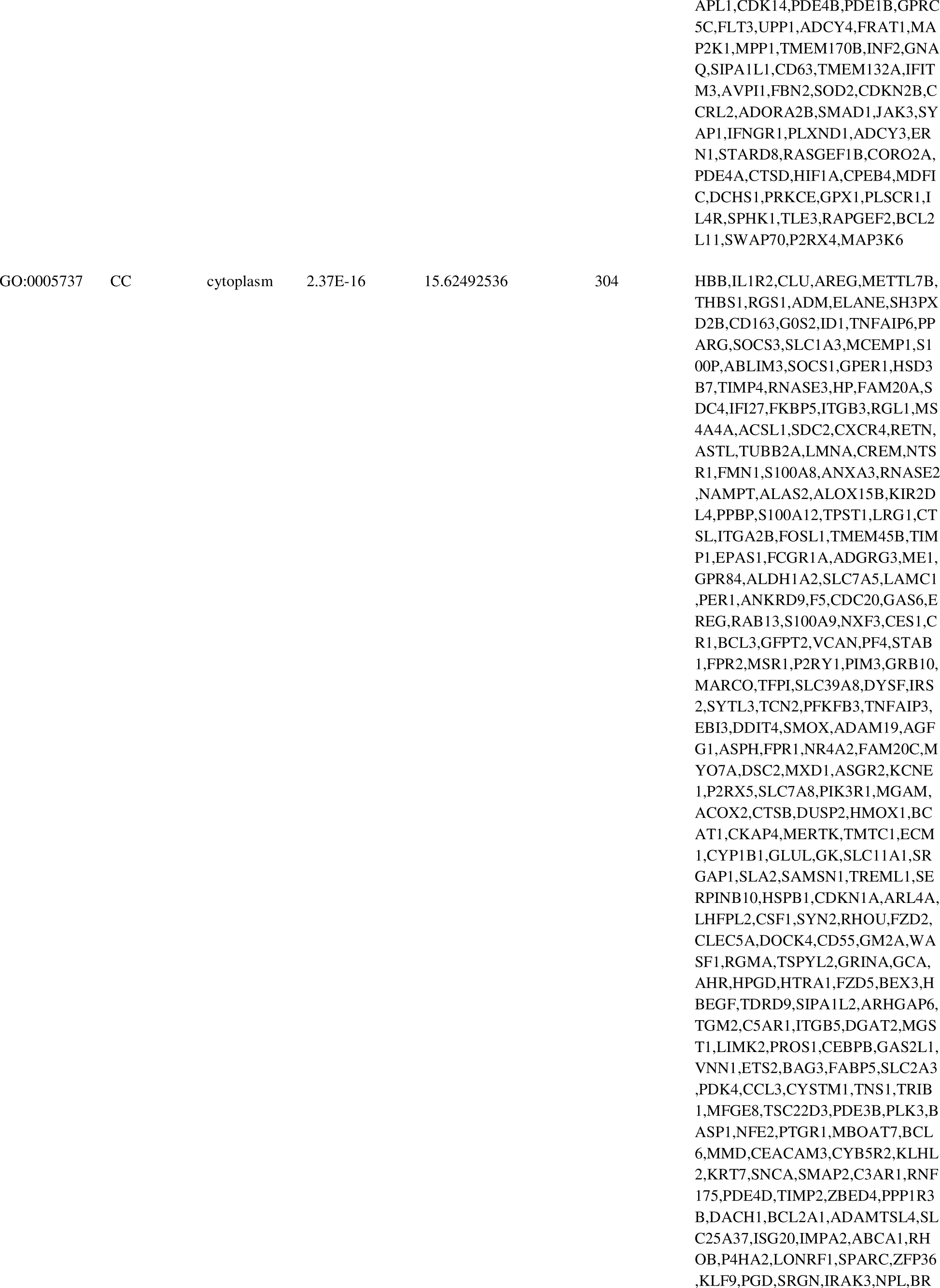
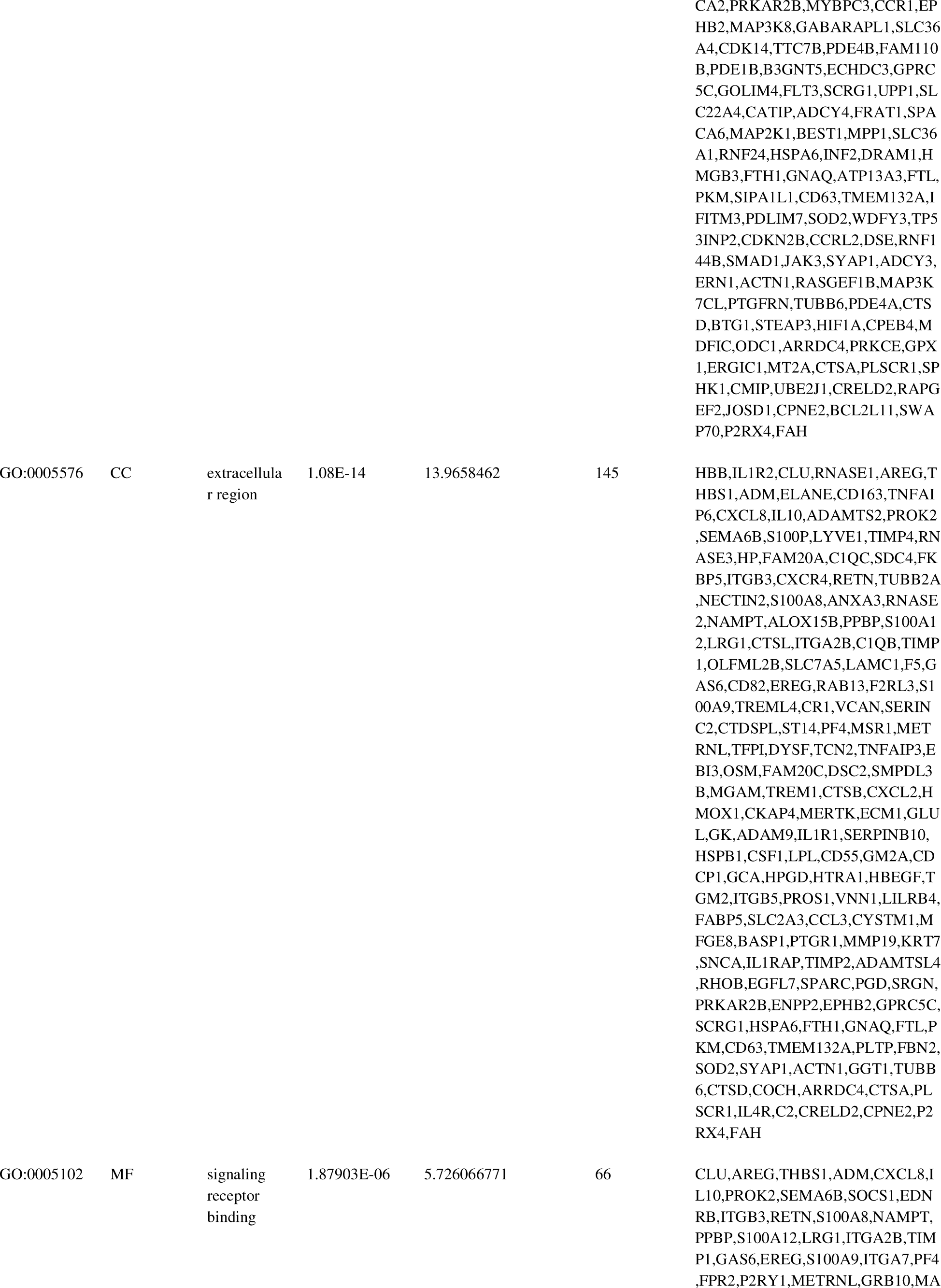
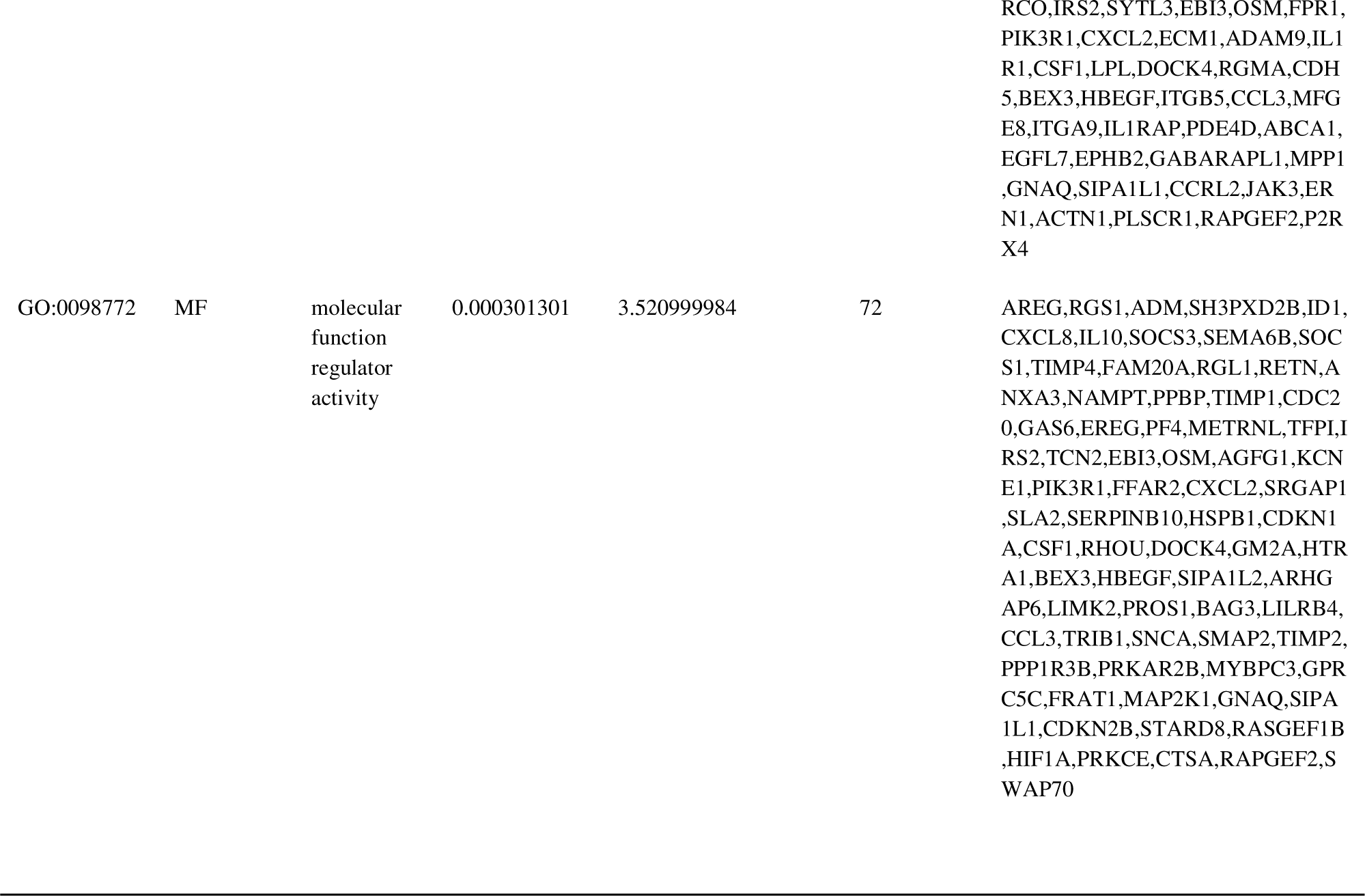
The enriched GO terms of the up and down regulated differentially expressed genes.

**Table 3.**
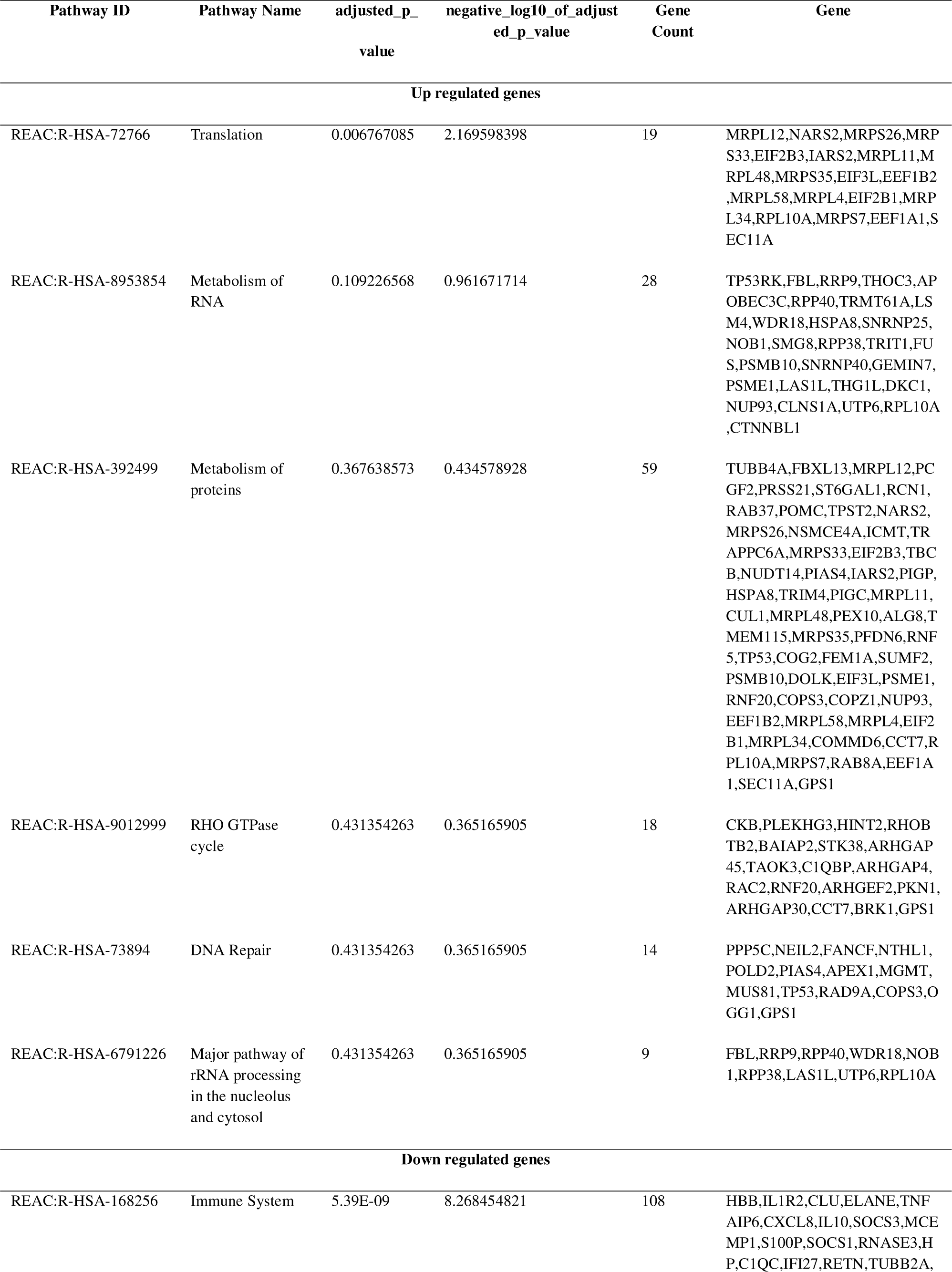
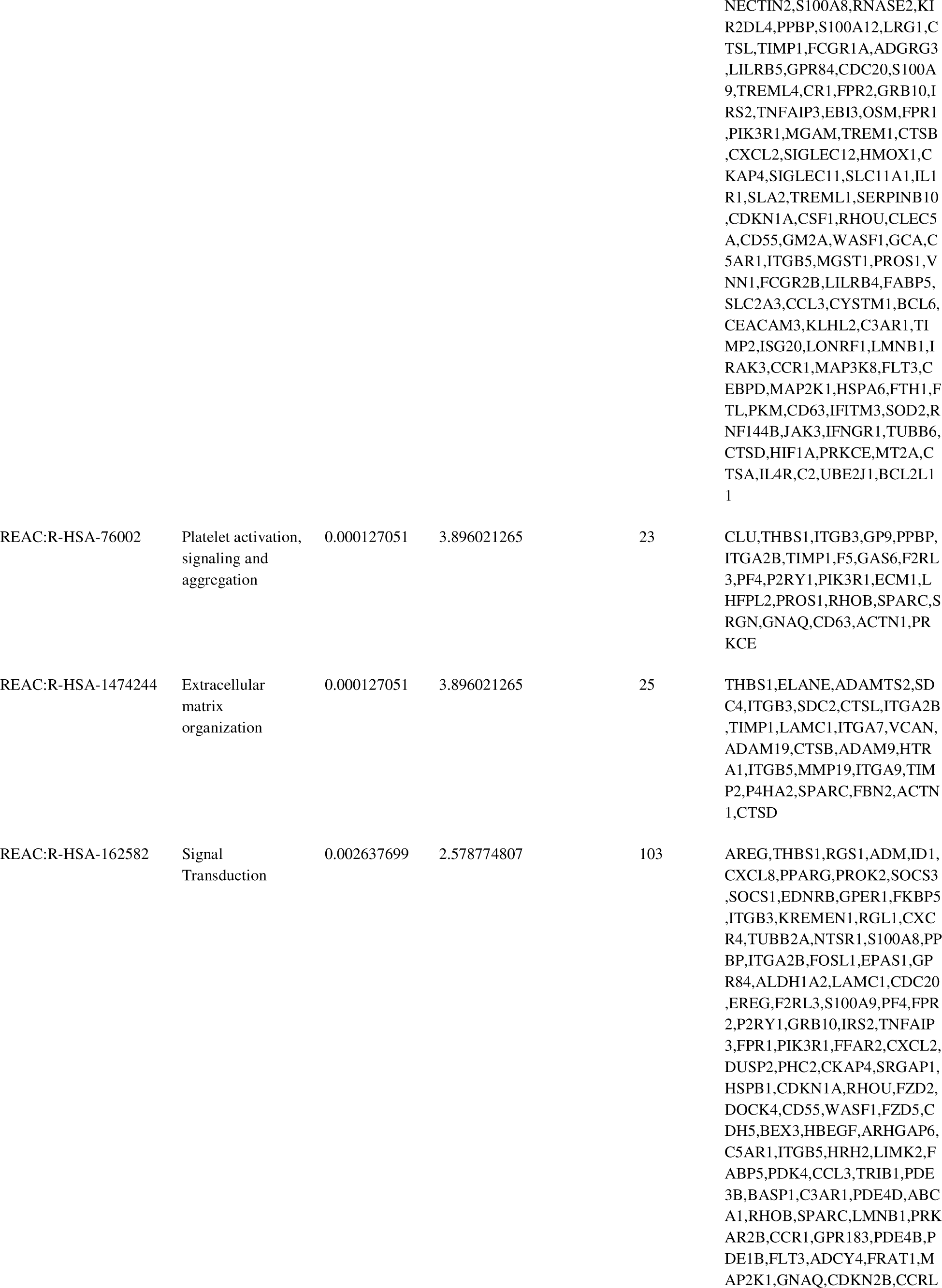
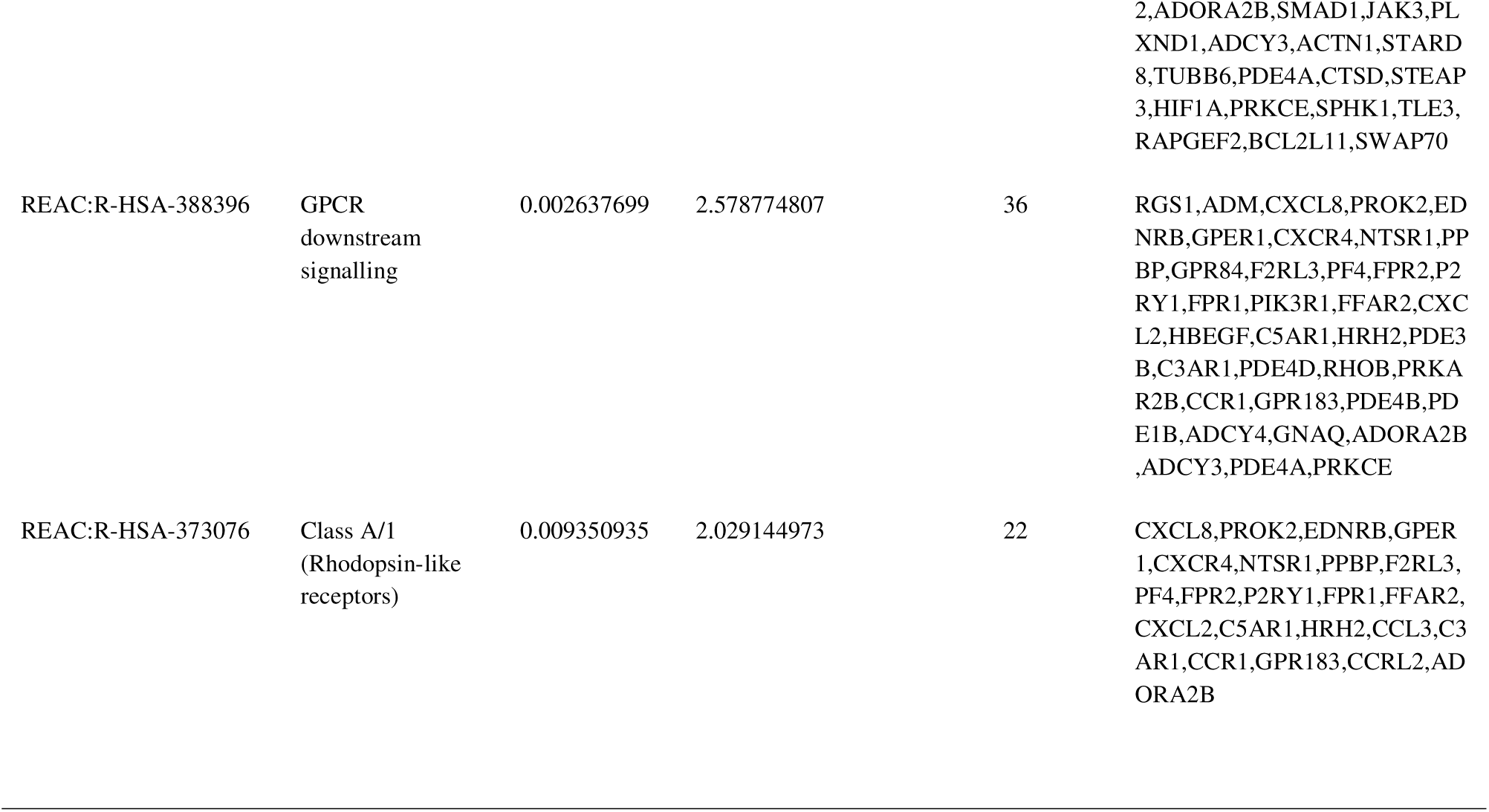
The enriched pathway terms of the up and down regulated differentially expressed genes.

### Construction of the PPI network and module analysis

A total of 844 DEGs were uploaded to the IID database to construct the PPI network. The PPI network is shown in Fig. 3. There were 7418 nodes and 15554 edges in the network. Then the most remarkable PPI was recognized by Network Analyzer plug-in according to the maximal node degree, betweenness, stress and closeness centrality topology analysis methods. The top 10 significant nodes with the highest connectivity among the network were appraised as hub genes: CUL1, HSPA8, HOXA1, INCA1, TP53, HSPB1, LMNA, SNCA, ADAMTSL4 and PDLIM7 (Table 4). Two closely connected gene modules were obtained through PEWCC plug-in of Cytoscape. All genes of these two modules belonged to the up and down regulated genes. The hub genes in each module were analyzed using g:Profiler to produce their relative GO term and REACTOME pathway enrichment. The module 1 harboring 58 nodes and 140 edges are shown in Fig. 4A. The module 2 harboring 88 nodes and 97 edges are shown in Fig. 4B. Module 1 was mainly enriched in metabolism of RNA, RHO GTPase cycle, metabolism of proteins, translation, organonitrogen compound metabolic process and biosynthetic process. Module 2 was mainly enriched in signal transduction, immune system, response to stimulus, cytoplasm, cell communication and extracellular region.

**Fig. 3.**
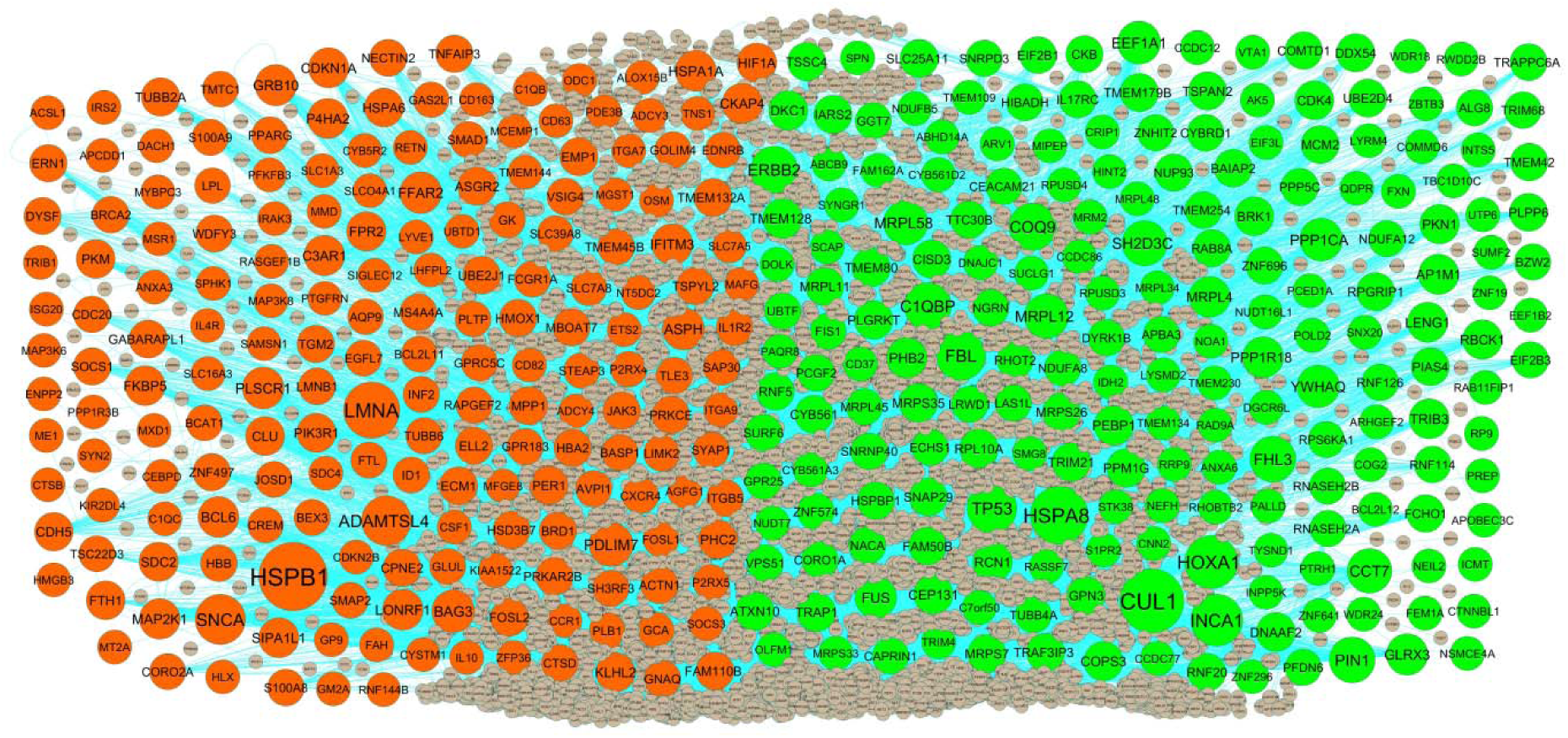
PPI network of DEGs. Up regulated genes are marked in parrot green; down regulated genes are marked in red.

**Table 4.**
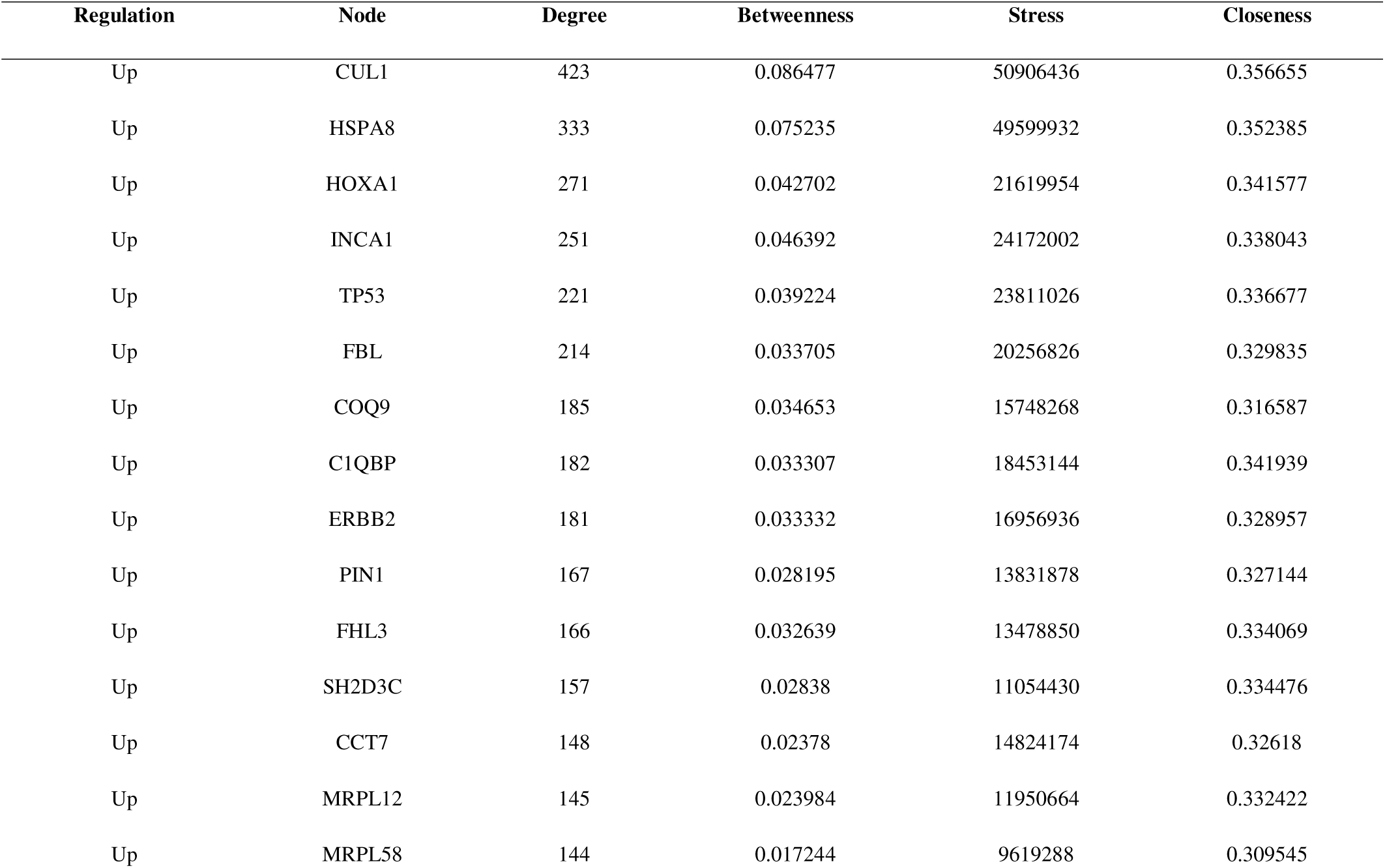
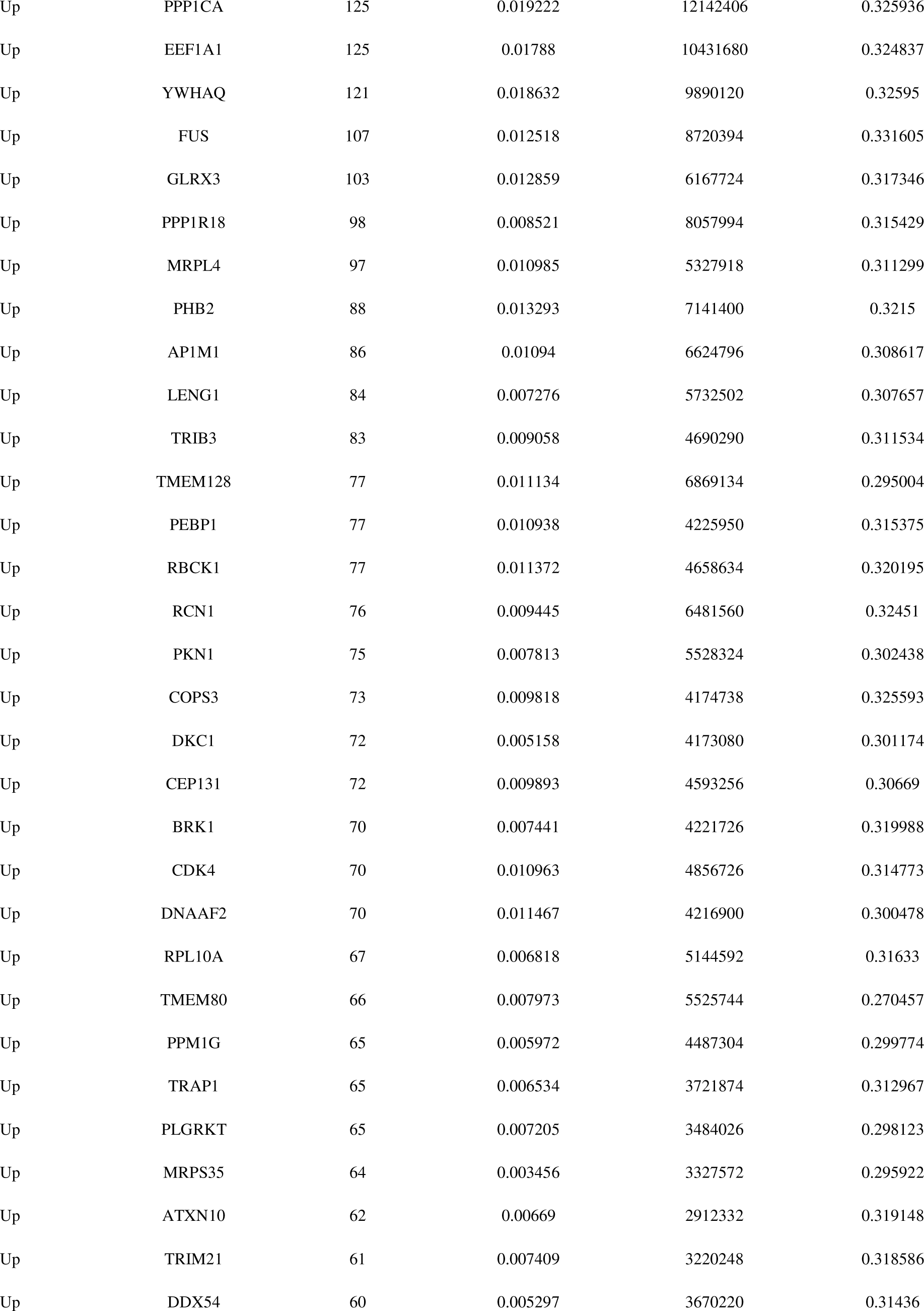
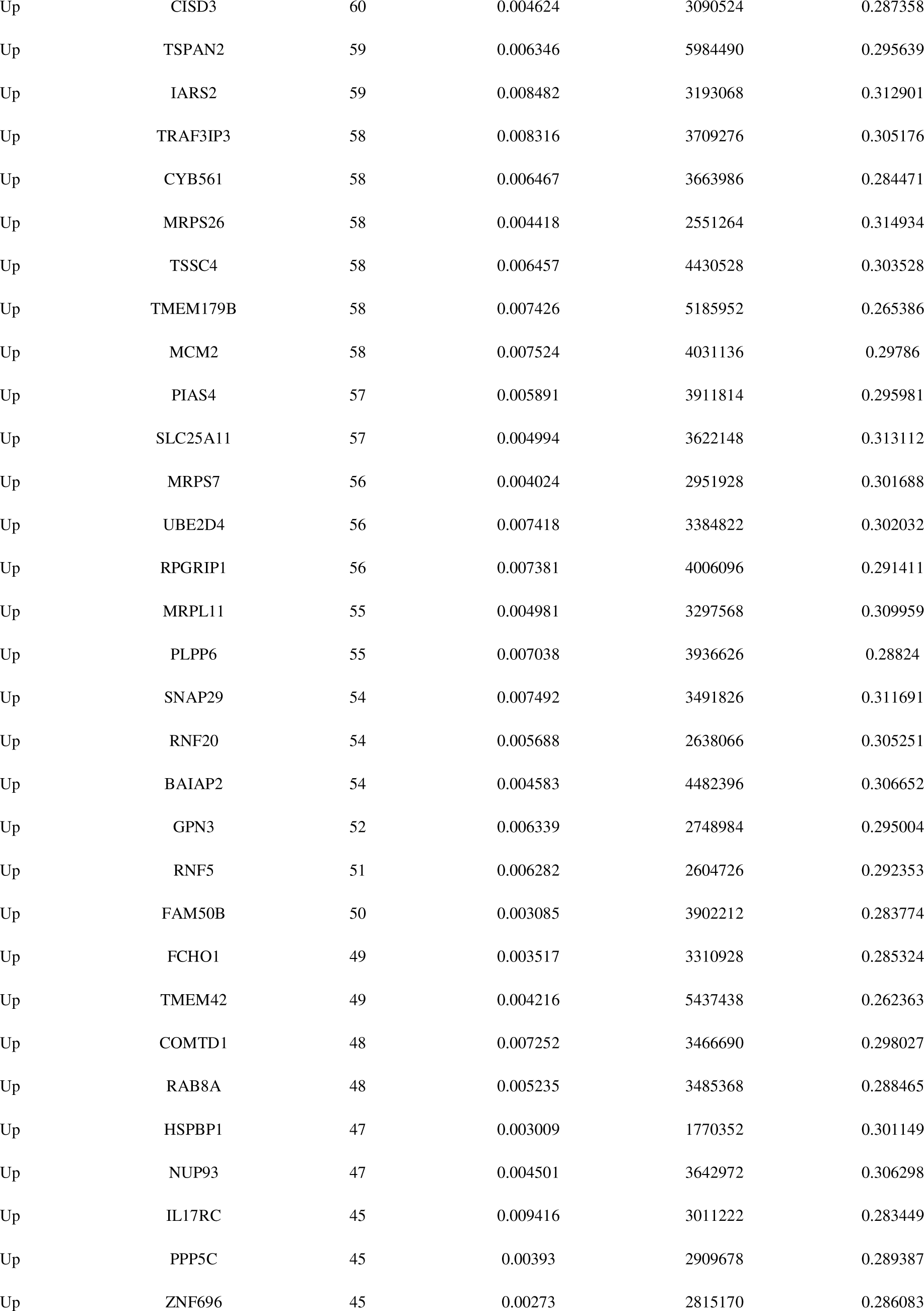
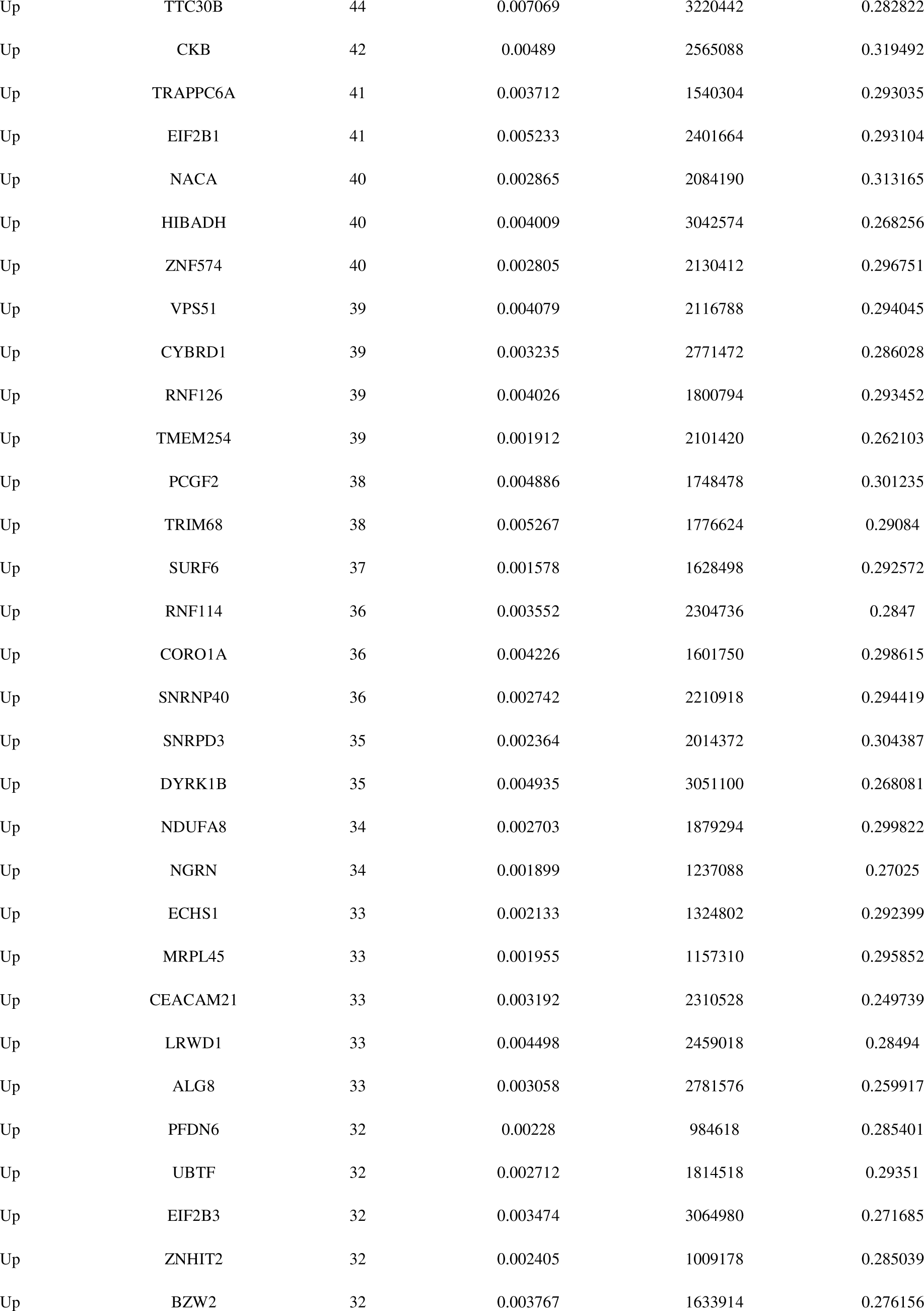
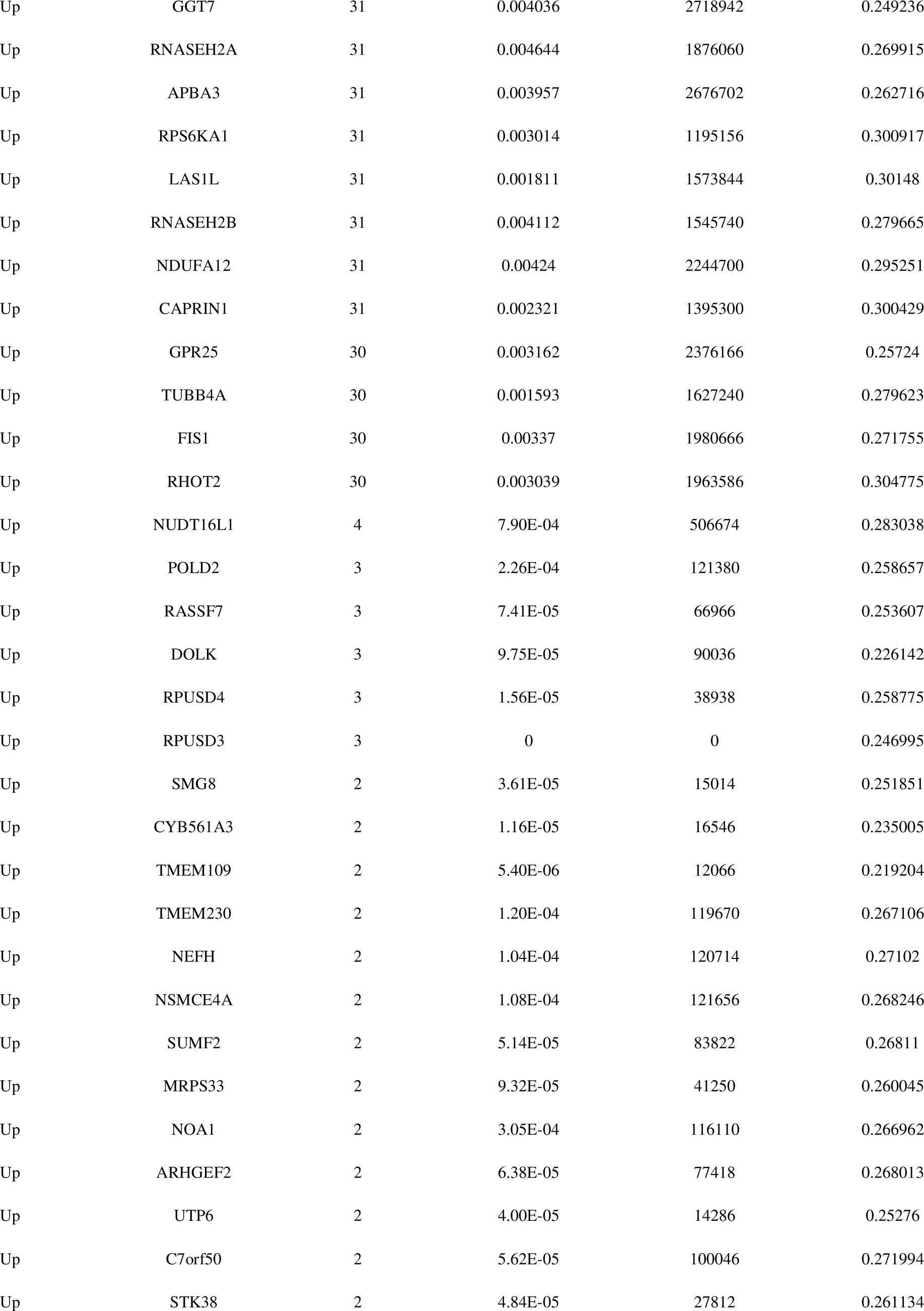
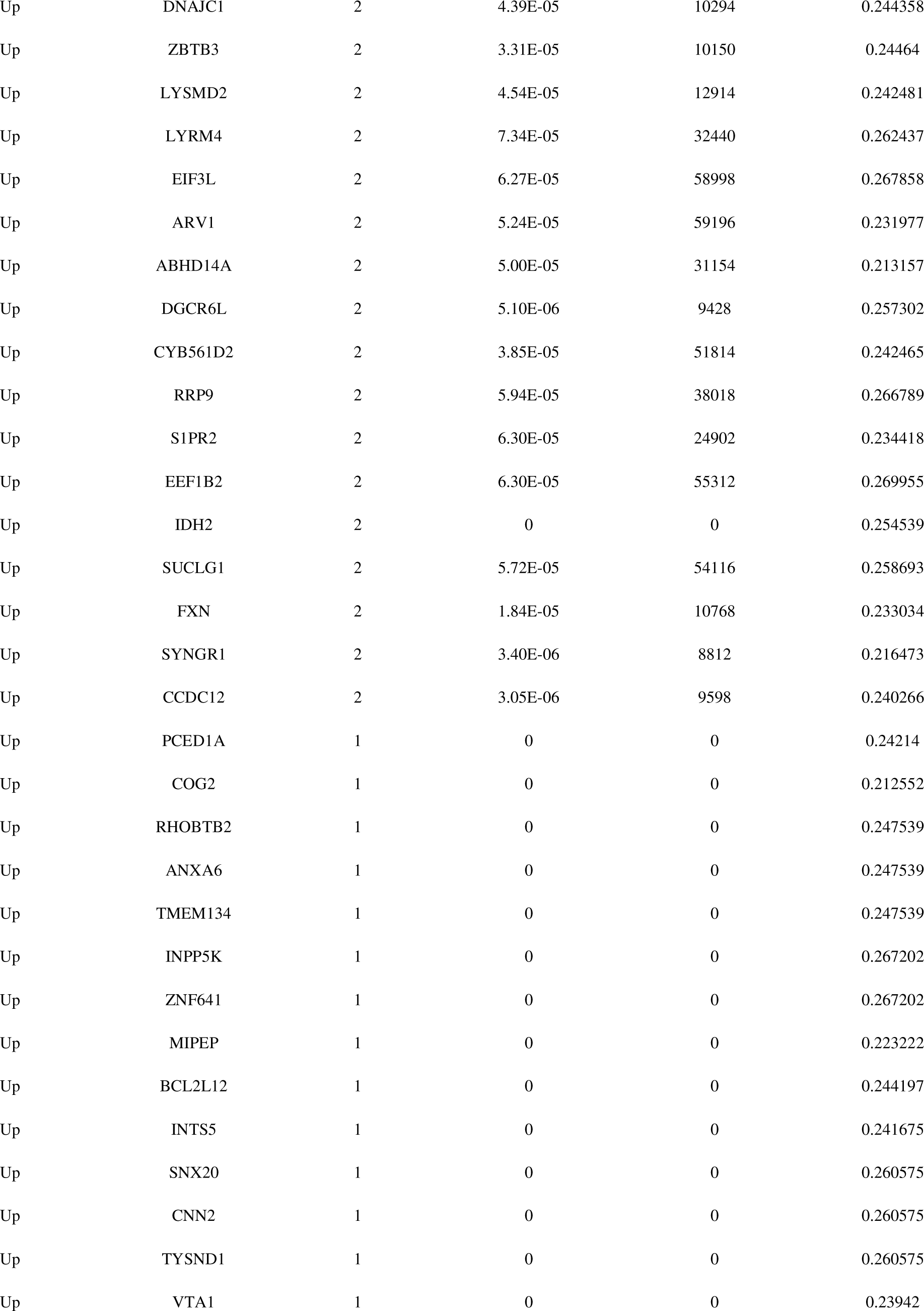
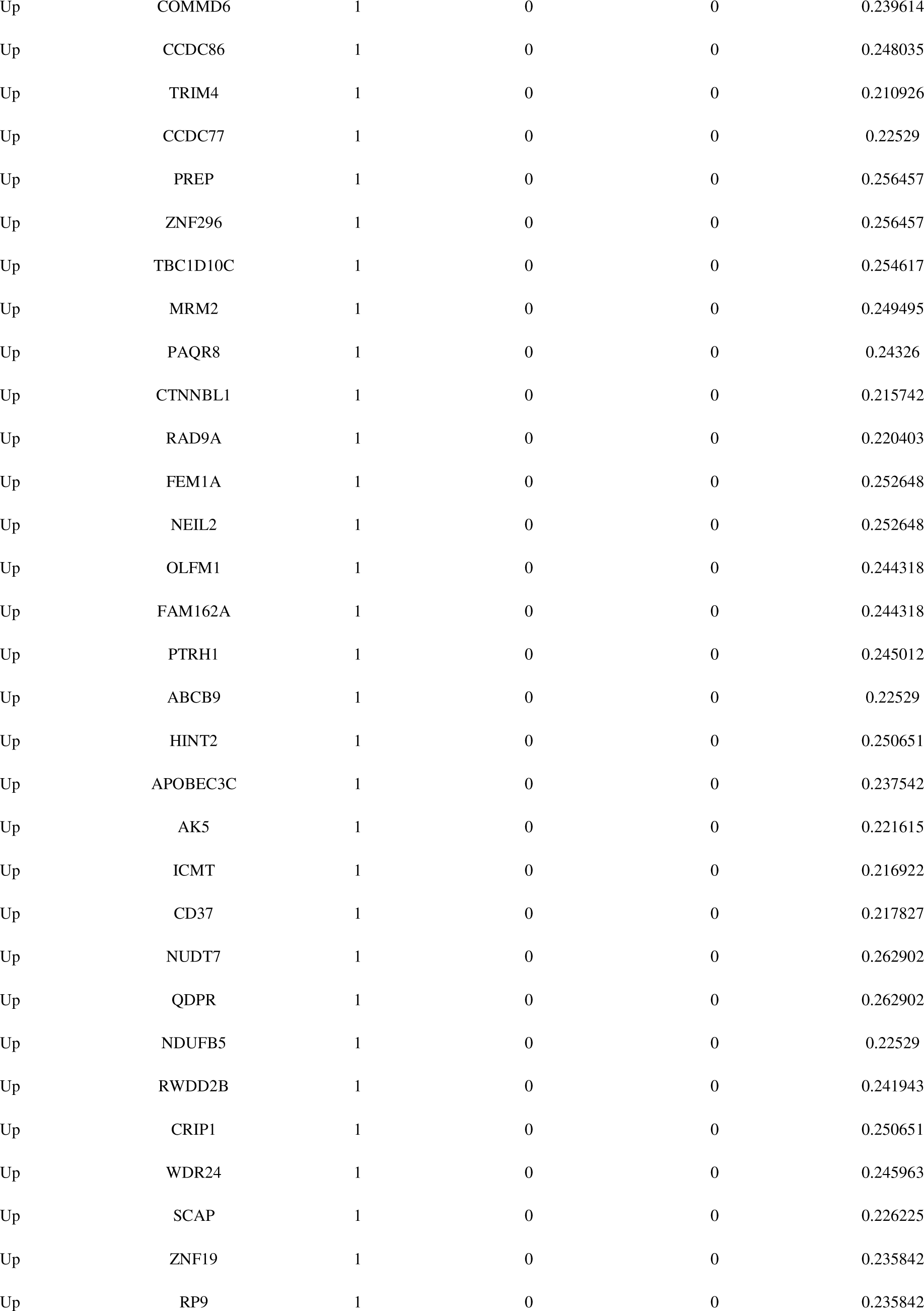
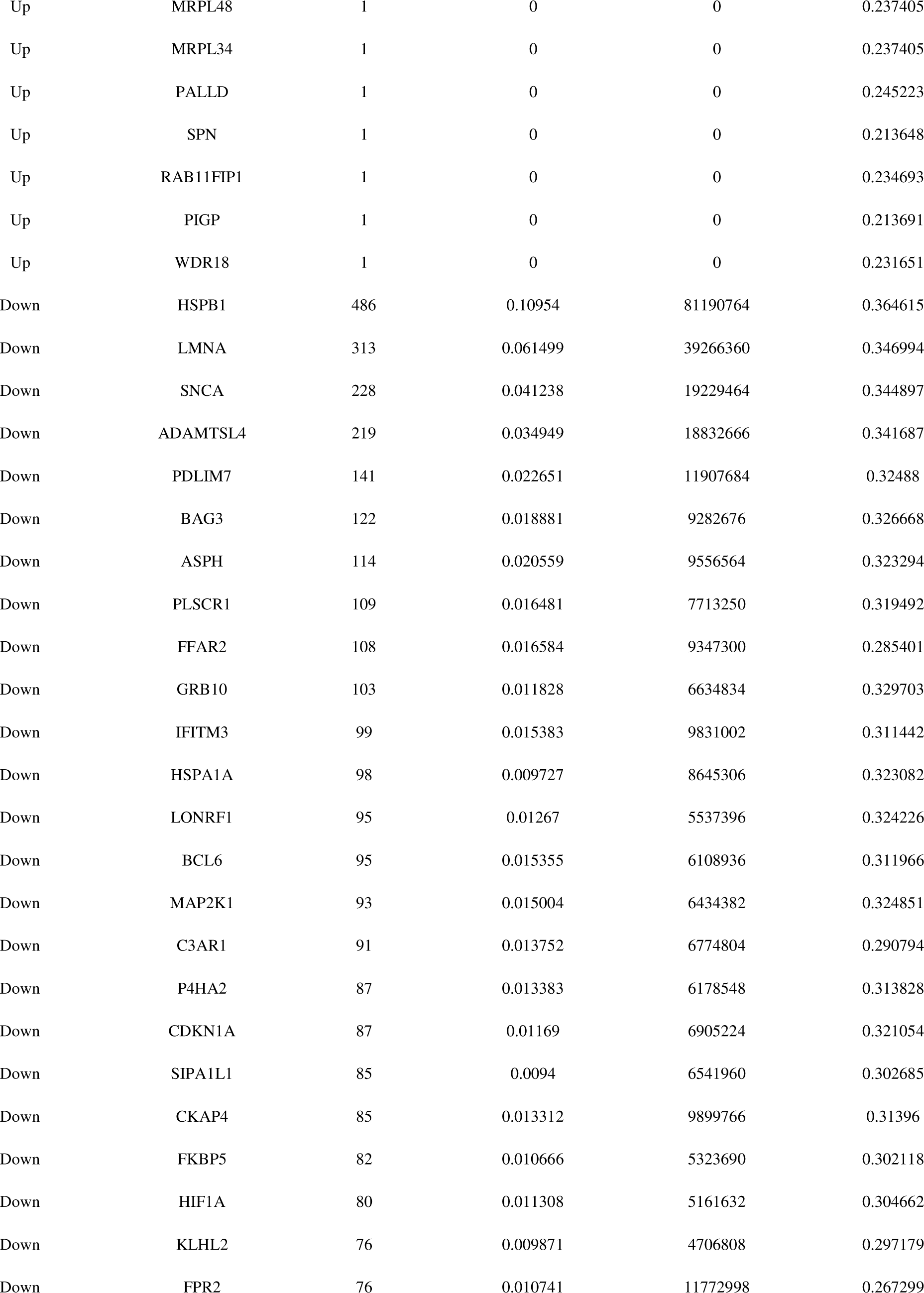
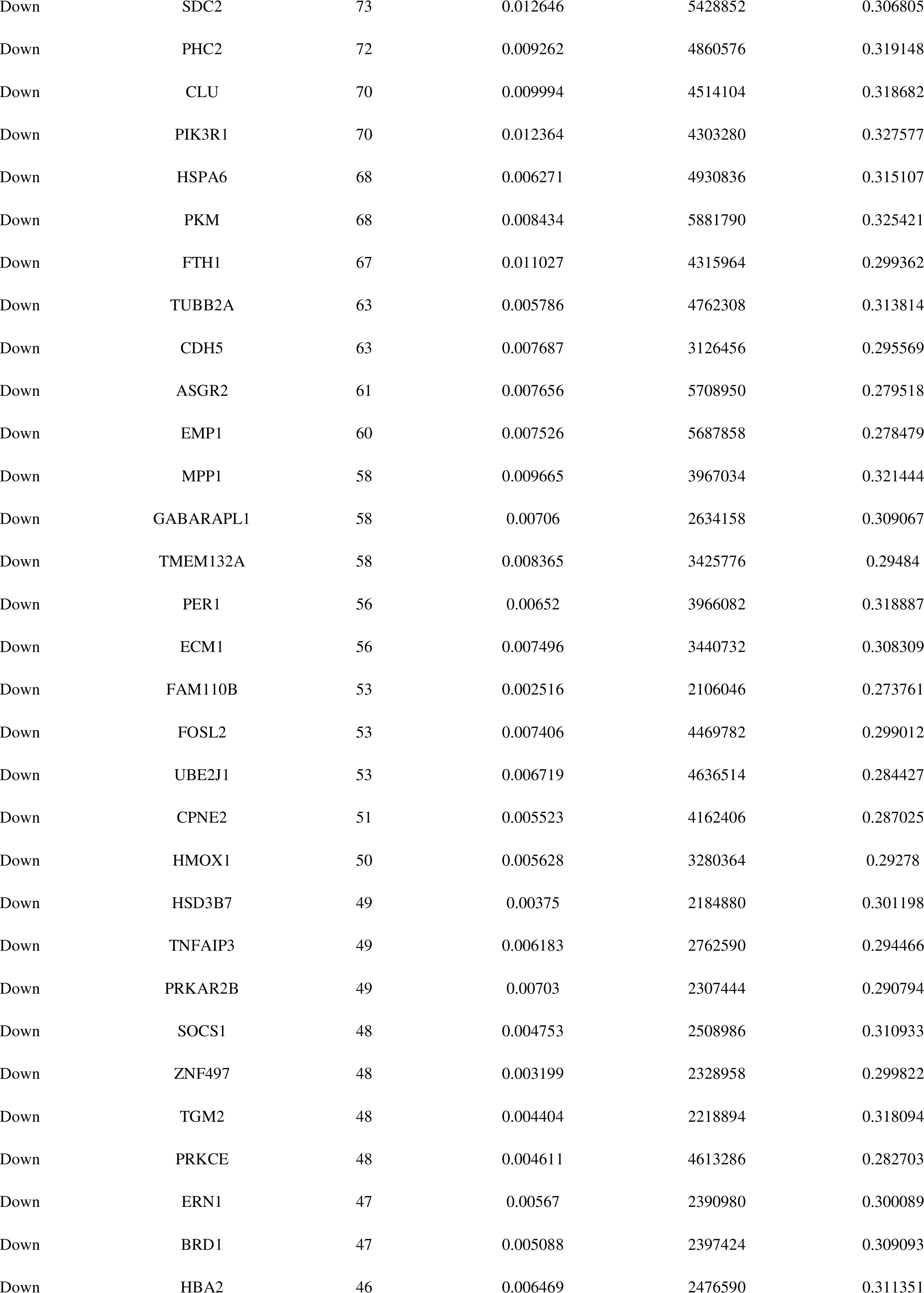
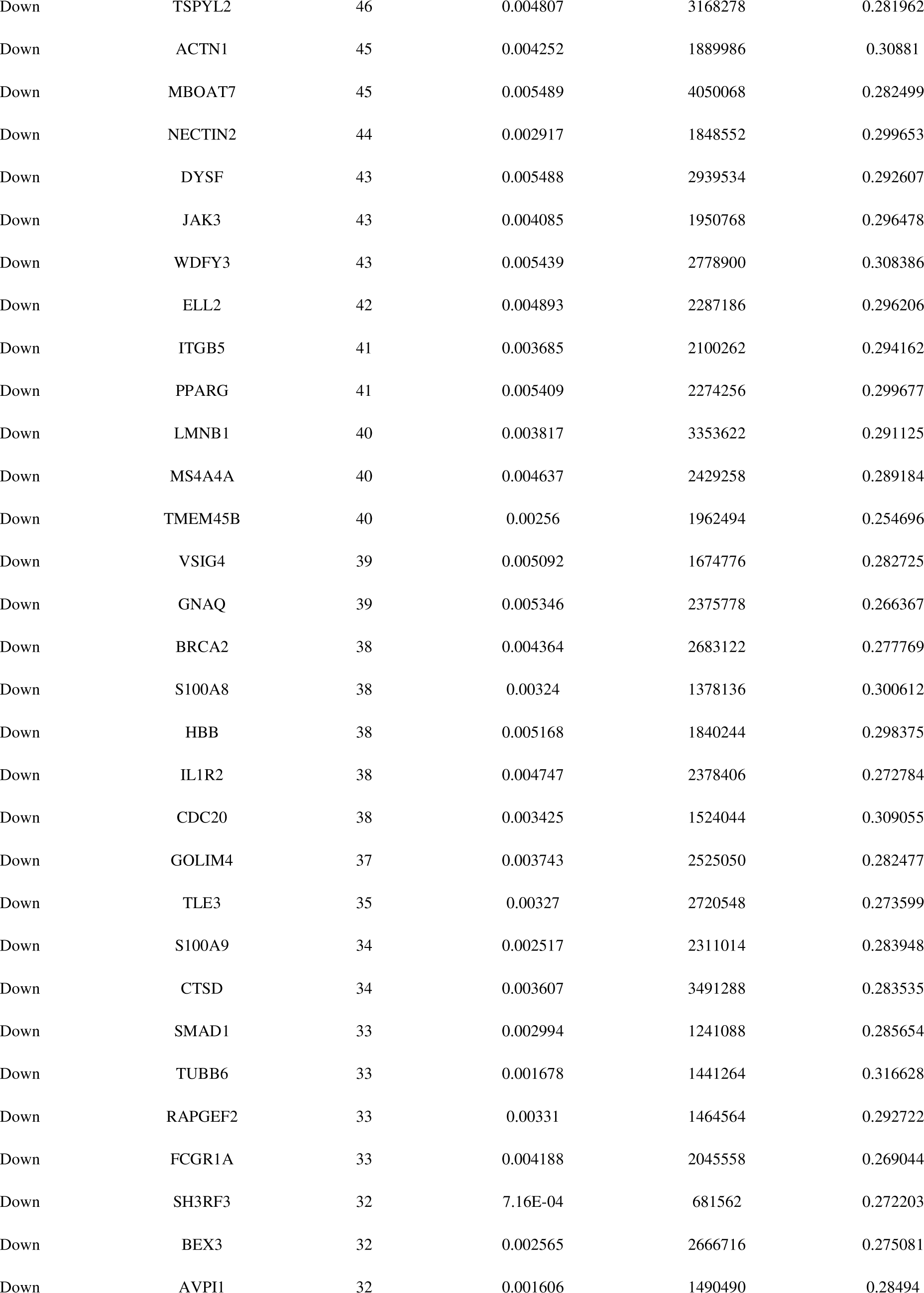
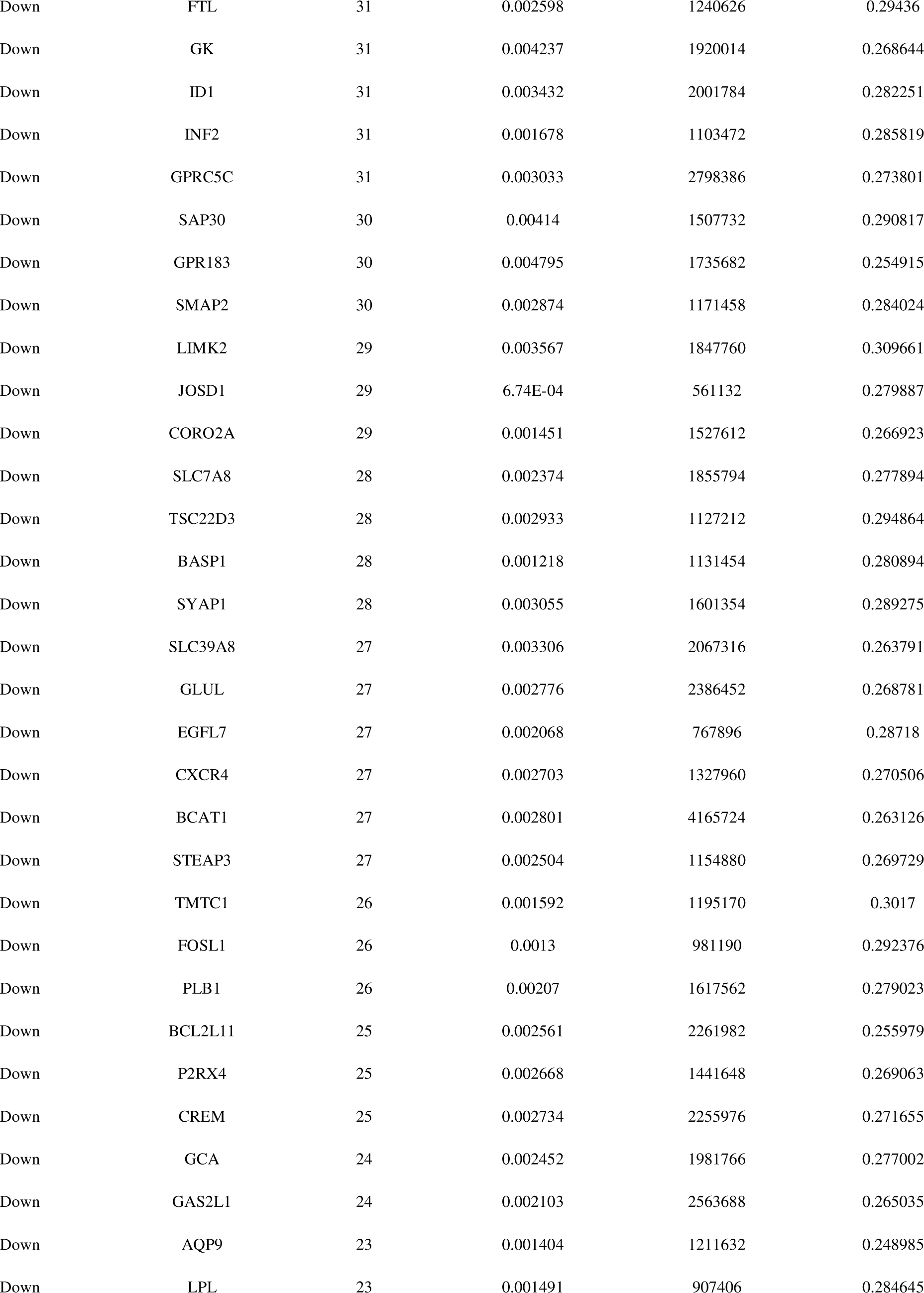
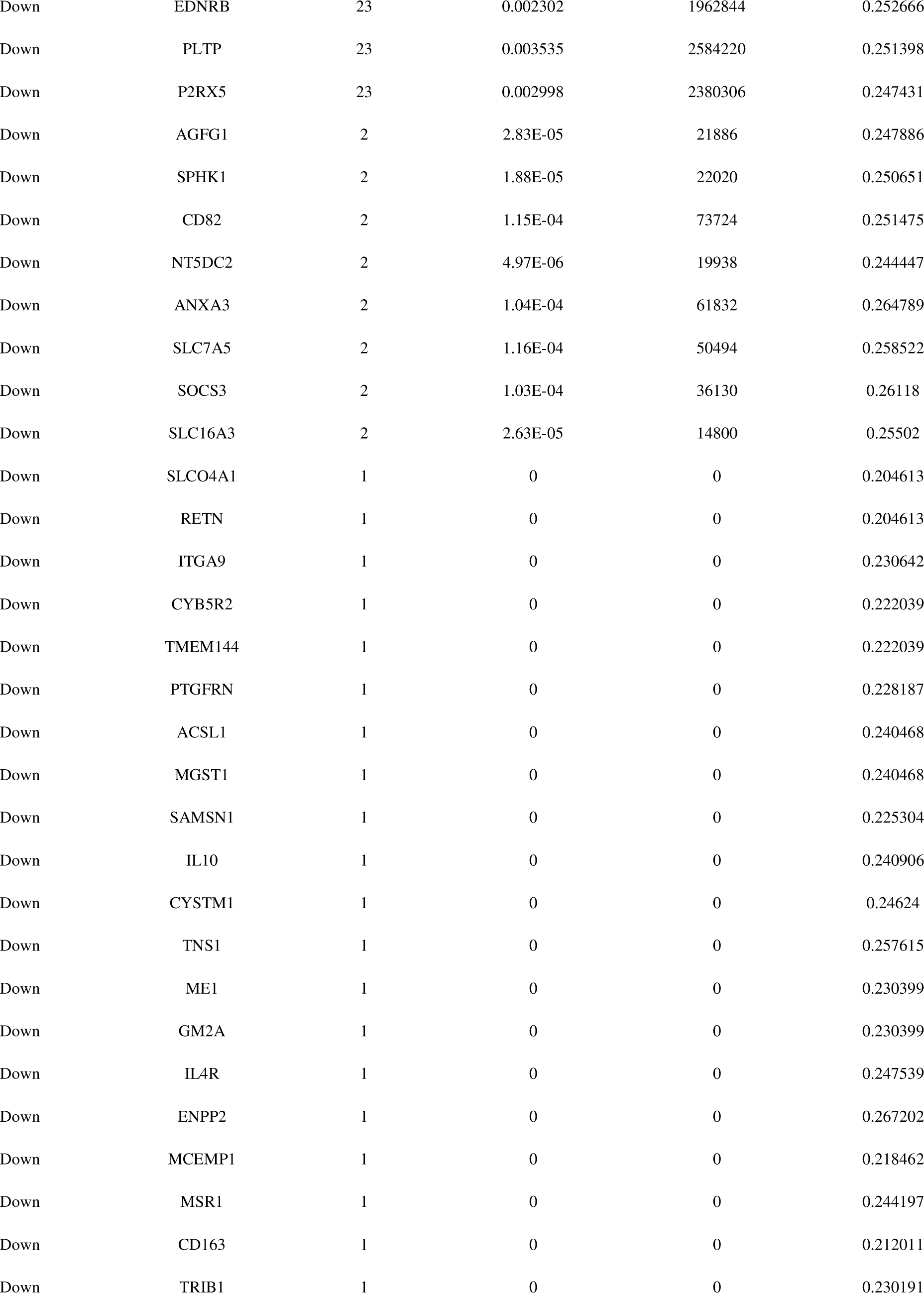
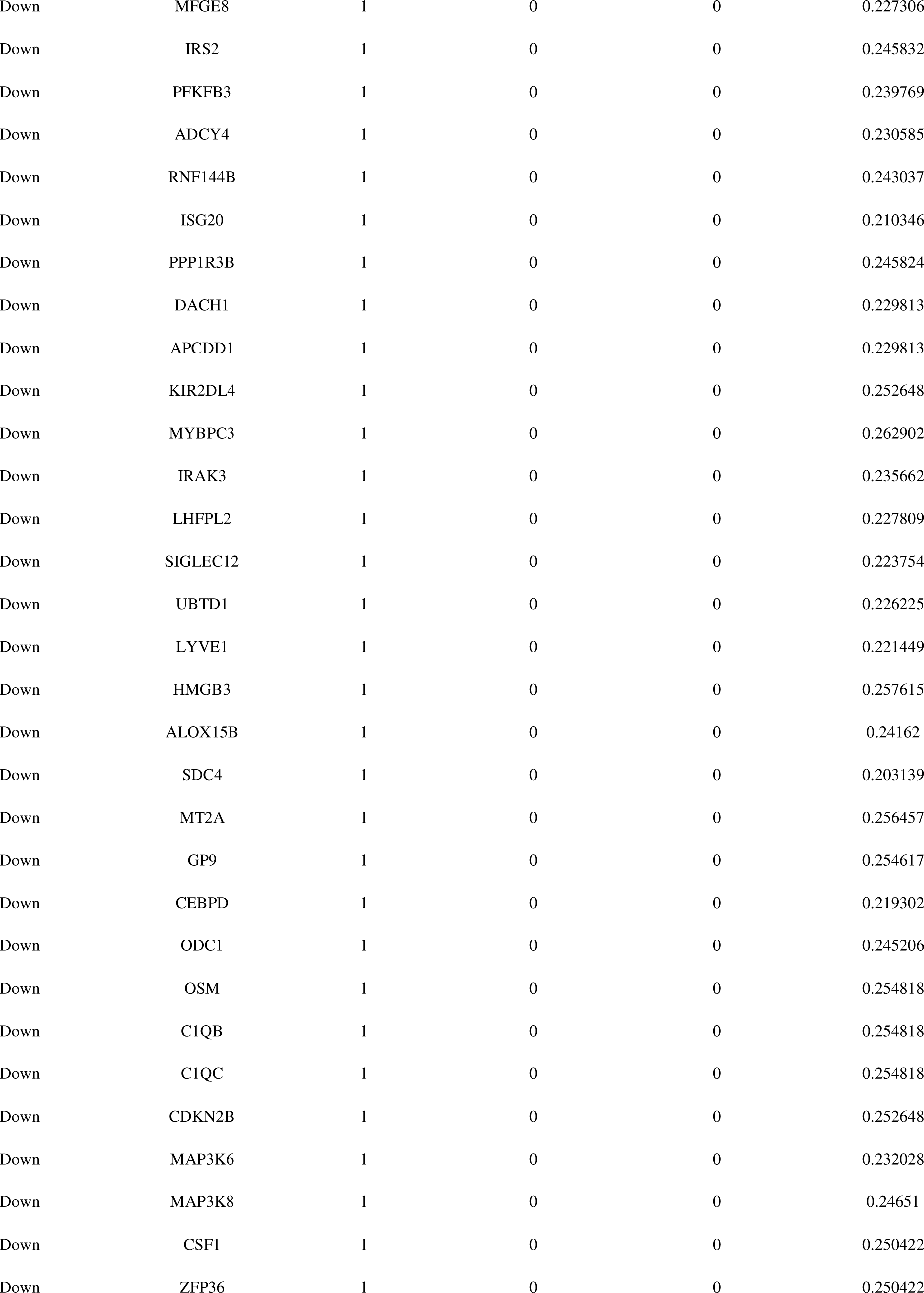
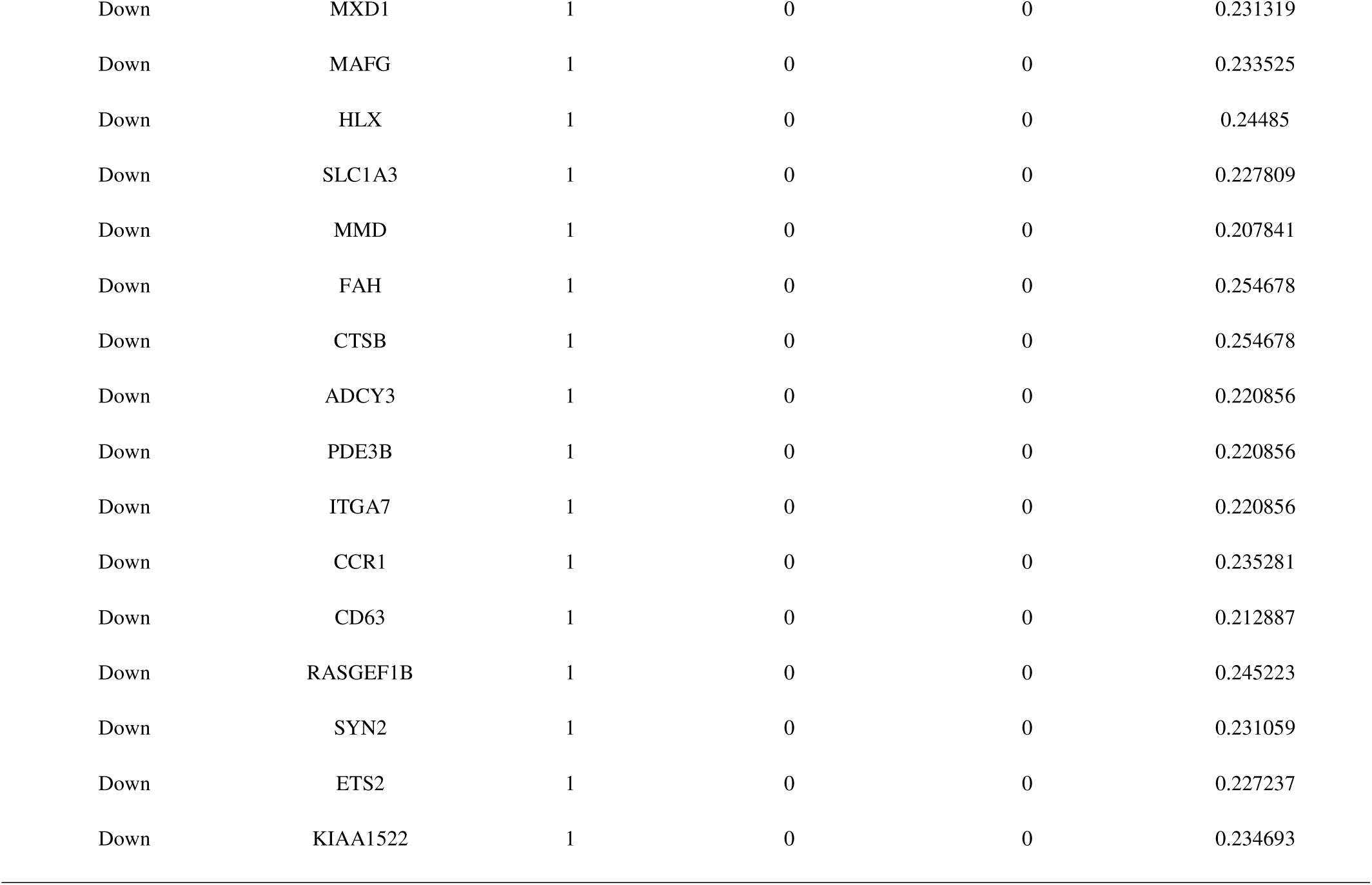
Topology table for up and down regulated genes.

| Regulation | Node | Degree | Betweenness | Stress | Closeness |
| --- | --- | --- | --- | --- | --- |
| Up | CUL1 | 423 | 0.086477 | 50906436 | 0.356655 |
| Up | HSPA8 | 333 | 0.075235 | 49599932 | 0.352385 |
| Up | HOXA1 | 271 | 0.042702 | 21619954 | 0.341577 |
| Up | INCA1 | 251 | 0.046392 | 24172002 | 0.338043 |
| Up | TP53 | 221 | 0.039224 | 23811026 | 0.336677 |
| Up | FBL | 214 | 0.033705 | 20256826 | 0.329835 |
| Up | COQ9 | 185 | 0.034653 | 15748268 | 0.316587 |
| Up | C1QBP | 182 | 0.033307 | 18453144 | 0.341939 |
| Up | ERBB2 | 181 | 0.033332 | 16956936 | 0.328957 |
| Up | PIN1 | 167 | 0.028195 | 13831878 | 0.327144 |
| Up | FHL3 | 166 | 0.032639 | 13478850 | 0.334069 |
| Up | SH2D3C | 157 | 0.02838 | 11054430 | 0.334476 |
| Up | CCT7 | 148 | 0.02378 | 14824174 | 0.32618 |
| Up | MRPL12 | 145 | 0.023984 | 11950664 | 0.332422 |
| Up | MRPL58 | 144 | 0.017244 | 9619288 | 0.309545 |
| Up | PPP1CA | 125 | 0.019222 | 12142406 | 0.325936 |
| Up | EEF1A1 | 125 | 0.01788 | 10431680 | 0.324837 |
| Up | YWHAQ | 121 | 0.018632 | 9890120 | 0.32595 |
| Up | FUS | 107 | 0.012518 | 8720394 | 0.331605 |
| Up | GLRX3 | 103 | 0.012859 | 6167724 | 0.317346 |
| Up | PPP1R18 | 98 | 0.008521 | 8057994 | 0.315429 |
| Up | MRPL4 | 97 | 0.010985 | 5327918 | 0.311299 |
| Up | PHB2 | 88 | 0.013293 | 7141400 | 0.3215 |
| Up | AP1M1 | 86 | 0.01094 | 6624796 | 0.308617 |
| Up | LENG1 | 84 | 0.007276 | 5732502 | 0.307657 |
| Up | TRIB3 | 83 | 0.009058 | 4690290 | 0.311534 |
| Up | TMEM128 | 77 | 0.011134 | 6869134 | 0.295004 |
| Up | PEBP1 | 77 | 0.010938 | 4225950 | 0.315375 |
| Up | RBCK1 | 77 | 0.011372 | 4658634 | 0.320195 |
| Up | RCN1 | 76 | 0.009445 | 6481560 | 0.32451 |
| Up | PKN1 | 75 | 0.007813 | 5528324 | 0.302438 |
| Up | COPS3 | 73 | 0.009818 | 4174738 | 0.325593 |
| Up | DKC1 | 72 | 0.005158 | 4173080 | 0.301174 |
| Up | CEP131 | 72 | 0.009893 | 4593256 | 0.30669 |
| Up | BRK1 | 70 | 0.007441 | 4221726 | 0.319988 |
| Up | CDK4 | 70 | 0.010963 | 4856726 | 0.314773 |
| Up | DNAAF2 | 70 | 0.011467 | 4216900 | 0.300478 |
| Up | RPL10A | 67 | 0.006818 | 5144592 | 0.31633 |
| Up | TMEM80 | 66 | 0.007973 | 5525744 | 0.270457 |
| Up | PPM1G | 65 | 0.005972 | 4487304 | 0.299774 |
| Up | TRAP1 | 65 | 0.006534 | 3721874 | 0.312967 |
| Up | PLGRKT | 65 | 0.007205 | 3484026 | 0.298123 |
| Up | MRPS35 | 64 | 0.003456 | 3327572 | 0.295922 |
| Up | ATXN10 | 62 | 0.00669 | 2912332 | 0.319148 |
| Up | TRIM21 | 61 | 0.007409 | 3220248 | 0.318586 |
| Up | DDX54 | 60 | 0.005297 | 3670220 | 0.31436 |
| Up | CISD3 | 60 | 0.004624 | 3090524 | 0.287358 |
| Up | TSPAN2 | 59 | 0.006346 | 5984490 | 0.295639 |
| Up | IARS2 | 59 | 0.008482 | 3193068 | 0.312901 |
| Up | TRAF3IP3 | 58 | 0.008316 | 3709276 | 0.305176 |
| Up | CYB561 | 58 | 0.006467 | 3663986 | 0.284471 |
| Up | MRPS26 | 58 | 0.004418 | 2551264 | 0.314934 |
| Up | TSSC4 | 58 | 0.006457 | 4430528 | 0.303528 |
| Up | TMEM179B | 58 | 0.007426 | 5185952 | 0.265386 |
| Up | MCM2 | 58 | 0.007524 | 4031136 | 0.29786 |
| Up | PIAS4 | 57 | 0.005891 | 3911814 | 0.295981 |
| Up | SLC25A11 | 57 | 0.004994 | 3622148 | 0.313112 |
| Up | MRPS7 | 56 | 0.004024 | 2951928 | 0.301688 |
| Up | UBE2D4 | 56 | 0.007418 | 3384822 | 0.302032 |
| Up | RPGRIP1 | 56 | 0.007381 | 4006096 | 0.291411 |
| Up | MRPL11 | 55 | 0.004981 | 3297568 | 0.309959 |
| Up | PLPP6 | 55 | 0.007038 | 3936626 | 0.28824 |
| Up | SNAP29 | 54 | 0.007492 | 3491826 | 0.311691 |
| Up | RNF20 | 54 | 0.005688 | 2638066 | 0.305251 |
| Up | BAIAP2 | 54 | 0.004583 | 4482396 | 0.306652 |
| Up | GPN3 | 52 | 0.006339 | 2748984 | 0.295004 |
| Up | RNF5 | 51 | 0.006282 | 2604726 | 0.292353 |
| Up | FAM50B | 50 | 0.003085 | 3902212 | 0.283774 |
| Up | FCHO1 | 49 | 0.003517 | 3310928 | 0.285324 |
| Up | TMEM42 | 49 | 0.004216 | 5437438 | 0.262363 |
| Up | COMTD1 | 48 | 0.007252 | 3466690 | 0.298027 |
| Up | RAB8A | 48 | 0.005235 | 3485368 | 0.288465 |
| Up | HSPBP1 | 47 | 0.003009 | 1770352 | 0.301149 |
| Up | NUP93 | 47 | 0.004501 | 3642972 | 0.306298 |
| Up | IL17RC | 45 | 0.009416 | 3011222 | 0.283449 |
| Up | PPP5C | 45 | 0.00393 | 2909678 | 0.289387 |
| Up | ZNF696 | 45 | 0.00273 | 2815170 | 0.286083 |
| Up | TTC30B | 44 | 0.007069 | 3220442 | 0.282822 |
| Up | CKB | 42 | 0.00489 | 2565088 | 0.319492 |
| Up | TRAPPC6A | 41 | 0.003712 | 1540304 | 0.293035 |
| Up | EIF2B1 | 41 | 0.005233 | 2401664 | 0.293104 |
| Up | NACA | 40 | 0.002865 | 2084190 | 0.313165 |
| Up | HIBADH | 40 | 0.004009 | 3042574 | 0.268256 |
| Up | ZNF574 | 40 | 0.002805 | 2130412 | 0.296751 |
| Up | VPS51 | 39 | 0.004079 | 2116788 | 0.294045 |
| Up | CYBRD1 | 39 | 0.003235 | 2771472 | 0.286028 |
| Up | RNF126 | 39 | 0.004026 | 1800794 | 0.293452 |
| Up | TMEM254 | 39 | 0.001912 | 2101420 | 0.262103 |
| Up | PCGF2 | 38 | 0.004886 | 1748478 | 0.301235 |
| Up | TRIM68 | 38 | 0.005267 | 1776624 | 0.29084 |
| Up | SURF6 | 37 | 0.001578 | 1628498 | 0.292572 |
| Up | RNF114 | 36 | 0.003552 | 2304736 | 0.2847 |
| Up | CORO1A | 36 | 0.004226 | 1601750 | 0.298615 |
| Up | SNRNP40 | 36 | 0.002742 | 2210918 | 0.294419 |
| Up | SNRPD3 | 35 | 0.002364 | 2014372 | 0.304387 |
| Up | DYRK1B | 35 | 0.004935 | 3051100 | 0.268081 |
| Up | NDUFA8 | 34 | 0.002703 | 1879294 | 0.299822 |
| Up | NGRN | 34 | 0.001899 | 1237088 | 0.27025 |
| Up | ECHS1 | 33 | 0.002133 | 1324802 | 0.292399 |
| Up | MRPL45 | 33 | 0.001955 | 1157310 | 0.295852 |
| Up | CEACAM21 | 33 | 0.003192 | 2310528 | 0.249739 |
| Up | LRWD1 | 33 | 0.004498 | 2459018 | 0.28494 |
| Up | ALG8 | 33 | 0.003058 | 2781576 | 0.259917 |
| Up | PFDN6 | 32 | 0.00228 | 984618 | 0.285401 |
| Up | UBTF | 32 | 0.002712 | 1814518 | 0.29351 |
| Up | EIF2B3 | 32 | 0.003474 | 3064980 | 0.271685 |
| Up | ZNHIT2 | 32 | 0.002405 | 1009178 | 0.285039 |
| Up | BZW2 | 32 | 0.003767 | 1633914 | 0.276156 |
| Up | GGT7 | 31 | 0.004036 | 2718942 | 0.249236 |
| Up | RNASEH2A | 31 | 0.004644 | 1876060 | 0.269915 |
| Up | APBA3 | 31 | 0.003957 | 2676702 | 0.262716 |
| Up | RPS6KA1 | 31 | 0.003014 | 1195156 | 0.300917 |
| Up | LAS1L | 31 | 0.001811 | 1573844 | 0.30148 |
| Up | RNASEH2B | 31 | 0.004112 | 1545740 | 0.279665 |
| Up | NDUFA12 | 31 | 0.00424 | 2244700 | 0.295251 |
| Up | CAPRIN1 | 31 | 0.002321 | 1395300 | 0.300429 |
| Up | GPR25 | 30 | 0.003162 | 2376166 | 0.25724 |
| Up | TUBB4A | 30 | 0.001593 | 1627240 | 0.279623 |
| Up | FIS1 | 30 | 0.00337 | 1980666 | 0.271755 |
| Up | RHOT2 | 30 | 0.003039 | 1963586 | 0.304775 |
| Up | NUDT16L1 | 4 | 7.90E-04 | 506674 | 0.283038 |
| Up | POLD2 | 3 | 2.26E-04 | 121380 | 0.258657 |
| Up | RASSF7 | 3 | 7.41E-05 | 66966 | 0.253607 |
| Up | DOLK | 3 | 9.75E-05 | 90036 | 0.226142 |
| Up | RPUSD4 | 3 | 1.56E-05 | 38938 | 0.258775 |
| Up | RPUSD3 | 3 | 0 | 0 | 0.246995 |
| Up | SMG8 | 2 | 3.61E-05 | 15014 | 0.251851 |
| Up | CYB561A3 | 2 | 1.16E-05 | 16546 | 0.235005 |
| Up | TMEM109 | 2 | 5.40E-06 | 12066 | 0.219204 |
| Up | TMEM230 | 2 | 1.20E-04 | 119670 | 0.267106 |
| Up | NEFH | 2 | 1.04E-04 | 120714 | 0.27102 |
| Up | NSMCE4A | 2 | 1.08E-04 | 121656 | 0.268246 |
| Up | SUMF2 | 2 | 5.14E-05 | 83822 | 0.26811 |
| Up | MRPS33 | 2 | 9.32E-05 | 41250 | 0.260045 |
| Up | NOA1 | 2 | 3.05E-04 | 116110 | 0.266962 |
| Up | ARHGEF2 | 2 | 6.38E-05 | 77418 | 0.268013 |
| Up | UTP6 | 2 | 4.00E-05 | 14286 | 0.25276 |
| Up | C7orf50 | 2 | 5.62E-05 | 100046 | 0.271994 |
| Up | STK38 | 2 | 4.84E-05 | 27812 | 0.261134 |
| Up | DNAJC1 | 2 | 4.39E-05 | 10294 | 0.244358 |
| Up | ZBTB3 | 2 | 3.31E-05 | 10150 | 0.24464 |
| Up | LYSMD2 | 2 | 4.54E-05 | 12914 | 0.242481 |
| Up | LYRM4 | 2 | 7.34E-05 | 32440 | 0.262437 |
| Up | EIF3L | 2 | 6.27E-05 | 58998 | 0.267858 |
| Up | ARV1 | 2 | 5.24E-05 | 59196 | 0.231977 |
| Up | ABHD14A | 2 | 5.00E-05 | 31154 | 0.213157 |
| Up | DGCR6L | 2 | 5.10E-06 | 9428 | 0.257302 |
| Up | CYB561D2 | 2 | 3.85E-05 | 51814 | 0.242465 |
| Up | RRP9 | 2 | 5.94E-05 | 38018 | 0.266789 |
| Up | S1PR2 | 2 | 6.30E-05 | 24902 | 0.234418 |
| Up | EEF1B2 | 2 | 6.30E-05 | 55312 | 0.269955 |
| Up | IDH2 | 2 | 0 | 0 | 0.254539 |
| Up | SUCLG1 | 2 | 5.72E-05 | 54116 | 0.258693 |
| Up | FXN | 2 | 1.84E-05 | 10768 | 0.233034 |
| Up | SYNGR1 | 2 | 3.40E-06 | 8812 | 0.216473 |
| Up | CCDC12 | 2 | 3.05E-06 | 9598 | 0.240266 |
| Up | PCED1A | 1 | 0 | 0 | 0.24214 |
| Up | COG2 | 1 | 0 | 0 | 0.212552 |
| Up | RHOBTB2 | 1 | 0 | 0 | 0.247539 |
| Up | ANXA6 | 1 | 0 | 0 | 0.247539 |
| Up | TMEM134 | 1 | 0 | 0 | 0.247539 |
| Up | INPP5K | 1 | 0 | 0 | 0.267202 |
| Up | ZNF641 | 1 | 0 | 0 | 0.267202 |
| Up | MIPEP | 1 | 0 | 0 | 0.223222 |
| Up | BCL2L12 | 1 | 0 | 0 | 0.244197 |
| Up | INTS5 | 1 | 0 | 0 | 0.241675 |
| Up | SNX20 | 1 | 0 | 0 | 0.260575 |
| Up | CNN2 | 1 | 0 | 0 | 0.260575 |
| Up | TYSND1 | 1 | 0 | 0 | 0.260575 |
| Up | VTA1 | 1 | 0 | 0 | 0.23942 |
| Up | COMMD6 | 1 | 0 | 0 | 0.239614 |
| Up | CCDC86 | 1 | 0 | 0 | 0.248035 |
| Up | TRIM4 | 1 | 0 | 0 | 0.210926 |
| Up | CCDC77 | 1 | 0 | 0 | 0.22529 |
| Up | PREP | 1 | 0 | 0 | 0.256457 |
| Up | ZNF296 | 1 | 0 | 0 | 0.256457 |
| Up | TBC1D10C | 1 | 0 | 0 | 0.254617 |
| Up | MRM2 | 1 | 0 | 0 | 0.249495 |
| Up | PAQR8 | 1 | 0 | 0 | 0.24326 |
| Up | CTNBNB1 | 1 | 0 | 0 | 0.215742 |
| Up | RAD9A | 1 | 0 | 0 | 0.220403 |
| Up | FEM1A | 1 | 0 | 0 | 0.252648 |
| Up | NEIL2 | 1 | 0 | 0 | 0.252648 |
| Up | OLFM1 | 1 | 0 | 0 | 0.244318 |
| Up | FAM162A | 1 | 0 | 0 | 0.244318 |
| Up | PTRH1 | 1 | 0 | 0 | 0.245012 |
| Up | ABCB9 | 1 | 0 | 0 | 0.22529 |
| Up | HINT2 | 1 | 0 | 0 | 0.250651 |
| Up | APOBEC3C | 1 | 0 | 0 | 0.237542 |
| Up | AK5 | 1 | 0 | 0 | 0.221615 |
| Up | ICMT | 1 | 0 | 0 | 0.216922 |
| Up | CD37 | 1 | 0 | 0 | 0.217827 |
| Up | NUDT7 | 1 | 0 | 0 | 0.262902 |
| Up | QDPR | 1 | 0 | 0 | 0.262902 |
| Up | NDUFB5 | 1 | 0 | 0 | 0.22529 |
| Up | RWDD2B | 1 | 0 | 0 | 0.241943 |
| Up | CRIP1 | 1 | 0 | 0 | 0.250651 |
| Up | WDR24 | 1 | 0 | 0 | 0.245963 |
| Up | SCAP | 1 | 0 | 0 | 0.226225 |
| Up | ZNF19 | 1 | 0 | 0 | 0.235842 |
| Up | RP9 | 1 | 0 | 0 | 0.235842 |
| Up | MRPL48 | 1 | 0 | 0 | 0.237405 |
| Up | MRPL34 | 1 | 0 | 0 | 0.237405 |
| Up | PALLD | 1 | 0 | 0 | 0.245223 |
| Up | SPN | 1 | 0 | 0 | 0.213648 |
| Up | RAB11FIP1 | 1 | 0 | 0 | 0.234693 |
| Up | PIGP | 1 | 0 | 0 | 0.213691 |
| Up | WDR18 | 1 | 0 | 0 | 0.231651 |
| Down | HSPB1 | 486 | 0.10954 | 81190764 | 0.364615 |
| Down | LMNA | 313 | 0.061499 | 39266360 | 0.346994 |
| Down | SNCA | 228 | 0.041238 | 19229464 | 0.344897 |
| Down | ADAMTSL4 | 219 | 0.034949 | 18832666 | 0.341687 |
| Down | PDLIM7 | 141 | 0.022651 | 11907684 | 0.32488 |
| Down | BAG3 | 122 | 0.018881 | 9282676 | 0.326668 |
| Down | ASPH | 114 | 0.020559 | 9556564 | 0.323294 |
| Down | PLSCR1 | 109 | 0.016481 | 7713250 | 0.319492 |
| Down | FFAR2 | 108 | 0.016584 | 9347300 | 0.285401 |
| Down | GRB10 | 103 | 0.011828 | 6634834 | 0.329703 |
| Down | IFITM3 | 99 | 0.015383 | 9831002 | 0.311442 |
| Down | HSPA1A | 98 | 0.009727 | 8645306 | 0.323082 |
| Down | LONRF1 | 95 | 0.01267 | 5537396 | 0.324226 |
| Down | BCL6 | 95 | 0.015355 | 6108936 | 0.311966 |
| Down | MAP2K1 | 93 | 0.015004 | 6434382 | 0.324851 |
| Down | C3AR1 | 91 | 0.013752 | 6774804 | 0.290794 |
| Down | P4HA2 | 87 | 0.013383 | 6178548 | 0.313828 |
| Down | CDKN1A | 87 | 0.01169 | 6905224 | 0.321054 |
| Down | SIPA1L1 | 85 | 0.0094 | 6541960 | 0.302685 |
| Down | CKAP4 | 85 | 0.013312 | 9899766 | 0.31396 |
| Down | FKBP5 | 82 | 0.010666 | 5323690 | 0.302118 |
| Down | HIF1A | 80 | 0.011308 | 5161632 | 0.304662 |
| Down | KLHL2 | 76 | 0.009871 | 4706808 | 0.297179 |
| Down | FPR2 | 76 | 0.010741 | 11772998 | 0.267299 |
| Down | SDC2 | 73 | 0.012646 | 5428852 | 0.306805 |
| Down | PHC2 | 72 | 0.009262 | 4860576 | 0.319148 |
| Down | CLU | 70 | 0.009994 | 4514104 | 0.318682 |
| Down | PIK3R1 | 70 | 0.012364 | 4303280 | 0.327577 |
| Down | HSPA6 | 68 | 0.006271 | 4930836 | 0.315107 |
| Down | PKM | 68 | 0.008434 | 5881790 | 0.325421 |
| Down | FTH1 | 67 | 0.011027 | 4315964 | 0.299362 |
| Down | TUBB2A | 63 | 0.005786 | 4762308 | 0.313814 |
| Down | CDH5 | 63 | 0.007687 | 3126456 | 0.295569 |
| Down | ASGR2 | 61 | 0.007656 | 5708950 | 0.279518 |
| Down | EMP1 | 60 | 0.007526 | 5687858 | 0.278479 |
| Down | MPP1 | 58 | 0.009665 | 3967034 | 0.321444 |
| Down | GABARAPL1 | 58 | 0.00706 | 2634158 | 0.309067 |
| Down | TMEM132A | 58 | 0.008365 | 3425776 | 0.29484 |
| Down | PER1 | 56 | 0.00652 | 3966082 | 0.318887 |
| Down | ECM1 | 56 | 0.007496 | 3440732 | 0.308309 |
| Down | FAM110B | 53 | 0.002516 | 2106046 | 0.273761 |
| Down | FOSL2 | 53 | 0.007406 | 4469782 | 0.299012 |
| Down | UBE2J1 | 53 | 0.006719 | 4636514 | 0.284427 |
| Down | CPNE2 | 51 | 0.005523 | 4162406 | 0.287025 |
| Down | HMOX1 | 50 | 0.005628 | 3280364 | 0.29278 |
| Down | HSD3B7 | 49 | 0.00375 | 2184880 | 0.301198 |
| Down | TNFAIP3 | 49 | 0.006183 | 2762590 | 0.294466 |
| Down | PRKAR2B | 49 | 0.00703 | 2307444 | 0.290794 |
| Down | SOCS1 | 48 | 0.004753 | 2508986 | 0.310933 |
| Down | ZNF497 | 48 | 0.003199 | 2328958 | 0.299822 |
| Down | TGM2 | 48 | 0.004404 | 2218894 | 0.318094 |
| Down | PRKCE | 48 | 0.004611 | 4613286 | 0.282703 |
| Down | ERN1 | 47 | 0.00567 | 2390980 | 0.300089 |
| Down | BRD1 | 47 | 0.005088 | 2397424 | 0.309093 |
| Down | HBA2 | 46 | 0.006469 | 2476590 | 0.311351 |
| Down | TSPYL2 | 46 | 0.004807 | 3168278 | 0.281962 |
| Down | ACTN1 | 45 | 0.004252 | 1889986 | 0.30881 |
| Down | MBOAT7 | 45 | 0.005489 | 4050068 | 0.282499 |
| Down | NECTIN2 | 44 | 0.002917 | 1848552 | 0.299653 |
| Down | DYSF | 43 | 0.005488 | 2939534 | 0.292607 |
| Down | JAK3 | 43 | 0.004085 | 1950768 | 0.296478 |
| Down | WDFY3 | 43 | 0.005439 | 2778900 | 0.308386 |
| Down | ELL2 | 42 | 0.004893 | 2287186 | 0.296206 |
| Down | ITGB5 | 41 | 0.003685 | 2100262 | 0.294162 |
| Down | PPARG | 41 | 0.005409 | 2274256 | 0.299677 |
| Down | LMNB1 | 40 | 0.003817 | 3353622 | 0.291125 |
| Down | MS4A4A | 40 | 0.004637 | 2429258 | 0.289184 |
| Down | TMEM45B | 40 | 0.00256 | 1962494 | 0.254696 |
| Down | VSIG4 | 39 | 0.005092 | 1674776 | 0.282725 |
| Down | GNAQ | 39 | 0.005346 | 2375778 | 0.266367 |
| Down | BRCA2 | 38 | 0.004364 | 2683122 | 0.277769 |
| Down | S100A8 | 38 | 0.00324 | 1378136 | 0.300612 |
| Down | HBB | 38 | 0.005168 | 1840244 | 0.298375 |
| Down | IL1R2 | 38 | 0.004747 | 2378406 | 0.272784 |
| Down | CDC20 | 38 | 0.003425 | 1524044 | 0.309055 |
| Down | GOLIM4 | 37 | 0.003743 | 2525050 | 0.282477 |
| Down | TLE3 | 35 | 0.00327 | 2720548 | 0.273599 |
| Down | S100A9 | 34 | 0.002517 | 2311014 | 0.283948 |
| Down | CTSD | 34 | 0.003607 | 3491288 | 0.283535 |
| Down | SMAD1 | 33 | 0.002994 | 1241088 | 0.285654 |
| Down | TUBB6 | 33 | 0.001678 | 1441264 | 0.316628 |
| Down | RAPGEF2 | 33 | 0.00331 | 1464564 | 0.292722 |
| Down | FCGR1A | 33 | 0.004188 | 2045558 | 0.269044 |
| Down | SH3RF3 | 32 | 7.16E-04 | 681562 | 0.272203 |
| Down | BEX3 | 32 | 0.002565 | 2666716 | 0.275081 |
| Down | AVPI1 | 32 | 0.001606 | 1490490 | 0.28494 |
| Down | FTL | 31 | 0.002598 | 1240626 | 0.29436 |
| Down | GK | 31 | 0.004237 | 1920014 | 0.268644 |
| Down | ID1 | 31 | 0.003432 | 2001784 | 0.282251 |
| Down | INF2 | 31 | 0.001678 | 1103472 | 0.285819 |
| Down | GPRC5C | 31 | 0.003033 | 2798386 | 0.273801 |
| Down | SAP30 | 30 | 0.00414 | 1507732 | 0.290817 |
| Down | GPR183 | 30 | 0.004795 | 1735682 | 0.254915 |
| Down | SMAP2 | 30 | 0.002874 | 1171458 | 0.284024 |
| Down | LIMK2 | 29 | 0.003567 | 1847760 | 0.309661 |
| Down | JOSD1 | 29 | 6.74E-04 | 561132 | 0.279887 |
| Down | CORO2A | 29 | 0.001451 | 1527612 | 0.266923 |
| Down | SLC7A8 | 28 | 0.002374 | 1855794 | 0.277894 |
| Down | TSC22D3 | 28 | 0.002933 | 1127212 | 0.294864 |
| Down | BASP1 | 28 | 0.001218 | 1131454 | 0.280894 |
| Down | SYAP1 | 28 | 0.003055 | 1601354 | 0.289275 |
| Down | SLC39A8 | 27 | 0.003306 | 2067316 | 0.263791 |
| Down | GLUL | 27 | 0.002776 | 2386452 | 0.268781 |
| Down | EGFL7 | 27 | 0.002068 | 767896 | 0.28718 |
| Down | CXCR4 | 27 | 0.002703 | 1327960 | 0.270506 |
| Down | BCAT1 | 27 | 0.002801 | 4165724 | 0.263126 |
| Down | STEAP3 | 27 | 0.002504 | 1154880 | 0.269729 |
| Down | TMTC1 | 26 | 0.001592 | 1195170 | 0.3017 |
| Down | FOSL1 | 26 | 0.0013 | 981190 | 0.292376 |
| Down | PLB1 | 26 | 0.00207 | 1617562 | 0.279023 |
| Down | BCL2L11 | 25 | 0.002561 | 2261982 | 0.255979 |
| Down | P2RX4 | 25 | 0.002668 | 1441648 | 0.269063 |
| Down | CREM | 25 | 0.002734 | 2255976 | 0.271655 |
| Down | GCA | 24 | 0.002452 | 1981766 | 0.277002 |
| Down | GAS2L1 | 24 | 0.002103 | 2563688 | 0.265035 |
| Down | AQP9 | 23 | 0.001404 | 1211632 | 0.248985 |
| Down | LPL | 23 | 0.001491 | 907406 | 0.284645 |
| Down | EDNRB | 23 | 0.002302 | 1962844 | 0.252666 |
| Down | PLTP | 23 | 0.003535 | 2584220 | 0.251398 |
| Down | P2RX5 | 23 | 0.002998 | 2380306 | 0.247431 |
| Down | AGFG1 | 2 | 2.83E-05 | 21886 | 0.247886 |
| Down | SPHK1 | 2 | 1.88E-05 | 22020 | 0.250651 |
| Down | CD82 | 2 | 1.15E-04 | 73724 | 0.251475 |
| Down | NT5DC2 | 2 | 4.97E-06 | 19938 | 0.244447 |
| Down | ANXA3 | 2 | 1.04E-04 | 61832 | 0.264789 |
| Down | SLC7A5 | 2 | 1.16E-04 | 50494 | 0.258522 |
| Down | SOC3 | 2 | 1.03E-04 | 36130 | 0.26118 |
| Down | SLC16A3 | 2 | 2.63E-05 | 14800 | 0.25502 |
| Down | SLCO4A1 | 1 | 0 | 0 | 0.204613 |
| Down | RETN | 1 | 0 | 0 | 0.204613 |
| Down | ITGA9 | 1 | 0 | 0 | 0.230642 |
| Down | CYB5R2 | 1 | 0 | 0 | 0.222039 |
| Down | TMEM144 | 1 | 0 | 0 | 0.222039 |
| Down | PTGFRN | 1 | 0 | 0 | 0.228187 |
| Down | ACSL1 | 1 | 0 | 0 | 0.240468 |
| Down | MGST1 | 1 | 0 | 0 | 0.240468 |
| Down | SAMSN1 | 1 | 0 | 0 | 0.225304 |
| Down | IL10 | 1 | 0 | 0 | 0.240906 |
| Down | CYSTM1 | 1 | 0 | 0 | 0.24624 |
| Down | TNS1 | 1 | 0 | 0 | 0.257615 |
| Down | ME1 | 1 | 0 | 0 | 0.230399 |
| Down | GM2A | 1 | 0 | 0 | 0.230399 |
| Down | IL4R | 1 | 0 | 0 | 0.247539 |
| Down | ENPP2 | 1 | 0 | 0 | 0.267202 |
| Down | MCEMP1 | 1 | 0 | 0 | 0.218462 |
| Down | MSR1 | 1 | 0 | 0 | 0.244197 |
| Down | CD163 | 1 | 0 | 0 | 0.212011 |
| Down | TRIB1 | 1 | 0 | 0 | 0.230191 |
| Down | MFGE8 | 1 | 0 | 0 | 0.227306 |
| Down | IRS2 | 1 | 0 | 0 | 0.245832 |
| Down | PFKFB3 | 1 | 0 | 0 | 0.239769 |
| Down | ADCY4 | 1 | 0 | 0 | 0.230585 |
| Down | RNF144B | 1 | 0 | 0 | 0.243037 |
| Down | ISG20 | 1 | 0 | 0 | 0.210346 |
| Down | PPP1R3B | 1 | 0 | 0 | 0.245824 |
| Down | DACH1 | 1 | 0 | 0 | 0.229813 |
| Down | APCDD1 | 1 | 0 | 0 | 0.229813 |
| Down | KIR2DL4 | 1 | 0 | 0 | 0.252648 |
| Down | MYBPC3 | 1 | 0 | 0 | 0.262902 |
| Down | IRAK3 | 1 | 0 | 0 | 0.235662 |
| Down | LHFPL2 | 1 | 0 | 0 | 0.227809 |
| Down | SIGLEC12 | 1 | 0 | 0 | 0.223754 |
| Down | UBTD1 | 1 | 0 | 0 | 0.226225 |
| Down | LYVE1 | 1 | 0 | 0 | 0.221449 |
| Down | HMGB3 | 1 | 0 | 0 | 0.257615 |
| Down | ALOX15B | 1 | 0 | 0 | 0.24162 |
| Down | SDC4 | 1 | 0 | 0 | 0.203139 |
| Down | MT2A | 1 | 0 | 0 | 0.256457 |
| Down | GP9 | 1 | 0 | 0 | 0.254617 |
| Down | CEBPD | 1 | 0 | 0 | 0.219302 |
| Down | ODC1 | 1 | 0 | 0 | 0.245206 |
| Down | OSM | 1 | 0 | 0 | 0.254818 |
| Down | C1QB | 1 | 0 | 0 | 0.254818 |
| Down | C1QC | 1 | 0 | 0 | 0.254818 |
| Down | CDKN2B | 1 | 0 | 0 | 0.252648 |
| Down | MAP3K6 | 1 | 0 | 0 | 0.232028 |
| Down | MAP3K8 | 1 | 0 | 0 | 0.24651 |
| Down | CSF1 | 1 | 0 | 0 | 0.250422 |
| Down | ZFP36 | 1 | 0 | 0 | 0.250422 |
| Down | MXD1 | 1 | 0 | 0 | 0.231319 |
| Down | MAFG | 1 | 0 | 0 | 0.233525 |
| Down | HLX | 1 | 0 | 0 | 0.24485 |
| Down | SLC1A3 | 1 | 0 | 0 | 0.227809 |
| Down | MMD | 1 | 0 | 0 | 0.207841 |
| Down | FAH | 1 | 0 | 0 | 0.254678 |
| Down | CTSB | 1 | 0 | 0 | 0.254678 |
| Down | ADCY3 | 1 | 0 | 0 | 0.220856 |
| Down | PDE3B | 1 | 0 | 0 | 0.220856 |
| Down | ITGA7 | 1 | 0 | 0 | 0.220856 |
| Down | CCR1 | 1 | 0 | 0 | 0.235281 |
| Down | CD63 | 1 | 0 | 0 | 0.212887 |
| Down | RASGEF1B | 1 | 0 | 0 | 0.245223 |
| Down | SYN2 | 1 | 0 | 0 | 0.231059 |
| Down | ETS2 | 1 | 0 | 0 | 0.227237 |
| Down | KIAA1522 | 1 | 0 | 0 | 0.234693 |

**Fig. 4.**
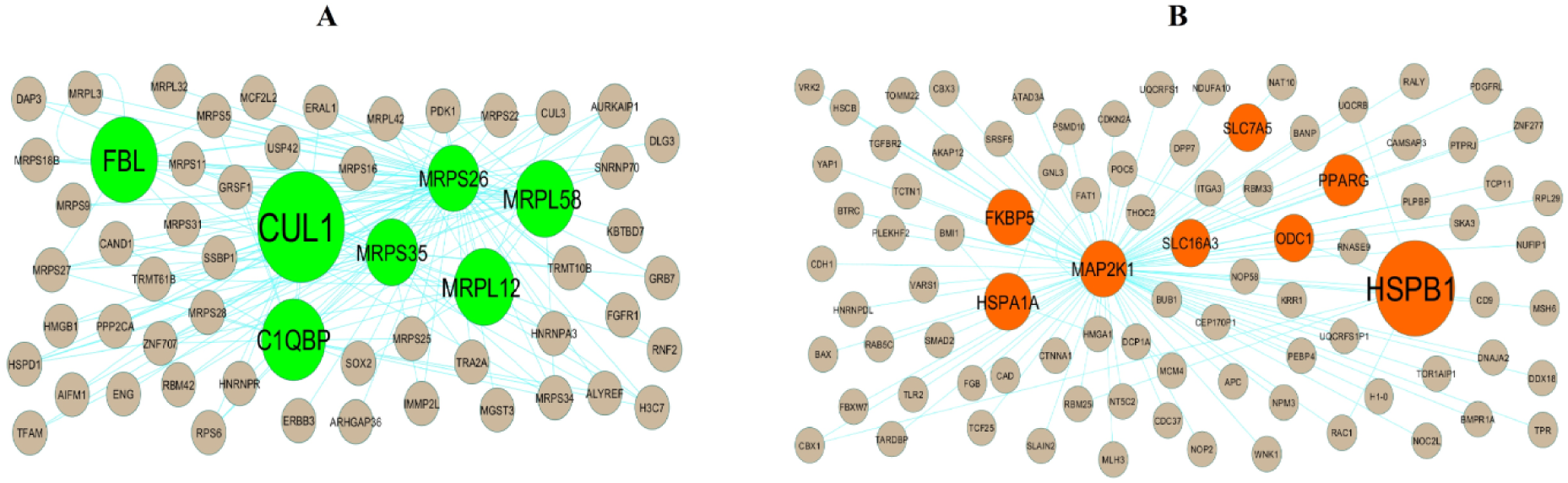
Modules selected from the PPI network. (A) The most significant module was obtained from PPI network with 58 nodes and 140 edges for up regulated genes (B) The most significant module was obtained from PPI network with 88 nodes and 97 edges for down regulated genes. Up regulated genes are marked in parrot green; down regulated genes are marked in red.

### Construction of the miRNA-hub gene regulatory network

A miRNA-hub gene regulatory network of the hub was constructed using the online website miRNet and software Cytoscape. The miRNA-hub gene regulatory network contained 5102 nodes, including 4700 miRNAs and 402 hub genes, and 61766 edges (Fig.5). 174 miRNAs (ex: hsa-mir-1914-5p) were forecasted as targets of HSPA8, 174 miRNAs (ex: hsa-mir-5197-5p) were forecasted as targets of TP53, 108 miRNAs (ex: hsa-mir-3664-3p) were forecasted as targets of MRPL12, 101 miRNAs (ex: hsa-mir-1260a) were forecasted as targets of ERBB2, 72 miRNAs (ex: hsa-mir-140-5p) were forecasted as targets of PIN1, 149 miRNAs (ex: hsa-mir-598-3p) were forecasted as targets of ASPH, 112 miRNAs (ex: hsa-mir-3609) were forecasted as targets of GRB10, 100 miRNAs (ex: hsa-mir-101-3p) were forecasted as targets of LONRF1, 94 miRNAs (ex: hsa-mir-449b-5p) were forecasted as targets of PLSCR1 and 84 miRNAs (ex: hsa-mir-369-3p) were forecasted as targets of MAP2K1 and are listed in Table 5. It can be speculated that the miRNAs regulates the advancement and progression of CA by activating or repressing the transcription of hub genes.

**Fig. 5.**
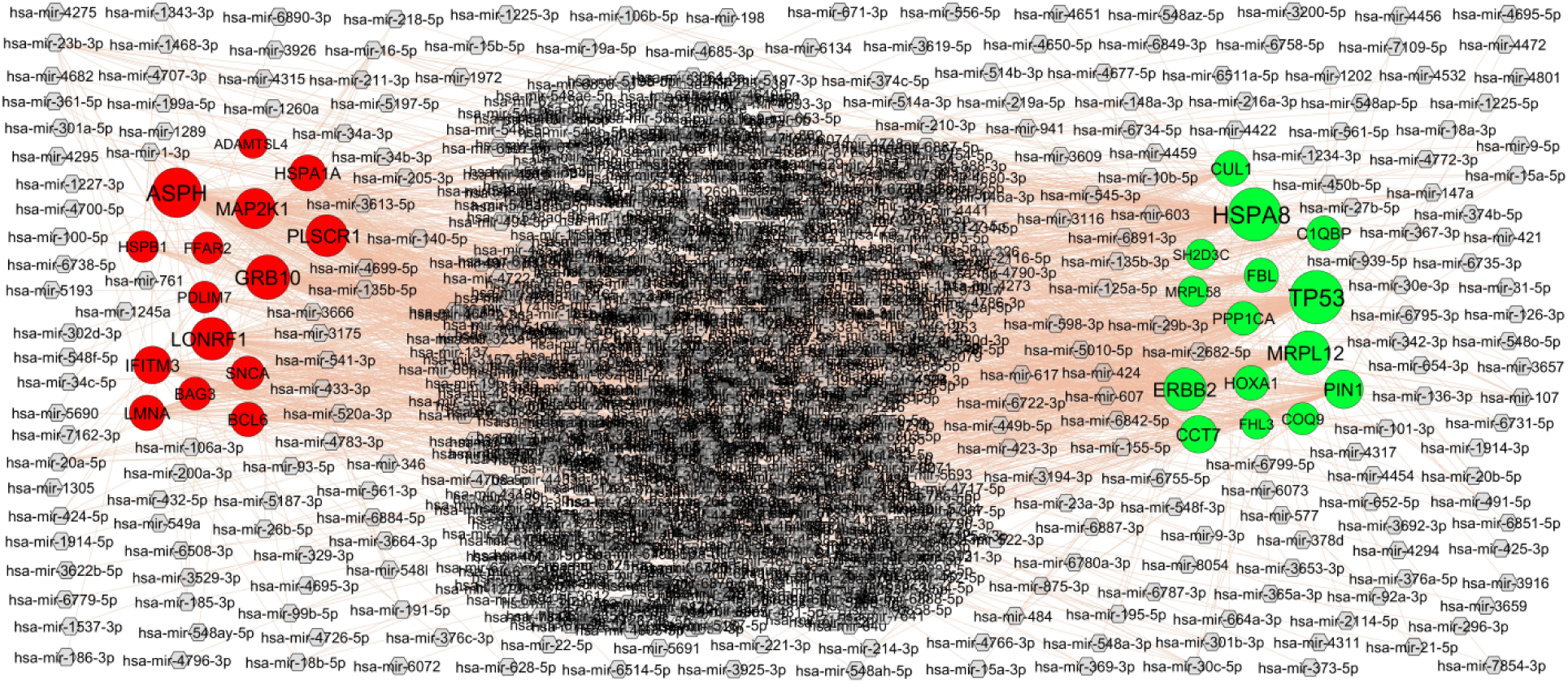
Hub gene - miRNA regulatory network. The ash color diamond nodes represent the key miRNAs; up regulated genes are marked in green; down regulated genes are marked in red.

**Table 5.**
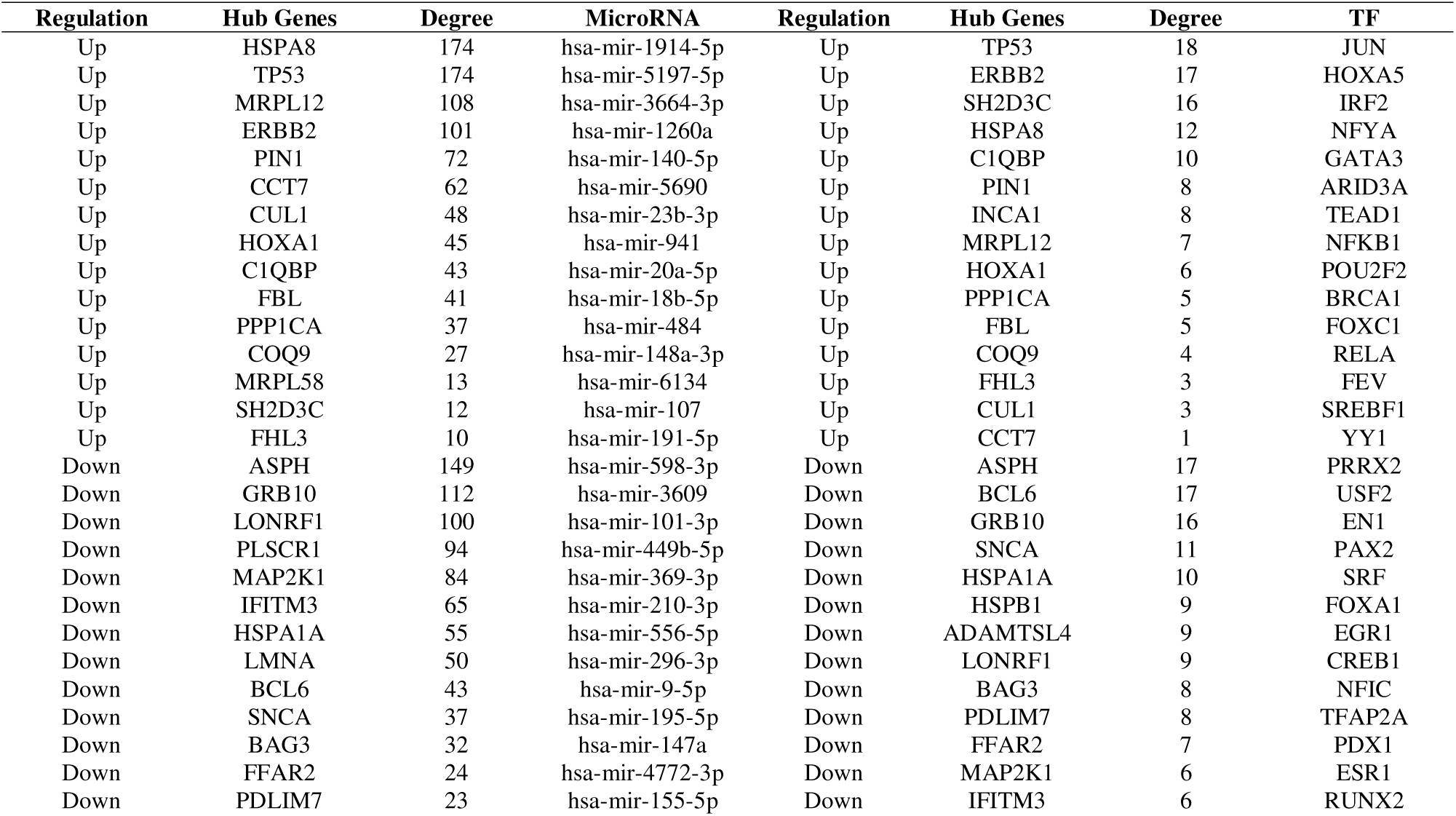
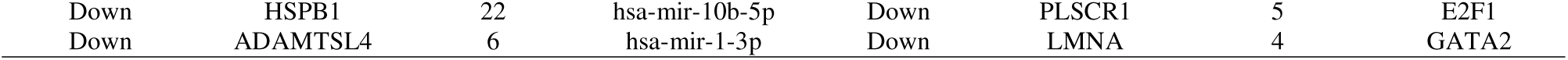
MiRNA - hub gene and TF – hub gene topology table.

| Regulation | Hub Genes | Degree | MicroRNA | Regulation | Hub Genes | Degree | TF |
| --- | --- | --- | --- | --- | --- | --- | --- |
| Up | HSPA8 | 174 | hsa-mir-1914-5p | Up | TP53 | 18 | JUN |
| Up | TP53 | 174 | hsa-mir-5197-5p | Up | ERBB2 | 17 | HOXA5 |
| Up | MRPL12 | 108 | hsa-mir-3664-3p | Up | SH2D3C | 16 | IRF2 |
| Up | ERBB2 | 101 | hsa-mir-1260a | Up | HSPA8 | 12 | NFYA |
| Up | PIN1 | 72 | hsa-mir-140-5p | Up | C1QBP | 10 | GATA3 |
| Up | CCT7 | 62 | hsa-mir-5690 | Up | PIN1 | 8 | ARID3A |
| Up | CUL1 | 48 | hsa-mir-23b-3p | Up | INCA1 | 8 | TEAD1 |
| Up | HOXA1 | 45 | hsa-mir-941 | Up | MRPL12 | 7 | NFKB1 |
| Up | C1QBP | 43 | hsa-mir-20a-5p | Up | HOXA1 | 6 | POU2F2 |
| Up | FBL | 41 | hsa-mir-18b-5p | Up | PPP1CA | 5 | BRCA1 |
| Up | PPP1CA | 37 | hsa-mir-484 | Up | FBL | 5 | FOXC1 |
| Up | COQ9 | 27 | hsa-mir-148a-3p | Up | COQ9 | 4 | RELA |
| Up | MRPL58 | 13 | hsa-mir-6134 | Up | FHL3 | 3 | FEV |
| Up | SH2D3C | 12 | hsa-mir-107 | Up | CUL1 | 3 | SREBF1 |
| Up | FHL3 | 10 | hsa-mir-191-5p | Up | CCT7 | 1 | YY1 |
| Down | ASPH | 149 | hsa-mir-598-3p | Down | ASPH | 17 | PRRX2 |
| Down | GRB10 | 112 | hsa-mir-3609 | Down | BCL6 | 17 | USF2 |
| Down | LONRF1 | 100 | hsa-mir-101-3p | Down | GRB10 | 16 | EN1 |
| Down | PLSCR1 | 94 | hsa-mir-449b-5p | Down | SNCA | 11 | PAX2 |
| Down | MAP2K1 | 84 | hsa-mir-369-3p | Down | HSPA1A | 10 | SRF |
| Down | IFITM3 | 65 | hsa-mir-210-3p | Down | HSPB1 | 9 | FOXA1 |
| Down | HSPA1A | 55 | hsa-mir-556-5p | Down | ADAMTSL4 | 9 | EGR1 |
| Down | LMNA | 50 | hsa-mir-296-3p | Down | LONRF1 | 9 | CREB1 |
| Down | BCL6 | 43 | hsa-mir-9-5p | Down | BAG3 | 8 | NFIC |
| Down | SNCA | 37 | hsa-mir-195-5p | Down | PDLIM7 | 8 | TFAP2A |
| Down | BAG3 | 32 | hsa-mir-147a | Down | FFAR2 | 7 | PDX1 |
| Down | FFAR2 | 24 | hsa-mir-4772-3p | Down | MAP2K1 | 6 | ESR1 |
| Down | PDLIM7 | 23 | hsa-mir-155-5p | Down | IFITM3 | 6 | RUNX2 |

### Construction of the TF-hub gene regulatory network

A TF-hub gene regulatory network of the hub was constructed using the online website NetworkAnalyst and software Cytoscape. The TF-hub gene regulatory network contained 482 nodes, including 86 TFs and 396 hub genes, and 3213 edges (Fig.6). 18 TFs (ex: JUN) were forecasted as targets of TP53, 17 TFs (ex: HOXA5) were forecasted as targets of ERBB2, 16 TFs (ex: IRF2) were forecasted as targets of SH2D3C, 12 TFs (ex: NFYA) were forecasted as targets of HSPA8, 10 TFs (ex: GATA3) were forecasted as targets of C1QBP, 17 TFs (ex: PRRX2) were forecasted as targets of ASPH, 17 TFs (ex: USF2) were forecasted as targets of BCL6, 16 TFs (ex: EN1) were forecasted as targets of GRB10, 11 TFs (ex: PAX2) were forecasted as targets of SNCA, and 10 TFs (ex: SRF) were forecasted as targets of HSPA1A, and are listed in Table 5. It can be speculated that the TFs regulates the advancement and progression of CA by activating or repressing the transcription of hub genes.

**Fig. 6.**
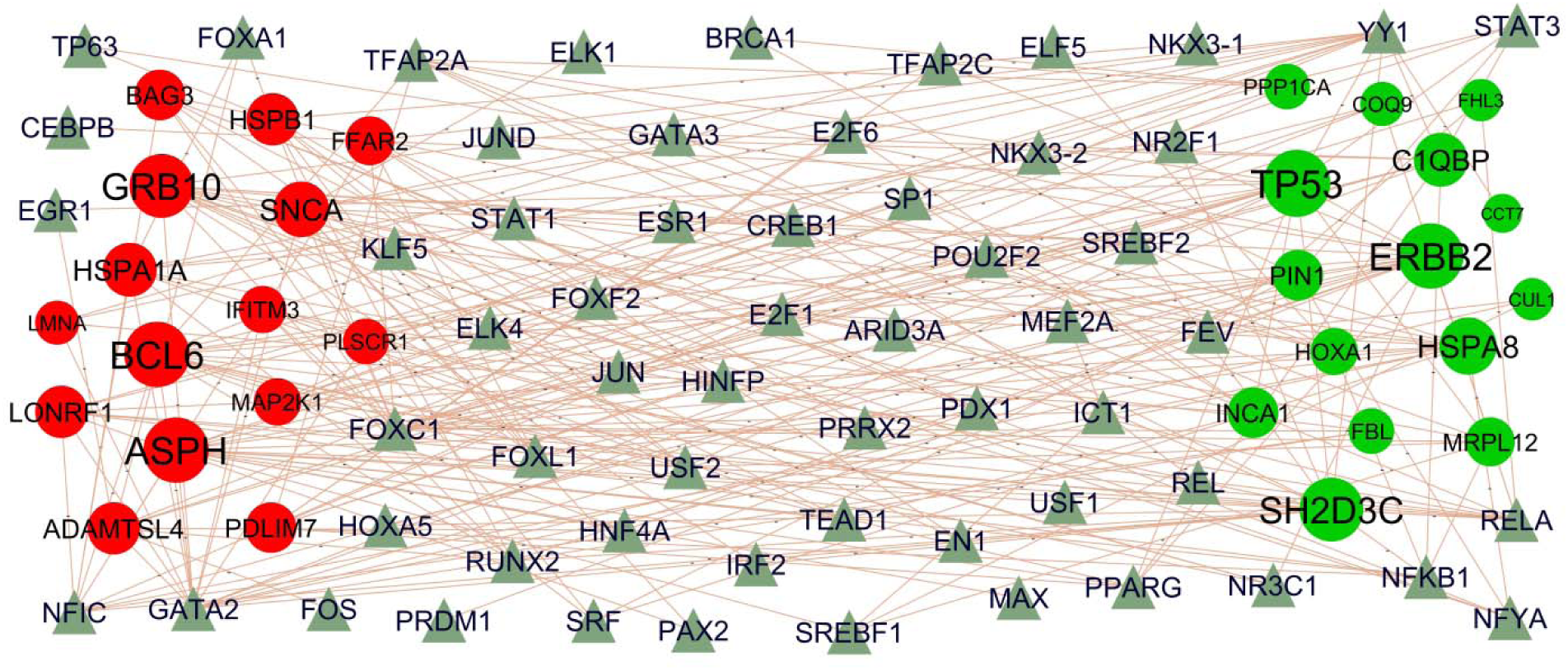
Hub gene - TF regulatory network. The olive green color triangle nodes represent the key TFs; up regulated genes are marked in dark green; down regulated genes are marked in dark red.

### Receiver operating characteristic curve (ROC) analysis

The ten hub genes were used as independent indicators to test the diagnostic efficacy and displayed by the ROC curve (Fig.7). The results revealed that CUL1 (AUC□=□0.905), HSPA8 (AUC□=□0.929), HOXA1 (AUC□=□0.909), INCA1 (AUC□=□0.917), TP53 (AUC□=□0.904), HSPB1 (AUC□=□0.939), LMNA (AUC□=□0.903), SNCA (AUC□=□0.915), ADAMTSL4 (AUC□=□0.901) and PDLIM7 (AUC□=□0.919) have good diagnostic efficacy as independent indicators for CA.

**Fig. 7.**
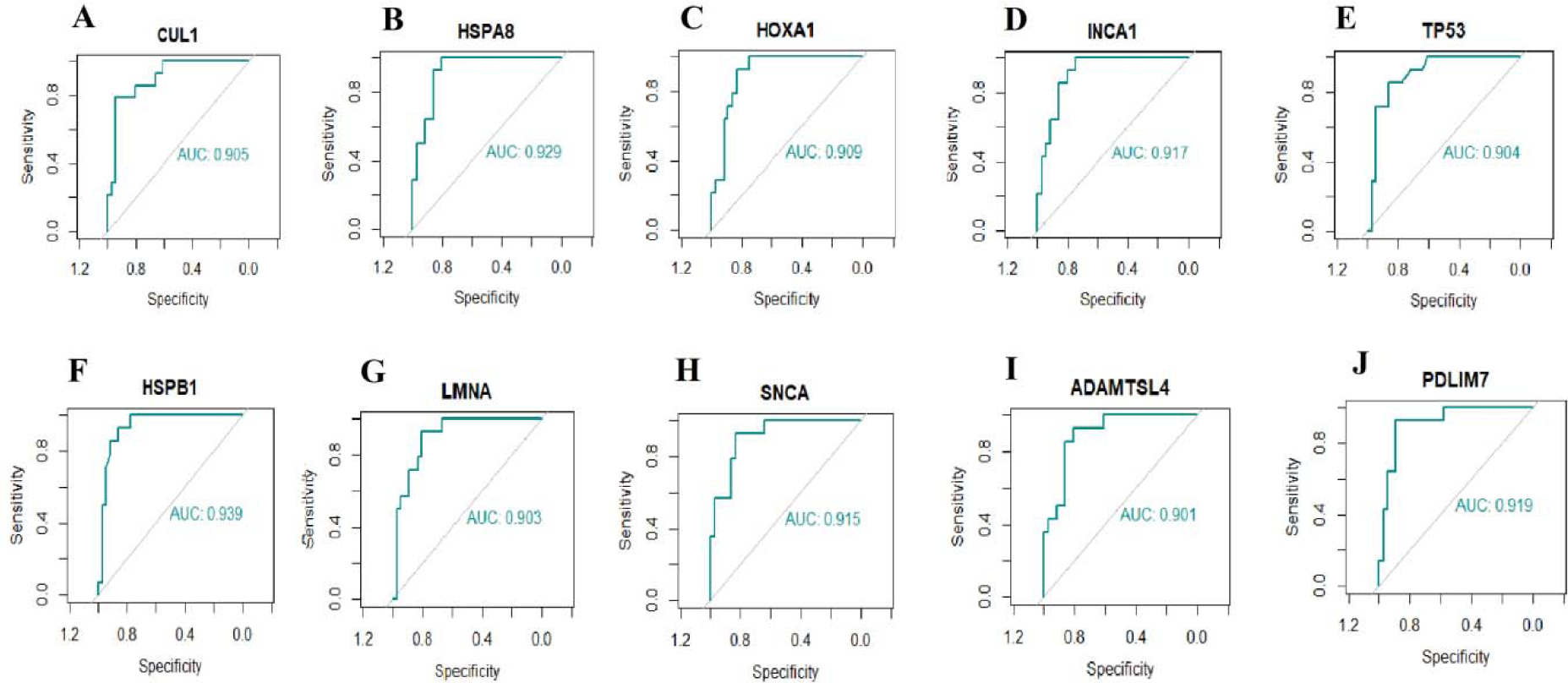
ROC curve analyses of hub genes. A) CUL1 B) HSPA8 C) HOXA1 D) INCA1 E) TP53 F) HSPB1 G) LMNA H) SNCA I) ADAMTSL4 J) PDLIM7

## Discussion

Despite advances in adjunctive pharmacotherapy and various surgical methods of CA, it is still the leading threat to human health in cardio vascular diseases [49–50]. Thus, successful screening techniques and accurate diagnosis remain the great challenges for decreasing the incidence of CA. In the current investigation, integrated bioinformatics analysis was used to identify the potential key genes related to CA. By performing DEGs analysis, 414 up regulated and 430 down regulated genes were successfully identified (P<0.05, [log□FC] > 0 for up regulated genes, log□FC] < −0.71 for down regulated genes and adjust P-value□<□.05), respectively. PRSS57 [51], IL1R2 [52], HBA2 [53], CLU (clusterin) [54], RNASE1 [55], THBS1 [56] and RGS1 [57] are associated with the prognosis of neurologic complications. TIFAB (TIFA inhibitor) [58], IL1R2 [59], CLU (clusterin) [60], RNASE1 [61], AREG (amphiregulin) [62], METTL7B [63], THBS1 [64], RGS1 [65] and VSIG4 [66] plays an important role in the inflammation. HBB (hemoglobin subunit beta) [67], CLU (clusterin) [68], AREG (amphiregulin) [69], THBS1 [70] and VSIG4 [71] were identified as a candidate biomarker for the diagnosis and treatment of diabetes mellitus. IL1R2 [72] and CLU (clusterin) [68] were reported to be associated with the prognosis of coronary artery disease. IL1R2 [73], CLU (clusterin) [74], RNASE1 [75] and VSIG4 [76] are a potential prognostic markers in myocardial ischemia. HBA2 [77], CLU (clusterin) [78], AREG (amphiregulin) [79], THBS1 [80], RGS1 [81] and VSIG4 [71] have been identified as a key genes in hypertension. CLU (clusterin) [82] and VSIG4 [83] might be associated with poor prognosis of heart failure. CLU (clusterin) [84] and RGS1 [85] plays significant roles in atherosclerosis. CLU (clusterin) [86], AREG (amphiregulin) [87] and VSIG4 [88] are mainly related to myocardial infarction. CLU (clusterin) [89] and THBS1 [90] was observed in obesity. RNASE1 [55] was found to regulate the progression of CA. Altered expression of RNASE1 [91] is associated with cardiomyopathy. Altered expression of AREG (amphiregulin) [92] and THBS1 [93] are involved in the development and progression of cardiac hypertrophy. Altered expression of THBS1 [94] and VSIG4 [88] were found to promote the cardiac fibrosis. The results clearly indicate that the genes discovered by DEG analysis might be essential for conditions beyond than underlying CA.

For a more in-depth understanding of these DEGs, we analyzed the selected genes for GO and REACTOME pathway enrichment analyses. Signaling pathways include translation [95], immune system [96] and signal transduction [97] were responsible for progression of CA. Recently, increasing evidence demonstrated that CKB (creatine kinase B) [98], CX3CR1 [99], S1PR2 [100], ADAM8 [101], HINT2 [102], CDK4 [103], TP53 [104], RGS14 [105], FUS (FUS RNA binding protein) [106], TRIM21 [107], IDH2 [108], ECHS1 [109], CXCR1 [110], TBC1D10C [111], POMC (proopiomelanocortin) [112], SCAP (SREBF chaperone) [113], LIMD2 [114], PFN1 [115], DKC1 [116], NUP93 [117], ADM (adrenomedullin) [118], ELANE (elastase, neutrophil expressed) [119], CD163 [120], G0S2 [121], IL10 [122], SOCS3 [123], S100P [124], TIMP4 [125], HP (haptoglobin) [126], SDC4 [127], FKBP5 [128], AQP9 [129], ACSL1 [130], CXCR4 [131], LMNA (lamin A/C) [132], S100A8 [133], ANXA3 [134], NAMPT (nicotinamidephosphoribosyltransferase) [135], ALAS2 [136], S100A12 [137], LRG1 [138], CTSL (cathepsin L) [139], TIMP1 [140], F5 [141], GAS6 [142], F2RL3 [143], S100A9 [144], CR1 [145], GFPT2 [146], PF4 [147], FPR2 [148], P2RY1 [149], TFPI (tissue factor pathway inhibitor) [150], HLX (H2.0 like homeobox) [151], DDIT4 [152], OSM (oncostatin M) [153], NR4A2 [154], KCNE1 [155], PIK3R1 [156], TREM1 [157], CTSB (cathepsin B) [158], FFAR2 [159], CXCL2 [160], ECM1 [161], CYP1B1 [162], IL1R1 [163], HSPB1 [164], CDKN1A [165], LPL (lipoprotein lipase) [166], CLEC5A [167], NEAT1 [168], HBEGF (heparin binding EGF like growth factor) [169], BAG3 [170], TRIB1 [171], PDE4D [172], TIMP2 [173], DACH1 [174], ABCA1 [175], KLF9 [176], BRCA2 [177], CCR1 [178], EPHB2 [179], PDE4B [180], FLT3 [181], HMGB3 [182], CD63 [183], SOD2 [184], CDKN2B [185], ADORA2B [186], IFNGR1 [187], PLXND1 [188], HIF1A [189], GPX1 [190], BCL2L11 [191], VCAN (versican) [192], PFKFB3 [193], BCAT1 [194], MFGE8 [195], MBOAT7 [196] and PPP1R3B [197] were altered expressed in myocardial infarction. Studies had shown that CKB (creatine kinase B) [198], CX3CR1 [199], MRPL12 [200], ADA (adenosine deaminase) [201], NARS2 [202], MIPEP (mitochondrial intermediate peptidase) [203], S1PR2 [204], TRIB3 [205], PIN1 [206], ADAM8 [207], TSPAN32 [208], MMAB (metabolism of cobalamin associated B) [209], CAMK1D [210], HSPA8 [211], PIGC (phosphatidylinositol glycan anchor biosynthesis class C) [212], ARV1 [213], SETD3 [214], PEBP1 [215], CAPRIN1 [216], TP53 [217], FUS (FUS RNA binding protein) [218], CD300A [219], DOLK (dolichol kinase) [220], TRAP1 [221], IDH2 [222], MRPL4 [223], EEF1A1 [224], ECHS1 [225], PHB2 [226], FIS1 [227], NEFH (neurofilament heavy chain) [228], SIGMAR1 [229], ANXA6 [230], TRAPPC6A [231], UBTF (upstream binding transcription factor) [232], TMEM175 [233], PFN1 [234], ATXN10 [235], TRIT1 [236], LAS1L [237], CCDC86 [238], THG1L [239], TMEM230 [240], SLC25A11 [241], LSP1 [242], ACO2 [243], ADM (adrenomedullin) [244], CD163 [245], CXCL8 [246], IL10 [247], PROK2 [248], SOCS3 [249], S100P [250], SOCS1 [251], EDNRB (endothelin receptor type B) [252], GPER1 [253], HP (haptoglobin) [254], FKBP5 [255], AQP9 [256], KREMEN1 [257], CXCR4 [258], RETN (resistin) [259], LMNA (lamin A/C) [260], S100A8 [261], ANXA3 [262], NAMPT (nicotinamidephosphoribosyltransferase) [263], S100A12 [264], LRG1 [265], FOSL1 [266], TIMP1 [267], GPR84 [268], ALDH1A2 [269], PER1 [270], GAS6 [271], S100A9 [272], CES1 [273], CR1 [274], PF4 [275], FPR2 [276], MSR1 [277], P2RY1 [278], TFPI (tissue factor pathway inhibitor) [279], DYSF (dysferlin) [280], IRS2 [281], SMOX (spermine oxidase) [282], NR4A2 [283], TREM1 [284], CTSB (cathepsin B) [285], HMOX1 [286], MAFB (MAF bZIP transcription factor B) [287], GLUL (glutamate-ammonia ligase) [288], SAMSN1 [289], LPL (lipoprotein lipase) [290], FZD2 [291], NEAT1 [292], HTRA1 [293], HBEGF (heparin binding EGF like growth factor) [294], C5AR1 [295], DGAT2 [296], CEBPB (CCAAT enhancer binding protein beta) [297], GAS2L1 [298], BAG3 [299], CCL3 [300], TRIB1 [301], BCL6 [302], SNCA (synuclein alpha) [303], PDE4D [304], TIMP2 [305], IMPA2 [306], ABCA1 [307], IRAK3 [308], EPHB2 [309], PDE4B [310], CEBPD (CCAAT enhancer binding protein delta) [311], GNAQ (G protein subunit alpha q) [312], CD63 [313], SOD2 [314], ADCY3 [315], ERN1 [316], CTSD (cathepsin D) [317], PLSCR1 [318], MAP3K6 [319], MCEMP1 [320], TUBB2A [321], SYTL3 [322], TCN2 [323], LIMK2 [324], SLC2A3 [325], BASP1 [326], MBOAT7 [327], PPP1R3B [328], TTC7B [329], SLC22A4 [330], FTL (ferritin light chain) [331], TUBB6 [242], CTSA (cathepsin A) [332] and CMIP (c-Maf inducing protein) [333] were associated with neurologic complications. CX3CR1 [334], ADA (adenosine deaminase) [335], ERBB2 [336], DYRK1B [337], TP53 [104], RGS14 [105], FUS (FUS RNA binding protein) [338], RBCK1 [339], OGG1 [340], PHB2 [341], MCM2 [342], TBC1D10C [111], POMC (proopiomelanocortin) [343], PALLD (palladin, cytoskeletal associated protein) [344], SIGMAR1 [345], SNAP29 [346], NUP93 [117], ADM (adrenomedullin) [347], CD163 [348], CXCL8 [349], IL10 [350], ITGA7 [351], SOCS3 [352], SOCS1 [353], GPER1 [354], TIMP4 [355], HP (haptoglobin) [356], SDC4 [127], CXCR4 [357], RETN (resistin) [358], CREM (cAMP responsive element modulator) [359], S100A8 [133], NAMPT (nicotinamidephosphoribosyltransferase) [360], S100A12 [361], LRG1 [362], TIMP1 [363], EPAS1 [364], GAS6 [365], S100A9 [366], CES1 [367], FPR2 [368], IRS2 [369], OSM (oncostatin M) [370], DSC2 [371], KCNE1 [372], TREM1 [373], NFIL3 [374], AHR (aryl hydrocarbon receptor) [375], NEAT1 [376], HRH2 [377], BAG3 [378], PDK4 [379], PDE4D [380], TIMP2 [381], EGFL7 [382], BRCA2 [383], PDE4B [384], MPP1 [385], GNAQ (G protein subunit alpha q) [386], SOD2 [387], CDKN2B [388], ADORA2B [389], SMAD1 [390], CTSD (cathepsin D) [391], PRKCE (protein kinase C epsilon) [392], GPX1 [393], PFKFB3 [394], ETS2 [395], SLC2A3 [396], MYBPC3 [397] and TP53INP2 [398] might be a potential therapeutic targets for heart failure treatment. Abnormal regulation of CX3CR1 [399], ADA (adenosine deaminase) [400], ERBB2 [401], S1PR2 [402], TRIB3 [403], PIN1 [404], MMAB (metabolism of cobalamin associated B) [209], HSPA8 [405], TP53 [406], IDH2 [407], OGG1 [408], CXCR1 [409], PROCR (protein C receptor) [410], INSIG1 [411], NUP93 [117], ADM (adrenomedullin) [412], CD163 [413], G0S2 [121], CXCL8 [414], IL10 [415], S100P [124], LYVE1 [416], SOCS1 [417], TIMP4 [418], HP (haptoglobin) [419], FKBP5 [420], CXCR4 [421], RETN (resistin) [422], LMNA (lamin A/C) [423], S100A8 [424], ANXA3 [425], NAMPT (nicotinamidephosphoribosyltransferase) [426], ALOX15B [427], PPBP (pro-platelet basic protein) [428], S100A12 [429], LRG1 [430], TIMP1 [431], F5 [432], GAS6 [433], F2RL3 [434], S100A9 [435], TREML4 [436], CES1 [437], CR1 [145], BCL3 [438], PF4 [439], FPR2 [440], MSR1 [441], P2RY1 [442], GRB10 [443], TFPI (tissue factor pathway inhibitor) [444], SLC39A8 [445], DYSF (dysferlin) [446], IRS2 [447], TNFAIP3 [448], OSM (oncostatin M) [449], NR4A2 [450], KCNE1 [451], TREM1 [452], CXCL2 [453], HMOX1 [454], MAFB (MAF bZIP transcription factor B) [287], MERTK (MER proto-oncogene, tyrosine kinase) [455], CYP1B1 [456], CSF1 [457], LPL (lipoprotein lipase) [458], CD55 [459], AHR (aryl hydrocarbon receptor) [460], NEAT1 [461], HTRA1 [462], CDH5 [463], HBEGF (heparin binding EGF like growth factor) [464], C5AR1 [465], PDK4 [466], CCL3 [467], TRIB1 [301], PDE4D [468], TIMP2 [173], DACH1 [174], ABCA1 [469], CD63 [470], SOD2 [471], CDKN2B [472], ADCY3 [473], BTG1 [474], GPX1 [475], MT2A [476], VCAN (versican) [477], SYTL3 [322], MFGE8 [478], MBOAT7 [479], PPP1R3B [328], MYBPC3 [480] and SIGLEC12 [481] are associated with coronary artery disease. Studies have found that CX3CR1 [482], ADA (adenosine deaminase) [483], ST6GAL1 [484], TRIB3 [485], PIN1 [486], ADAM8 [487], CUL1 [488], PEBP1 [489], GLRX3 [490], TP53 [491], FUS (FUS RNA binding protein) [492], TRIM21 [493], OGG1 [494], HOXA1 [495], MCM2 [342], POMC (proopiomelanocortin) [496], PALLD (palladin, cytoskeletal associated protein) [497], SCAP (SREBF chaperone) [498], PFN1 [499], RAC2 [500], CNN2 [501], ADM (adrenomedullin) [502], CD163 [503], G0S2 [121], ID1 [504], CXCL8 [505], IL10 [506], SOCS3 [507], SOCS1 [508], GPER1 [509], HP (haptoglobin) [510], SDC4 [511], ITGB3 [512], AQP9 [513], RGL1 [514], ACSL1 [515], CXCR4 [516], RETN (resistin) [517], LMNA (lamin A/C) [518], S100A8 [519], NAMPT (nicotinamidephosphoribosyltransferase) [520], ALOX15B [521], S100A12 [522], LRG1 [523], CTSL (cathepsin L) [524], TIMP1 [431], GAS6 [525], EREG (epiregulin) [526], S100A9 [527], TREML4 [436], CES1 [528], BCL3 [529], PF4 [530], STAB1 [531], FPR2 [532], MSR1 [533], P2RY1 [534], TFPI (tissue factor pathway inhibitor) [535], DYSF (dysferlin) [536], IRS2 [537], OSM (oncostatin M) [538], ASPH (aspartate beta-hydroxylase) [539], FPR1 [540], NR4A2 [450], PIK3R1 [541], TREM1 [542], NFIL3 [543], CTSB (cathepsin B) [544], CXCL2 [545], HMOX1 [546], MAFB (MAF bZIP transcription factor B) [547], MERTK (MER proto-oncogene, tyrosine kinase) [548], CYP1B1 [549], ADAM9 [550], CDKN1A [551], LPL (lipoprotein lipase) [552], DOCK4 [553], CD55 [554], AHR (aryl hydrocarbon receptor) [555], NEAT1 [556], FZD5 [557], HBEGF (heparin binding EGF like growth factor) [558], DGAT2 [559], HRH2 [377], VNN1 [560], BAG3 [561], LILRB4 [562], FABP5 [563], TRIB1 [564], PDE3B [565], BCL6 [566], IL1RAP [567], PDE4D [568], TIMP2 [569], ZFP36 [570], IRAK3 [571], ENPP2 [572], CCR1 [573], EPHB2 [574], FLT3 [575], CD63 [313], SOD2 [576], CDKN2B [577], CCRL2 [578], SMAD1 [579], JAK3 [580], PLXND1 [581], HIF1A [582], GPX1 [583], MT2A [584], SPHK1 [585], C2 [586], SDC2 [587], VCAN (versican) [477], PFKFB3 [588], BCAT1 [589], ETS2 [590], MFGE8 [591], PPP1R3B [592], PDLIM7 [593], SIGLEC12 [481], and LILRB4 [562] are altered expressed in atherosclerosis. CX3CR1 [594], ERBB2 [595], MIPEP (mitochondrial intermediate peptidase) [596], TRIB3 [597], ARV1 [598], C1QBP [599], TP53 [600], RGS14 [105], DOLK (dolichol kinase) [601], TRAP1 [602], RBCK1 [603], IDH2 [604], OGG1 [605], NDUFB5 [606], ECHS1 [607], PHB2 [608], FIS1 [609], PALLD (palladin, cytoskeletal associated protein) [610], SIGMAR1 [611], SNAP29 [346], NUP93 [117], LSP1 [612], ELMO1 [613], ADM (adrenomedullin) [614], IL10 [615], LYVE1 [616], HP (haptoglobin) [617], FKBP5 [618], CXCR4 [619], RETN (resistin) [620], LMNA (lamin A/C) [621], CREM (cAMP responsive element modulator) [622], S100A8 [623], NAMPT (nicotinamidephosphoribosyltransferase) [624], S100A12 [625], CTSL (cathepsin L) [626], TIMP1 [627], TREML4 [628], CES1 [629], P2RY1 [630], SLC39A8 [631], DYSF (dysferlin) [632], IRS2 [633], NR4A2 [634], PIK3R1 [635], TREM1 [373], CTSB (cathepsin B) [636], HMOX1 [637], LPL (lipoprotein lipase) [638], AHR (aryl hydrocarbon receptor) [639], NEAT1 [640], HTRA1 [641], BAG3 [642], PDK4 [643], TIMP2 [644], KLF9 [645], BRCA2 [646], CCR1 [647], FBN2 [648], SOD2 [649], HIF1A [650], GPX1 [651], PFKFB3 [652], MYO7A [653], MFGE8 [654], MYBPC3 [397] and LMNB1 [655] plays an indispensable role in cardiomyopathy. CX3CR1 [656], ADA (adenosine deaminase) [657], RNF126 [658], ERBB2 [659], PARP10 [660], MIPEP (mitochondrial intermediate peptidase) [596], PIN1 [661], DYRK1B [337], CDK4 [662], HSPA8 [663], PEBP1 [664], CTRL (chymotrypsin like) [665], RNF5 [666], TP53 [667], RGS14 [105], FUS (FUS RNA binding protein) [668], IDH2 [669], OGG1 [670], TBC1D10C [111], RCN1 [671], POMC (proopiomelanocortin) [672], ANXA6 [673], PFN1 [674], LSP1 [612], ADM (adrenomedullin) [675], IL10 [676], LYVE1 [616], GPER1 [677], TIMP4 [678], HP (haptoglobin) [617], FKBP5 [679], ACSL1 [680], RETN (resistin) [681], S100A8 [682], NAMPT (nicotinamidephosphoribosyltransferase) [683], CTSL (cathepsin L) [684], ME1 [685], CDC20 [686], GAS6 [687], EREG (epiregulin) [688], S100A9 [689], ITGA7 [351], CES1 [629], P2RY1 [630], SLC39A8 [631], FAM20C [690], DSC2 [371], PIK3R1 [691], NFIL3 [692], CTSB (cathepsin B) [158], HMOX1 [693], CYP1B1 [694], ADAM9 [695], LPL (lipoprotein lipase) [696], AHR (aryl hydrocarbon receptor) [697], NEAT1 [698], HBEGF (heparin binding EGF like growth factor) [699], BAG3 [700], LILRB4 [701], PDE4D [702], TIMP2 [703], BRCA2 [704], FLT3 [705], GNAQ (G protein subunit alpha q) [706], SOD2 [707], HIF1A [650], GPX1 [708], SWAP70 [709], ETS2 [710], MFGE8 [711], MYBPC3 [712], TP53INP2 [713], STEAP3 [714] and LILRB4 [715] were frequently altered in cardiac hypertrophy. CX3CR1 [716], RNF126 [658], TRIB3 [717], PIN1 [718], GGT7 [719], CAMK1D [720], HSPA8 [721], SETD3 [722], PPP1CA [723], PEBP1 [724], TP53 [725], TRIM21 [726], IDH2 [727], OGG1 [728], MRPL4 [223], ALAD (aminolevulinatedehydratase) [729], FIS1 [730], CXCR1 [731], ARMC5 [732], CRIP1 [733], SIGMAR1 [734], PFN1 [735], CNN2 [736], ADM (adrenomedullin) [737], CD163 [738], CXCL8 [739], IL10 [740], TIMP4 [741], HP (haptoglobin) [742], SDC4 [743], ITGB3 [744], CXCR4 [745], RETN (resistin) [746], LMNA (lamin A/C) [747], NTSR1 [748], NAMPT (nicotinamidephosphoribosyltransferase) [749], S100A12 [750], CTSL (cathepsin L) [751], TIMP1 [752], ME1 [753], PER1 [754], GAS6 [755], F2RL3 [756], S100A9 [757], CES1 [758], PF4 [759], TFPI (tissue factor pathway inhibitor) [760], TNFAIP3 [761], FPR1 [762], ASGR2 [763], PIK3R1 [764], ACOX2 [765], CTSB (cathepsin B) [766], HMOX1 [693], CYP1B1 [767], IL1R1 [768], LPL (lipoprotein lipase) [769], TSPYL2 [770], AHR (aryl hydrocarbon receptor) [771], HPGD (15-hydroxyprostaglandin dehydrogenase) [772], NEAT1 [773], HBEGF (heparin binding EGF like growth factor) [774], C5AR1 [775], VNN1 [776], BAG3 [777], FABP5 [778], PDK4 [779], BCL6 [780], ITGA9 [781], PDE4D [782], TIMP2 [783], ABCA1 [784], BRCA2 [704], PDE4B [785], CEBPD (CCAAT enhancer binding protein delta) [786], SOD2 [787], CDKN2B [788], ADORA2B [789], SMAD1 [790], HIF1A [791], PRKCE (protein kinase C epsilon) [392], GPX1 [708], SPHK1 [792], VCAN (versican) [793], TCN2 [794], PFKFB3 [795], MFGE8 [796], KLHL2 [797] and ATP13A3 [798] could be an early detection biomarker for hypertension. CX3CR1 [799], ADA (adenosine deaminase) [800], ST6GAL1 [801], ERBB2 [802], FBL (fibrillarin) [803], S1PR2 [804], NEIL2 [805], PIN1 [806], ADAM8 [807], NUAK2 [808], ICMT (isoprenylcysteine carboxyl methyltransferase) [809], PREP (prolylendopeptidase) [810], DYRK1B [811], PIAS4 [812], CDK4 [813], STK38 [814], TAOK3 [815], BCL2L12 [816], PEBP1 [215], CTRL (chymotrypsin like) [817], C1QBP [818], RNF5 [819], TP53 [667], NDUFA12 [820], CD300A [821], PPM1G [822], TRAP1 [823], RNF20 [824], TRIM21 [107], RBCK1 [825], ARHGEF2 [826], IDH2 [827], OGG1 [828], ECHS1 [829], FIS1 [830], MCM2 [831], CXCR1 [832], POMC (proopiomelanocortin) [833], APBA3 [834], DDX54 [835], CLUH (clustered mitochondria homolog) [836], FCHO1 [837], SIGMAR1 [838], RNASEH2A [839], RNASEH2B [840], SLC18B1 [841], LIMD2 [114], RAB11FIP1 [842], TMEM175 [843], APEX1 [844], PFN1 [845], ATXN10 [846], SUMF2 [847], RAC2 [848], ELF4 [849], CNN2 [850], FRMD8 [851], INTS9 [852], TNFAIP8L2 [853], LSP1 [854], ADM (adrenomedullin) [855], ELANE (elastase, neutrophil expressed) [856], CD163 [857], G0S2 [858], ID1 [859], TNFAIP6 [860], CXCL8 [861], IL10 [862], PROK2 [248], SOCS3 [863], LYVE1 [864], SOCS1 [508], EDNRB (endothelin receptor type B) [865], TIMP4 [866], HP (haptoglobin) [742], FAM20A [867], SDC4 [868], FKBP5 [128], AQP9 [129], KREMEN1 [257], ACSL1 [869], GPER1 [870], CXCR4 [871], RETN (resistin) [872], LMNA (lamin A/C) [873], CREM (cAMP responsive element modulator) [874], NTSR1 [875], S100A8 [876], ANXA3 [877], RNASE2 [878], NAMPT (nicotinamidephosphoribosyltransferase) [879], ALOX15B [521], S100A12 [522], LRG1 [880], CTSL (cathepsin L) [881], FOSL1 [266], TMEM45B [882], TIMP1 [883], EPAS1 [884], GPR84 [885], SLC7A5 [886], LAMC1 [887], PER1 [888], GAS6 [755], EREG (epiregulin) [889], S100A9 [527], S100A9 [757], TREML4 [890], ITGA7 [891], CES1 [892], BCL3 [893], PF4 [894], FPR2 [276], MSR1 [895], TFPI (tissue factor pathway inhibitor) [896], DYSF (dysferlin) [897], IRS2 [898], TNFAIP3 [899], EBI3 [900], DDIT4 [901], OSM (oncostatin M) [902], SMOX (spermine oxidase) [903], ASPH (aspartate beta-hydroxylase) [539], FPR1 [904], NR4A2 [905], DSC2 [906], PIK3R1 [907], TREM1 [908], NFIL3 [909], CTSB (cathepsin B) [910], FFAR2 [911], CXCL2 [912], DUSP2 [913], HMOX1 [914], MAFB (MAF bZIP transcription factor B) [915], ECM1 [161], CYP1B1 [916], ADAM9 [917], SLC11A1 [918], IL1R1 [919], HSPB1 [920], CDKN1A [921], CSF1 [922], LPL (lipoprotein lipase) [923], CLEC5A [924], CD55 [925], AHR (aryl hydrocarbon receptor) [926], NEAT1 [927], HTRA1 [928], FZD5 [557], HBEGF (heparin binding EGF like growth factor) [929], TGM2 [930], C5AR1 [295], DGAT2 [931], HRH2 [932], PROS1 [933], CEBPB (CCAAT enhancer binding protein beta) [934], VNN1 [935], LILRB4 [562], E2F2 [936], FABP5 [937], PDK4 [938], CCL3 [939], PDE3B [940], NFE2 [941], BCL6 [942], CEACAM3 [943], MMP19 [944], SNCA (synuclein alpha) [945], C3AR1 [946], IL1RAP [567], PDE4D [947], TIMP2 [948], DACH1 [949], BCL2A1 [950], ABCA1 [469], EGFL7 [951], ZFP36 [570], KLF9 [176], SRGN (serglycin) [952], IRAK3 [953], BRCA2 [954], ENPP2 [572], CCR1 [178], GPR183 [955], EPHB2 [309], MAP3K8 [956], PDE4B [957], FLT3 [958], SBNO2 [959], CEBPD (CCAAT enhancer binding protein delta) [960], MAP2K1 [961], FTH1 [962], CD63 [963], IFITM3 [964], SOD2 [965], CDKN2B [966], CCRL2 [967], ADORA2B [968], SMAD1 [969], JAK3 [970], ERN1 [971], CORO2A [972], PDE4A [973], CTSD (cathepsin D) [974], BTG1 [975], HIF1A [976], CPEB4 [977], COCH (cochlin) [978], ODC1 [979], PRKCE (protein kinase C epsilon) [980], GPX1 [981], MT2A [584], PLSCR1 [982], IL4R [983], SPHK1 [984], BCL2L11 [985], SWAP70 [986], P2RX4 [987], SDC2 [988], VCAN (versican) [989], PFKFB3 [990], ADAM19 [991], BCAT1 [992], SERPINB10 [993], GCA (grancalcin) [994], ETS2 [995], MFGE8 [996], BASP1 [997], MBOAT7 [998], CYB5R2 [999], P4HA2 [1000], MYBPC3 [1001], SCRG1 [1002], SLC22A4 [1003], DRAM1 [1004], FTL (ferritin light chain) [1005], PDLIM7 [1006], TP53INP2 [1007], STEAP3 [1008] and LILRB4 [562] expression had been confirmed in inflammation. A study indicated that the altered expression of CX3CR1 [1009], ADA (adenosine deaminase) [1010], TRIB3 [1011], PIN1 [1012], DYRK1B [1013], CAMK1D [1014], CDK4 [1015], TAOK3 [1016], PEBP1 [1017], GLRX3 [1018], TP53 [725], TRIM21 [1019], IDH2 [1020], OGG1 [1021], FASTK (Fas activated serine/threonine kinase) [1022], RPS6KA1 [1023], ECHS1 [1024], CLIC5 [1025], POMC (proopiomelanocortin) [1026], INSIG1 [1027], SCAP (SREBF chaperone) [1028], NOB1 [1029], PFN1 [1030], DKC1 [116], CTNNBL1 [1031], ADM (adrenomedullin) [1032], ELANE (elastase, neutrophil expressed) [856], CD163 [1033], ID1 [1034], CXCL8 [1035], IL10 [1036], SOCS3 [1037], LYVE1 [1038], SOCS1[1037], TIMP4 [1039], HP (haptoglobin) [742], FKBP5 [1040], AQP9 [1041], ACSL1 [1042], CXCR4 [1043], RETN (resistin) [1044], LMNA (lamin A/C) [1045], S100A8 [1046], NAMPT (nicotinamidephosphoribosyltransferase) [1047], LRG1 [1048], CTSL (cathepsin L) [881], TIMP1 [1049], ME1 [1050], GPR84 [1051], SLC7A5 [1052], LAMC1 [1053], F5 [1054], GAS6 [1055], EREG (epiregulin) [1056], S100A9 [1046], CES1 [1057], MSR1 [1058], TFPI (tissue factor pathway inhibitor) [1059], IRS2 [1060], EBI3 [1061], DDIT4 [1062], OSM (oncostatin M) [1063], SLC7A8 [1064], PIK3R1 [1065], TREM1 [1066], CTSB (cathepsin B) [1067], CXCL2 [1068], DUSP2 [1069], HMOX1 [1070], MAFB (MAF bZIP transcription factor B) [1071], ECM1 [1072], HSPB1 [1073], APCDD1 [1074], LPL (lipoprotein lipase) [1075], AHR (aryl hydrocarbon receptor) [1076], NEAT1 [1077], HBEGF (heparin binding EGF like growth factor) [464], DGAT2 [1078], CEBPB (CCAAT enhancer binding protein beta) [1079], FABP5 [1080], PDK4 [1081], PDE3B [1082], BCL6 [1083], MMP19 [1084], C3AR1 [1085], IL1RAP [1086], PDE4D [1087], TIMP2 [1088], DACH1 [1089], ABCA1 [1090], ZFP36 [1091], KLF9 [1092], SRGN (serglycin) [952], IRAK3 [571], BRCA2 [1093], ENPP2 [1094], CCR1 [1095], EPHB2 [1096], PDE4B [1097], FLT3 [1098], CEBPB (CCAAT enhancer binding protein delta) [1079], IFITM3 [1099], SOD2 [1100], CCRL2 [1101], JAK3 [1102], ADCY3 [1103], CTSD (cathepsin D) [1104], HIF1A [1105], CPEB4 [1106], GPX1 [1107], SPHK1 [1108], VCAN (versican) [989], TCN2 [1109], PFKFB3 [1110], GCA (grancalcin) [994], MFGE8 [1111], MBOAT7 [1112], PPP1R3B [1113] and CMIP (c-Maf inducing protein) [1114] are a critical determinant of obesity. CX3CR1 [1115], ADA (adenosine deaminase) [1010], ERBB2 [1116], NARS2 [1117], S1PR2 [1118], TRIB3 [403], PIN1 [1119], HINT2 [1120], DYRK1B [1013], CAMK1D [720], OXSM (3-oxoacyl-ACP synthase, mitochondrial) [1121], CDK4 [1015], HSPA8 [1122], PEBP1 [1017], TP53 [725], TRIM21 [1123], IDH2 [1124], OGG1 [1125], EIF2B1 [1126], ECHS1 [1024], HLA-DQA2 [1127], CXCR1 [1128], PLEK (pleckstrin) [1129], SCAP (SREBF chaperone) [1028], NOB1 [1130], SNAP29 [346], ENO3 [1131], TNFAIP8L2 [1132], ELMO1 [1133], BST2 [1134], ADM (adrenomedullin) [1135], CD163 [1033], CXCL8 [1136], IL10 [1036], PROK2 [1137], SOCS3 [1138], LYVE1 [1038], SOCS1 [1139], TIMP4 [418], HP (haptoglobin) [742], FKBP5 [1040], ITGB3 [512], AQP9 [1140], ACSL1 [1141], CXCR4 [1142], RETN (resistin) [1044], LMNA (lamin A/C) [747], SMAD1 [1143], CREM (cAMP responsive element modulator) [1144], S100A8 [1145], ANXA3 [1146], NAMPT (nicotinamidephosphoribosyltransferase) [1147], S100A12 [361], CTSL (cathepsin L) [1148], FOSL1 [1149], C1QB [1150], TIMP1 [1151], GPR84 [1152], GAS6 [525], S100A9 [1153], BCL3 [1154], GFPT2 [1155], PF4 [1156], FPR2 [1157], MSR1 [1058], P2RY1 [1158], GRB10 [443], TFPI (tissue factor pathway inhibitor) [1059], IRS2 [369], TNFAIP3 [448], OSM (oncostatin M) [1159], PIK3R1 [156], TREM1 [1160], CTSB (cathepsin B) [1161], DUSP2 [1069], HMOX1 [1162], GLUL (glutamate-ammonia ligase) [288], SLC11A1 [1163], IL1R1 [1164], HSPB1 [1165], CDKN1A [1166], CSF1 [1167], LPL (lipoprotein lipase) [1168], CD55 [1169], AHR (aryl hydrocarbon receptor) [1170], HPGD (15-hydroxyprostaglandin dehydrogenase) [1171], NEAT1 [1172], HBEGF (heparin binding EGF like growth factor) [1173], TGM2 [1174], DGAT2 [1175], VNN1 [1176], BAG3 [777], E2F2 [1177], FABP5 [1178], PDK4 [1179], CCL3 [467], PDE3B [1180], PLK3 [1181], TIMP2 [1182], DACH1 [1089], ABCA1 [1090], ZFP36 [1091], IRAK3 [571], ENPP2 [1094], PDE4B [1183], ECHDC3 [1184], IFITM3 [1185], SOD2 [1186], CDKN2B [1187], CCRL2 [1101], JAK3 [1188], ADCY3 [1189], CTSD (cathepsin D) [1190], HIF1A [1191], GPX1 [1192], MT2A [584], IL4R [1193], SPHK1 [1194], C2 [1195], P2RX4 [1196], VCAN (versican) [1197], PFKFB3 [1198], BCAT1 [1199], GCA (grancalcin) [994], MFGE8 [1200], BASP1 [1201], PPP1R3B [1202], TP53INP2 [1203], STEAP3 [1204], ARRDC4 [1205] and CMIP (c-Maf inducing protein) [1114] were observed to be associated with the risk of diabetes mellitus. ADA (adenosine deaminase) [1206], ERBB2 [1207], S1PR2 [804], CTRL (chymotrypsin like) [1208], TP53 [1209], TRAP1 [1210], PKN1 [1211], IDH2 [669], OGG1 [1212], FASTK (Fas activated serine/threonine kinase) [1213], ECHS1 [1024], PHB2 [1214], CXCR1 [1215], FHL3 [1216], ADM (adrenomedullin) [1217], CD163 [1218], IL10 [1219], SOCS3 [1220], S100P [124], SOCS1 [1221], GPER1 [1222], HP (haptoglobin) [1223], SDC4 [127], ITGB3 [1224], CXCR4 [1225], RETN (resistin) [1226], S100A8 [1227], ALAS2 [1228], TIMP1 [363], GAS6 [1229], S100A9 [1227], PF4 [1230], PIM3 [1231], TFPI (tissue factor pathway inhibitor) [1232], DYSF (dysferlin) [1233], OSM (oncostatin M) [1159], FPR1 [1234], NR4A2 [1235], CTSB (cathepsin B) [1236], HMOX1 [1237], LPL (lipoprotein lipase) [1238], AHR (aryl hydrocarbon receptor) [1239], NEAT1 [1240], HTRA1 [641], BAG3 [1241], PDK4 [1242], TRIB1 [1243], PDE3B [1244], DACH1 [1245], ABCA1 [1246], KLF9 [1247], CCR1 [1248], SOD2 [1249], CDKN2B [388], ADORA2B [1250], SMAD1 [1251], JAK3 [1252], CTSD (cathepsin D) [1253], HIF1A [1254], PRKCE (protein kinase C epsilon) [1255], GPX1 [1256], SPHK1 [1257], BCL2L11 [1258], BCAT1 [1259], P4HA2 [1000] and ARRDC4 [1260] have been reported its expression in the myocardial ischemia. ADAM8 [101], HSPA8 [663], CTRL (chymotrypsin like) [1261], LYRM4 [1262], SOCS3 [1263], SOCS1 [1263], GPER1 [253], RETN (resistin) [1264], LMNA (lamin A/C) [1265], S100A8 [1227], S100A9 [1227], PF4 [1266], FPR2 [276], TFPI (tissue factor pathway inhibitor) [444], KCNE1 [451], CTSB (cathepsin B) [1267], AHR (aryl hydrocarbon receptor) [375], TIMP2 [1268], BRCA2 [1269], CTSD (cathepsin D) [1267], SPHK1 [1270] and MYBPC3 [712] have been detected in the CA patients. Altered expression of PIN1 [1271], ADAM8 [1272], HSPA8 [663], TP53 [600], OGG1 [1273], PHB2 [1274], CD163 [1275], IL10 [1276], SOCS3 [1277], CXCR4 [619], RETN (resistin) [681], NECTIN2 [1278], LMNA (lamin A/C) [1279], LRG1 [1280], TIMP1 [1281], EREG (epiregulin) [688], S100A9 [1282], ITGA7 [351], OSM (oncostatin M) [1283], DSC2 [906], AHR (aryl hydrocarbon receptor) [1284], NEAT1 [1285], FZD5 [1286], ARHGAP6 [1287], TGM2 [1288], LILRB4 [1289], FABP5 [1290], TIMP2 [1291], SH3RF3 [1292], CD63 [1293], CDKN2B [388], HIF1A [1294], SPHK1 [1295], VCAN (versican) [1296], PFKFB3 [1297], MFGE8 [1298], MYBPC3 [1299], DRAM1 [1300], CTSA (cathepsin A) [1301] and LILRB4 [1302] are observed in cardiac fibrosis. TP53 [1303], IL10 [1304], CXCR4 [1305], S100A8 [1306], S100A12 [1307], TIMP1 [1308], CES1 [1309], TFPI (tissue factor pathway inhibitor) [1310], IRS2 [1311], LPL (lipoprotein lipase) [1312], TRIB1 [1313], ABCA1 [1314] and IFITM3 [1185] positively correlate with hypercholesterolemia. PSMB10 [1315], IL10 [1316], SOCS3 [249], HP (haptoglobin) [1317], C1QC [1318], FKBP5 [1319], CXCR4 [1320], LMNA (lamin A/C) [1321], CREM (cAMP responsive element modulator) [622], NAMPT (nicotinamidephosphoribosyltransferase) [1322], TIMP1 [1323], F5 [1324], GAS6 [1325], S100A9 [1282], CES1 [1326], PF4 [1327], KCNE1 [1328], TREM1 [1329], NEAT1 [1330], PDE4D [1331], TIMP2 [1332], PLXND1 [1333], MFGE8 [1334], MYBPC3 [1335] and WDFY3 [1336] are the contributing factor to atrial fibrillation pathogenesis. These GO terms and the enrichment results of the REACTOME pathway suggest that the DEGs detected in the current investigation might be involvement in CA progression through the approaches described above.

A PPI network was constructed for the identified DEGs was defined by the degree, betweenness, stress and closeness rank. The most significant module was subsequently extracted from the PPI network. The results of PPI network and module analysis revealed that hub genes have a significant relationship with CA and its associated complications. Research has shown that CUL1 [488], HOXA1 [495], TP53 [491], LMNA (lamin A/C) [518], PDLIM7 [593] and HSPA1A [1337] plays an important role in the pathogenesis of atherosclerosis. HSPA8 [211], TP53 [217], LMNA (lamin A/C) [260], SNCA (synuclein alpha) [303], MRPL12 [200], FKBP5 [255] and HSPA1A [1338] are associated with the prognosis of neurologic complications. HSPA8 [405], TP53 [406], LMNA (lamin A/C) [423], FKBP5 [420] and HSPA1A [1339] have been and identified as a key genes in coronary artery disease. HSPA8 [663], TP53 [667] and FKBP5 [679] might be a potential target for cardiac hypertrophy treatment. HSPA8 [721], TP53 [725] and LMNA (lamin A/C) [747] are an important regulators of the hypertension. HSPA8 [1122], TP53 [725], HSPB1 [1165], LMNA (lamin A/C) [747], FKBP5 [1040] and HSPA1A [1340] might be involved in the development of diabetes mellitus. HSPA8 [663] and HSPA1A [1341] might serve as therapeutic targets for CA. The altered expression of HSPA8 [663], TP53 [600], LMNA (lamin A/C) [1265] and were observed to be associated with the progression of cardiac fibrosis. TP53 [104], HSPB1 [164], LMNA (lamin A/C) [132] and FKBP5 [128] have been shown to be associated with myocardial infarction. Altered levels of TP53 [104] and HSPA1A [1342] have been observed in heart failure patients. TP53 [600], LMNA (lamin A/C) [621], C1QBP [599] and FKBP5 [618] genetic risk factors has been shown to increase susceptibility to cardiomyopathy. A study showed that the genetic locus of TP53 [667], HSPB1 [920], LMNA (lamin A/C) [873], SNCA (synuclein alpha) [945], PDLIM7 [1006], FBL (fibrillarin) [803], C1QBP [818], FKBP5 [128], HSPA1A [1343], MAP2K1 [961], SLC7A5 [886] and ODC1 [979] were associated with susceptibility to inflammation. TP53 [725], HSPB1 [1073], LMNA (lamin A/C) [1045], FKBP5 [1040], HSPA1A [1344] and SLC7A5 [1052] have emerged as a potential target for obesity. TP53 [1209] and HSPA1A [1345] expression might be regarded as an indicator of susceptibility to myocardial ischemia. TP53 [1303] is a potential marker for the detection and prognosis of hypercholesterolemia. A previous study reported that LMNA (lamin A/C) [1321] and FKBP5 [1319] are altered expressed in atrial fibrillation. INCA1, ADAMTSL4, MRPS35, MRPS26, MRPL58, SLC16A3 and PPARG (peroxisome proliferator activated receptor gamma) might potentially be novel biomarkers or therapeutic targets for CA. The above results suggest that hub genes might be involved in CA and influence the progression of myocardial infarction, neurologic complications, heart failure, coronary artery disease, atherosclerosis, cardiomyopathy, cardiac hypertrophy, hypertension, inflammation, obesity, diabetes mellitus, myocardial ischemia, cardiac fibrosis, hypercholesterolemia and atrial fibrillation.

It is evident that miRNA-hub gene regulatory network and TF-hub gene regulatory network play a key part in the development of CA. In this way, it increases the knowledge of CA identification and contributes to targeted therapeutic management strategies and CA prediction. HSPB1 [1165] TP53 [725], ERBB2 [1116], PIN1 [1119], HSPA8 [1122], GRB10 [443], HSPA1A [1340], hsa-mir-1260a [1346], hsa-mir-101-3p [1347], GATA3 [1348] and PAX2 [1349] plays an important role in diabetes mellitus. Abnormal activation of HSPB1 [164], TP53 [104], IRF2 [1350], GATA3 [1351] and PRRX2 [1352] in myocardial infarction is a significant cause of disease progression. Some studies have shown that HSPB1 [920], TP53 [667], ERBB2 [802], PIN1 [806], C1QBP [818], ASPH (aspartate beta-hydroxylase) [539], PLSCR1 [982], MAP2K1 [961], BCL6 [942], SNCA (synuclein alpha) [945], HSPA1A [1343], HOXA5 [1353], GATA3 [1354] and SRF (serum response factor) [1355] plays a certain role in inflammation. A previous study reported that the HSPB1 [1073], TP53 [725], PIN1 [1012], BCL6 [1083], hsa-mir-369-3p [1356], HOXA5 [1357] and GATA3 [1348] biomarkers were associated with the obesity. The expression of TP53 [491], PIN1 [486], ASPH (aspartate beta-hydroxylase) [539], BCL6 [566], hsa-mir-140-5p [1358], HOXA5 [1359] and SRF (serum response factor) [1360] were altered in the heart following atherosclerosis. Altered expression of TP53 [217], MRPL12 [200], PIN1 [206], HSPA8 [211], PLSCR1 [318], BCL6 [302], SNCA (synuclein alpha) [303], HSPA1A [1337], hsa-mir-140-5p [1361], hsa-mir-101-3p [1362], hsa-mir-449b-5p [1363], HOXA5 [1364] and SRF (serum response factor) [1365] promotes the development of neurologic complications. TP53 [406], ERBB2 [401], PIN1 [404], HSPA8 [405], GRB10 [443], HSPA1A [1339], hsa-mir-1260a [1366], hsa-mir-369-3p [1367], GATA3 [1368] and SRF (serum response factor) [1369] are associated to the risk of coronary artery disease. TP53 [667], ERBB2 [659], PIN1 [661], HSPA8 [663], hsa-mir-140-5p [1370] and GATA3 [1351] are found to be associated with cardiac hypertrophy. Studies show that TP53 [725], PIN1 [718], HSPA8 [721], BCL6 [780], hsa-mir-140-5p [1371], HOXA5 [1372], NFYA (nuclear transcription factor Y subunit alpha) [1373] and SRF (serum response factor) [1374] are involved in the progression of hypertension. TP53 [600], PIN1 [1271], HSPA8 [663] and PRRX2 [1352] have been proved to be altered in cardiac fibrosis patients. Excessive activation of TP53 [104], ERBB2 [336], HSPA1A [1342], hsa-mir-101-3p [1375], NFYA (nuclear transcription factor Y subunit alpha) [1373] and SRF (serum response factor) [1376] have been observed in heart failure. Research has shown that TP53 [600] ERBB2 [595], C1QBP [599], hsa-mir-140-5p [1370], IRF2 [1377], GATA3 [1378] and SRF (serum response factor) [1379] participates in **cardiomyopathy.** Changes in TP53 [1209], ERBB2 [1207] and HSPA1A [1344] expression have been observed in myocardial ischemia. Previously, several studies have reported altered TP53 [1303] levels in patients with hypercholesterolemia. HSPA8 [663], HSPA1A [1341] and GATA3 [1380] expression were significantly altered in CA patients. Our results indicate that SH2D3C, LONRF1, hsa-mir-1914-5p, hsa-mir-5197-5p, hsa-mir-3664-3p, hsa-mir-598-3p, hsa-mir-3609, JUN (Jun proto-oncogene, AP-1 transcription factor subunit), USF2 and EN1 might play an important role in CA, providing new targets for treatment. Therefore, studying the biomarkers involved in the regulation of CA might be helpful to clarify the incidence or pathogenic mechanisms of CA.

## Conclusions

Based on the current research findings, it is feasible to use bioinformatics and NGS data analysis to screen and identify key biological markers related to CA and its associated complications includ myocardial infarction, neurologic complications, heart failure, coronary artery disease, atherosclerosis, cardiomyopathy, cardiac hypertrophy, hypertension, inflammation, obesity, diabetes mellitus, myocardial ischemia, cardiac fibrosis, hypercholesterolemia and atrial fibrillation. In the investigation of CA and and its associated complications, bioinformatics and NGS data analysis have been widely used and has made some important progress. Therefore, bioinformatics and NGS data analysis to screen and identify key biological markers related to CA and and its associated complications is a very promising research direction. Through in-depth bioinformatics and NGS data, we may discover more biological markers, providing more accurate means for early diagnosis, treatment, and prognostic assessment of these diseases.

## Acknowledgement

I thanks very much to Changde Cheng, St. Jude Children’s Research Hospital, Memphis, TN, USA, the author who deposited their NGS dataset GSE200117, into the public GEO database.

## Ethical approval

This article does not contain any studies with human participants or animals performed by any of the authors.

## Informed consent

No informed consent because this study does not contain human or animals participants.

## Availability of data and materials

The datasets supporting the conclusions of this article are available in the GEO (Gene Expression Omnibus) (https://www.ncbi.nlm.nih.gov/geo/) repository. [(GSE200117) https://www.ncbi.nlm.nih.gov/geo/query/acc.cgi?acc=GSE200117]

## Consent for publication

Not applicable.

## Author Contributions

B. V. - Writing original draft, and review and editing

C. V. - Software and investigation

